# GhostBuster: A Deep-Learning-based, Literature-Unbiased Gene Prioritization Tool for Gene Annotation Prediction

**DOI:** 10.1101/2025.06.22.660948

**Authors:** Giulio Deangeli, Maria Grazia Spillantini, Pietro Liò

## Abstract

All genes are not equal before literature. Despite the explosion of genomic data, a significant proportion of human protein-coding genes remain poorly characterized (“ghost genes”). Due to sociological dynamics in research, scientific literature disproportionately focuses on already well-annotated genes, reinforcing existing biases (bandwagon effect). This literature bias often permeates machine learning (ML) models trained on gene annotation tasks, leading to predictions that favor well-studied genes. Consequently, standard ML performance metrics may overestimate biological relevance by overfitting literature-derived patterns.

To address this challenge, we developed GhostBuster, an encoder-decoder ML platform designed to predict gene functions, disease associations and interactions while minimizing literature bias. We first compared the impact of biased (Gene Ontology) versus unbiased training datasets (LINCS, TCGA, STRING). While literature-biased sources yielded higher ML metrics, they also amplified bias by prioritizing well-characterized genes. In contrast, models trained on unbiased datasets were 2-3× more effective at identifying recently discovered gene annotations. Notably, one of the unbiased channels (TCGA), combined minimal amounts of literature bias with robust performance, at a test ROC-AUC of 0.8-0.95.

We demonstrate that GhostBuster can be applied to predict novel gene functions, refine pathway memberships, and prioritize intergenic GWAS hits. As the first ML framework explicitly designed to counteract literature bias, GhostBuster offers a powerful tool for uncovering the roles of understudied genes in cellular function, disease, and molecular networks.

**Graphical Abstract:** 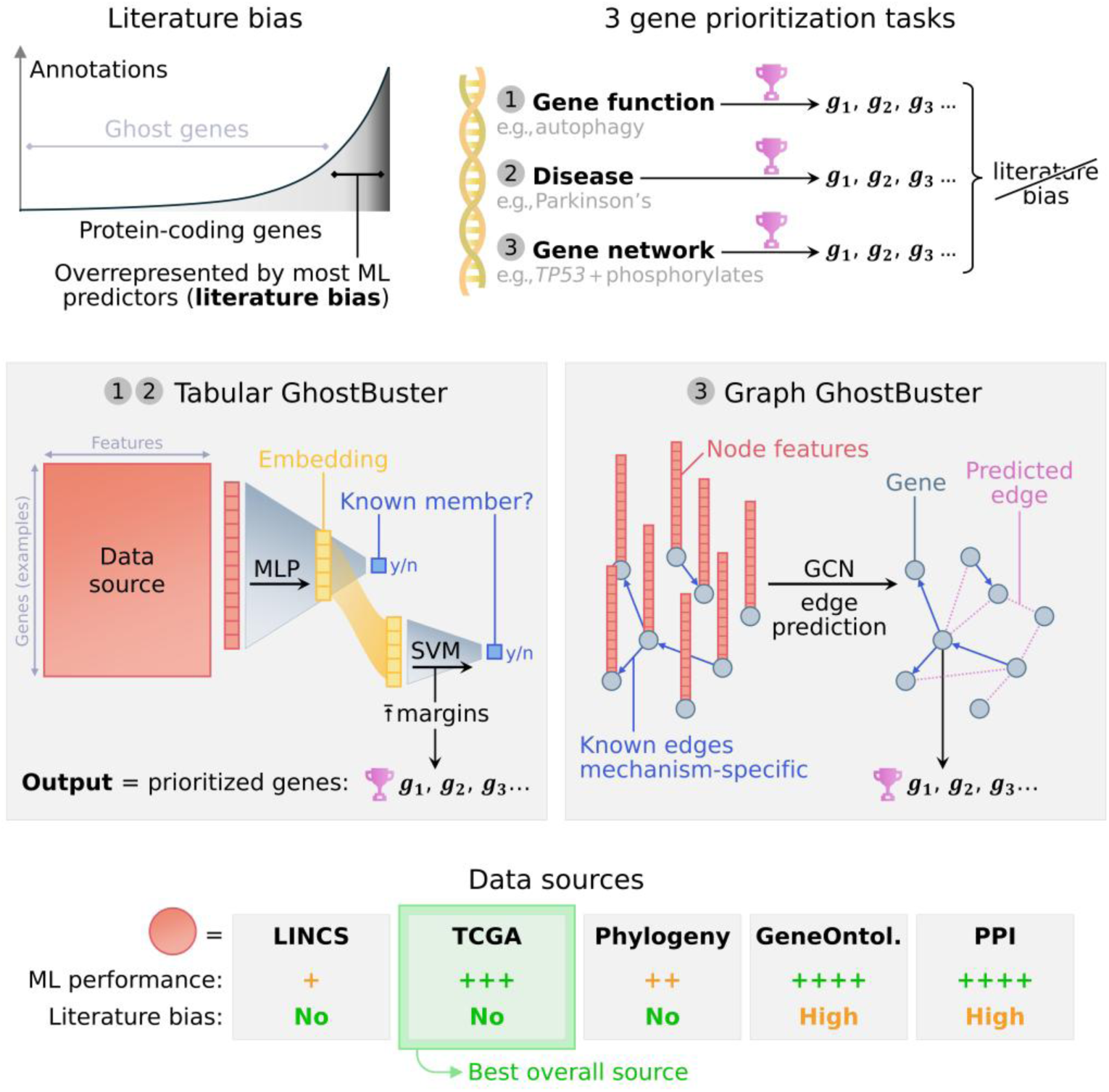

GCN: graph convolutional neural network; GeneOntol.: Gene Ontology; LINCS: Library of Integrated Network-Based Cellular Signatures; MLP: multi-layer perceptron (deep learning); PPI: (physical) protein-protein interactions; SVM: support vector machine; TCGA: The Cancer Genome Atlas; y/n: yes or no (binary classification).

## Introduction

Our mechanistic understanding of biology and disease is constrained by limited knowledge of genes’ functions and interplay. This process involves three major steps: sequencing the genome, identifying the genes, and functionally characterizing them. Thanks to advancements in omic techniques, major progress has been made on the former two steps: genomics has enabled a complete sequencing of the human genome in 2022,^1^ and the sequencing of about 500,000 other species.^2^ In parallel, next-generation RNA sequencing and modern proteomics can experimentally detect RNA-transcribed genes or protein-coding genes, scalably across tissues and species. For example, we officially recognize about 20,000 protein-coding and 20-40,000 non-coding genes in the human species (Table 1).

**Table 1.** Human genes in the NCBI Gene dataset and the EBI Ensembl dataset. Genes are defined as the genome portions that are transcribed into RNAs, while protein-coding genes are defined as the genome portions that are transcribed into RNAs and translated into proteins. Before the advent of next-generation RNA sequencing and modern proteomics, gene identification was a highly contentious matter. Initial estimates for the human protein-coding genes ranged between 50,000 and >140,000, while the official figure from the Human Genome Project was of 26,588.^60^ With the refinement of omic techniques over the years, the number eventually settled to the currently accepted value of about 20,000 protein-coding genes in the human species, and about 5 times as many transcript products due to alternative splicing, and alternative transcription initiation or termination sites).^61^ While the protein-coding genes account for 2% of the genome, most of the remainder 75-90% “junk” DNA is pervasively transcribed as non-coding RNA genes (ncRNAs), which include, among others: ribosomal RNAs (rRNAs), transfer RNAs (tRNAs), microRNAs (miRNAs), long non-coding RNAs (lncRNAs), small nuclear RNA (snRNA) and small nucleolar RNA (snoRNA).^62^ Interestingly, recent evidence from ribosome profiling and mass spectrometry demonstrated that several ncRNAs can actually bind to the ribosome and be translated into proteins. This unexpected turn makes the coding/non-coding dichotomy blurrier than expected in the classical paradigm: this “hidden” proteome encoded by ncRNAs is an active area of research, still in its infancy.^62–64^ (a) Number of genes recognized by the National Center for Biotechnology Information (NCBI) Gene^65^ dataset, subdivided into different categories, as of January 1, 2025. (b) Number of genes recognized by the European Bioinformatics Institute (EBI) Ensembl^66^ dataset, subdivided into different categories, as of January 1, 2025.

|  |  |  |  |
| --- | --- | --- | --- |
| **a** |  | **b** |  |
| **NCBI Gene dataset** |  | **EBI Ensembl dataset** |  |
| Coding genes | **20,621** | Coding genes | **20,527** (of which 659 readthrough) |
| Non-coding genes | **24,974** | ↳ gene transcripts | **387,944** |
| ↳ rRNA | 785 | Non-coding genes | **42,160** |
| ↳ snRNA | 166 | ↳ small non-coding | 4,867 |
| ↳ snoRNA | 1,201 | ↳ long non-coding | 35,376 (of which 300 readthrough) |
| ↳ tRNA | 701 | ↳ other non-coding | 2,217 |
| ↳ other ncRNA | 22,121 | Pseudogenes | **15,207** (of which 1 readthrough) |
| Pseudogenes | **17,483** |  |  |
| Biological region | **12,8261** |  |  |
| Other | 845 |  |  |
| Unknown | 1,189 |  |  |

Whereas gene sequencing has grown exponentially since the late ‘90s, gene functional characterization has grown only linearly, as it still relies on gathering large amounts of non-scalable experimental investigations.^2,3^ Large biocurated repositories have been set up, harmonizing annotations in a controlled vocabulary.^4,5^ We may classify them in three domains:

- **Gene functions/properties**, such as: molecular functions (GO^6^, InterPro^7^; Pfam^8^; UniProt^9^); biological processes (FunCat^10^; GO^6^); high/low expression in a certain tissue (Human Protein Atlas^11^); or cellular localizations (GO^6^, COMPARTMENTS,^12^ LOCATE,^13^ SynGo^14^).
- **Disease involvement**, such as: contributing to disease mechanism according to experimental literature (DisGeNET^15^; KEGG^16^; HuGE^17^; MSigDB^18^; RGD^19^); being genetically involved based on Mendelian inheritance (OMIM^20^); or being a candidate hit based on genetic association studies (PheGenI^21^, GAD^22^).
- **Gene regulatory networks**, such as: protein-protein interaction (BioGRID^23^; I2D^24^; IntAct^25^; iRefIndex^26^; MiNT^27^; MIPS^28^); phylogenetic similarity (STRING^29^); catalyzing successive reactions in a pathway (BioCarta^30^; ConsensusPathDB^31^; hiPathDB^32^; HIPPIE^33^; KEGG^16^; MSigDB^18^; Reactome^34^; RGD^19^; Wikipathways^35^); or functional annotations such as up/downregulation and biochemical modifications (KEGG^16^; Signor^36^).

A number of bioinformatic tools have been developed to predict gene annotation *in silico* across all three of these domains, namely gene function prediction (GFP, Table 2), gene disease prediction (GDP, Table 3) and gene network prediction (GNP, Table 4). They generally train their models based on gene annotations from the “highly annotated” organisms like *H. sapiens*. However, a major problem has been often overlooked: even the annotations in such model organisms are “literature-biased”, namely (i) dramatically incomplete, (ii) highly biased in favor of a subset of genes, and (iii) sometimes even erroneous.

i. Let’s start from incompleteness, taking the human species as an example. The number of annotations (“richness”) available for each gene is extremely imbalanced. Reportedly, 1/3 protein-coding genes have essentially no literature or known function, and are colloquially known as “ghost” genes^37,38^; most research is concentrated on 2,000 of the ∼20,000 human genes; and among those, a very small gene subset have dramatically more coverage than anyone else.^3^ Indeed, richness fits an exponential curve, and so do genes’ references in PubMed abstracts, PubMed titles and research funding grants.^39^ By our own measurement, the gene’s Gini coefficient with respect to PubMed papers stands at 0.67 for protein-coding genes, and at 0.90 across all genes. Even if we look at highly expressed genes, the picture does not improve much. In 2014, Pandey *et al.* coined the term “ignorome”, defined as the “set of genes with intense *and* selective expression in specific tissues […] uncharacterized with respect to biological function”. They showed that, among the 648 genes that are highly and specifically expressed in the brain: 5% genes absorbed 70% of neuroscience literature; 30% genes absorbed only 0.1% of literature; and 20% genes had essentially no literature at all.^37^ One could argue that, perhaps, experimentalists *started* from characterizing a limited group of genes, and science coverage is progressively focusing on filling the gaps. Strikingly, the opposite is true. The rich genes only become richer over time (Matthew effect).^3,40^ For instance, the Gini coefficient of Gene Ontology^41^ went up from 0.25 in 2001, to 0.47 in 2017, indicating that rich genes continued to receive even more annotations over time.^42^
ii. When investigators studied which factors explain such a dramatic difference in literature coverage, they determined that genes’ popularity does NOT correlate with their amount of differential expression in disease (Spearman *ρ* = –0.003, *p* = 0.836)^42^; their genetic association with disease (*ρ* = 0.017, *p* = 0.836)^42^; their connectivity in coexpression networks^37^; their phylogenetic conservation in terms of ortholog (Pearson *r* = – 0.085)^37^ or paralog abundance (*r* = 0.16).^37^ Instead, it correlates significantly with several bias factors, which have nothing to do with their intrinsic biology: things like their annotation richness (*ρ* = 0.110, *p* = 2.1 · 10^−16^)^42^; the year of their first publication (*ρ* = –0.58); the number of publications in the previous decade (*ρ* = 0.84)^3^; NIH funding allocation (Spearman *ρ* = 0.95)^3^; and some physical and chemical features, such as gene length, hydrophobicity and expression level in certain organs.^3^ Overall, the best predictor of scientific interest is exactly the bandwagon effect: research focuses on the genes that we already know well, likely as a result of research social structures and funding opportunities.^37^ Importance is a self-fulfilling prophecy. This tendency cannot be in the best interest of science: notably, some of today’s strongest disease-related genes were ghost genes when their disease involvement was first described, such as *BRCA1*,^43^ *SIRT1*,^44^ *SNCA*^45^ and *TARDP*.^46^
iii. Annotation reportedly suffers from reliability issues, being sourced from scientific publications. Reproducibility studies have consistently indicated that the majority of claims reported in literature are false.^47^ For example, a survey showed that only 1 in 4 pre-clinical papers are reproducible.^48^ This crisis has been attributed for the most part to research social structures.^47,48^

**Table 2.**
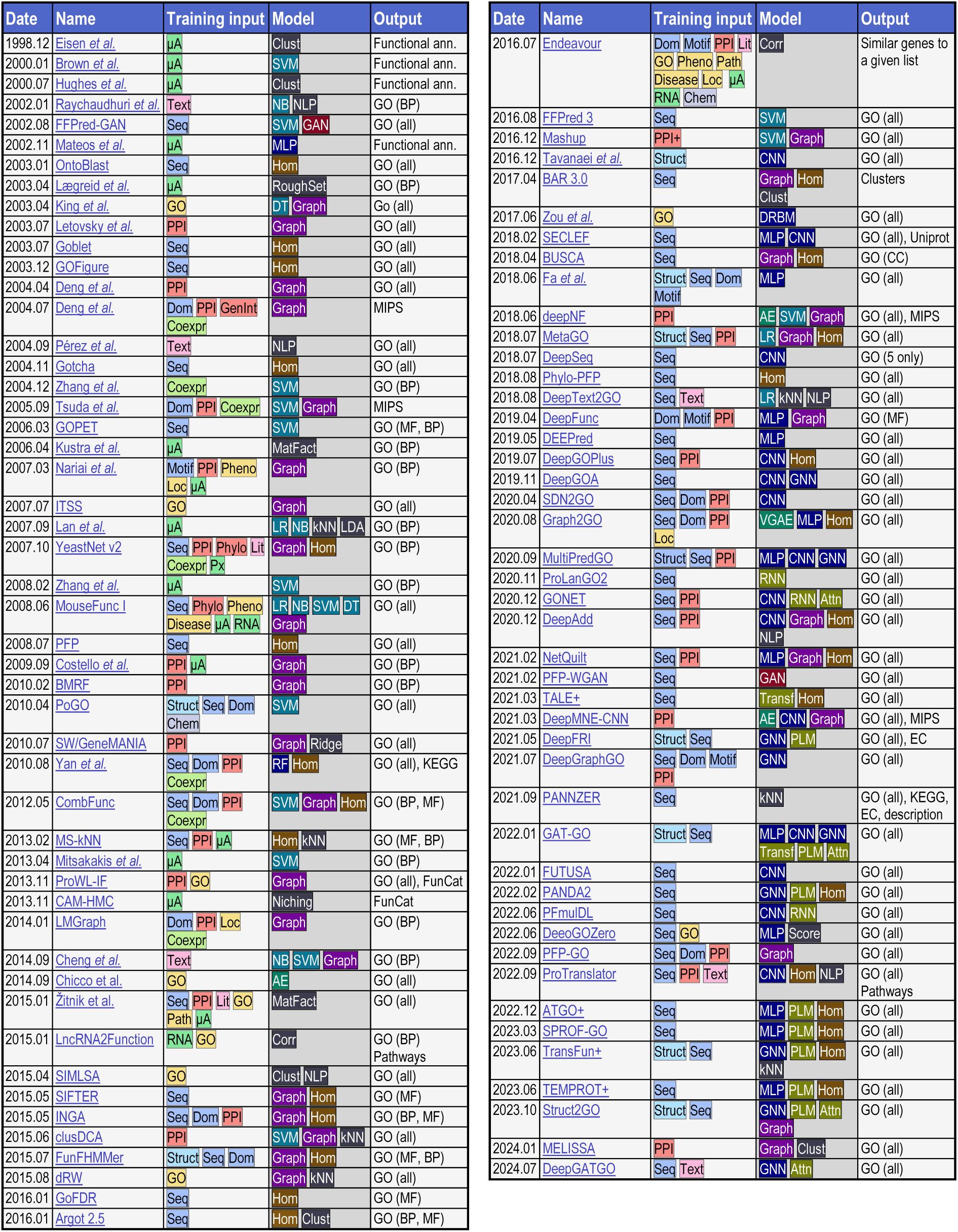
Literature overview of Gene Function Prediction (GFP), examining the training input data, architecture and prediction output. The table offers a bird’s eye overview of the last 25 years of gene function prediction (GFP) models. While it does not have any ambition of completeness—as it would be beyond the scope of the present research—it makes it easy to spot general patterns in researchers’ preference along the years, in terms of both input data and model choice. Papers are sorted chronologically by their publication date. AE: autoencoder; Attn: attention; BP: Gene Ontology biological process; CC: Gene Ontology cellular compartment; Chem: biochemical information; Clust: clustering; CNN: convolutional neural network; Coexpr: coexpression; Disease: information comparing multiple diseases; Dom: protein domains; DRBM: restricted Boltzmann machine; DT: decision tree; GAN: generative adversarial network; GNN: graph neural network; GO: Gene Ontology (or analogous repositories); Graph: graph-based scoring system (non-machine learning); Hom: homology search; *k* NN: *k* -nearest neighbors; Lit: literature-based metrics; Loc: cell localization information; LR: linear regression; MatFact: matrix factorisation; MF: Gene Ontology molecular function; MLP: multi-layer perceptron; Motif: protein motifs; NB: naïve Bayes models; NLP: natural language processing; Path: pathway membership; Pheno: phenotypic information; Phylo: phylogenetic information; PLM: protein language model; PPI: protein-protein interactions; in the broadest sense; Px: proteomics; ridge; ridge regression; RF: random forest; RNA: bulk RNA sequencing; RNN: recurrent neural network; Score: algorithmic scoring system (non-machine learning); Seq: sequence; Struct: structural information; SVM: support vector machine; Trans: transformer; VGAE: variational graph autoencoder; μA: DNA microarrays.

**Table 3.**
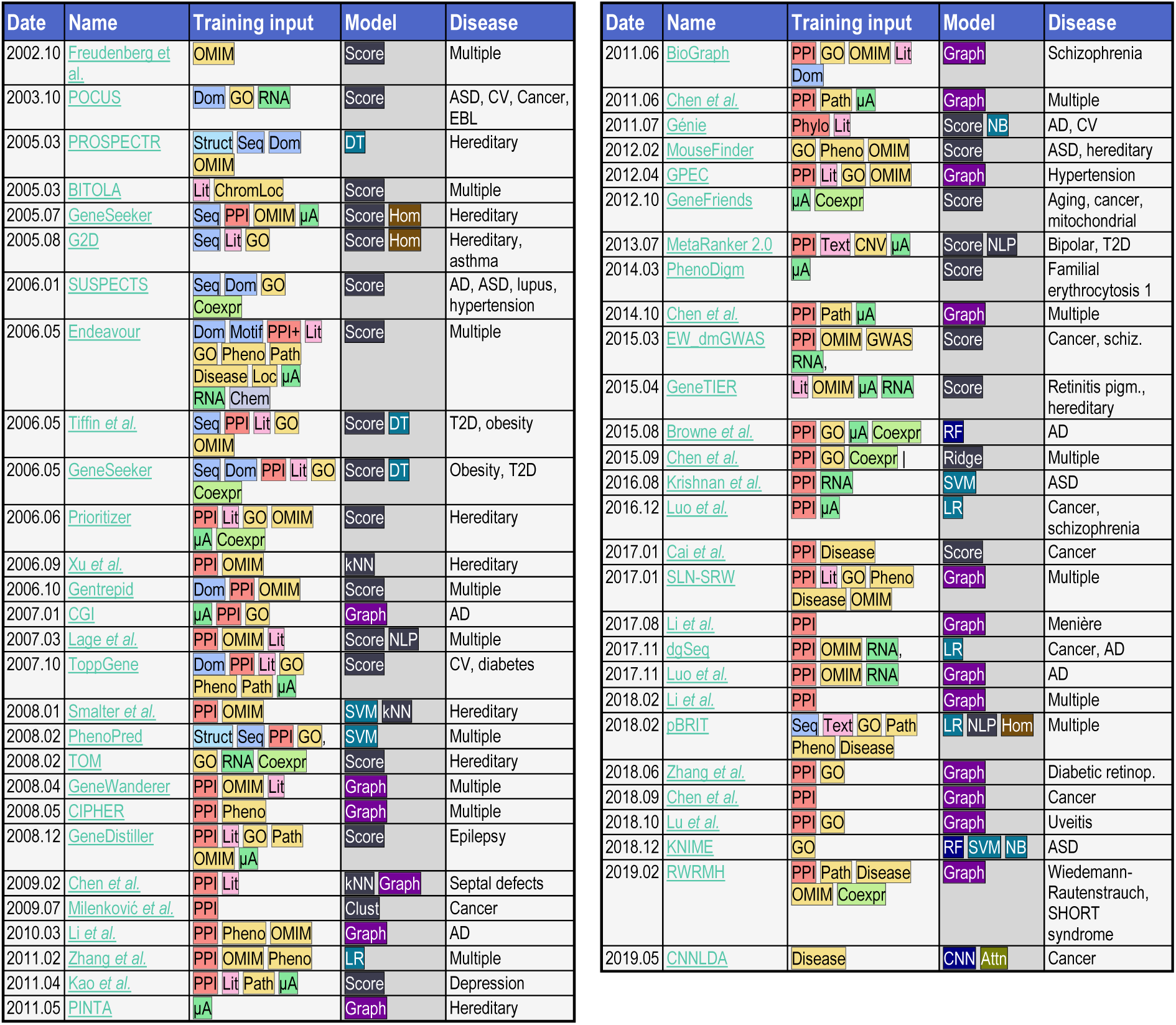
Literature overview of Gene Disease Prediction (GDP), examining the training input data, architecture and prediction output. The table offers a bird’s eye overview of the last 25 years of gene disease prediction (GDP) models. While it does not have any ambition of completeness—as it would be beyond the scope of the present research—it makes it easy to spot general patterns in researchers’ preference along the years, in terms of both input data and model choice. Papers are sorted chronologically by their publication date. AD: Alzheimer’s disease; ASD: autism spectrum disorder; Clust: clustering; CNN: convolutional neural network; Coexpr: coexpression; CV: cardiovascular diseases; diabetic retinop.: diabetic retinopathy; Disease: information comparing multiple diseases; Dom: protein domains; DT: decision tree; EBL: epidermolysis bullosa letalis; GO: Gene Ontology (or analogous repositories); Graph: graph-based scoring system (non-machine learning); Hom: homology search; *k*NN: *k*-nearest neighbors; Lit: literature-based metrics; LR: linear regression; Motif: protein motifs; NB: naïve Bayes models; NLP: natural language processing; OMIM: Online Mendelian Inheritance in Man (or analogous repositories); Path: pathway membership; Pheno: phenotypic information; Phylo: phylogenetic information; PPI: protein-protein interactions, in the broadest sense; Px: proteomics; retinitis pigm.: retinitis pigmentosa; RF: random forest; RNA: bulk RNA sequencing; Score: algorithmic scoring system (non-machine learning); Seq: sequence; Struct: structural information; SVM: support vector machine; T2D: type 2 diabetes; μA: DNA microarrays.

**Table 4.** Literature overview of gene network prediction (GNP), examining the training input data, architecture and prediction output. The table offers a bird’s eye overview of the last 10 years of gene network prediction (GNP) models. While it does not have any ambition of completeness—as it would be beyond the scope of the present research—it makes it easy to spot general patterns in researchers’ preference along the years, in terms of both input data and model choice. Papers are sorted chronologically by their publication date. Of note, modern machine learning has started to be applied to GNP from the mid 2010’s. Previous approaches were generally rule-based, and are mostly out of scope for our analysis. For a review of such legacy methods, please consult Madhukar *et al.* (2015),^67^ and Delgado & Gómez-Vela (2019).^68^ ChIP-seq: chromatin immunoprecipitation followed by sequencing; CNN: convolutional neural network; GAT: graph attention network; GCN: graph convolutional neural network; GOF: gain of function; KEGG: Kyoto Encyclopedia of Genes and Genomes; LOF: loss of function; LR: linear regression; MLP: multi-layer perceptron; ODE: ordinary differential equation; PID: partial information decomposition; PPI: protein-protein interaction; RF: random forest; RNA: bulk RNA sequencing; Score: algorithmic scoring system (non-machine learning); scRNA: single-cell RNA sequencing; TF: transcription factor; μA: DNA microarrays.

| Date | Name | Training input | Model | Output |
| --- | --- | --- | --- | --- |
| 2010.09 | GENIE3 | Network | RF | TF-target (RegulonDB) |
| 2017.08 | SCODE | scRNA | LR\|ODE | TF-target (DNase footprint, TF-binding motifs) |
| 2017.09 | SINCERITIES | scRNA | Score | Gene interactions (any kind) |
| 2019.04 | GNE | μA | MLP | PPI (Biogrid) |
| 2019.07 | CNNC | scRNA RNA | GCN | TF-target (ChIP-seq), activation/inhibition (KEGG; Reactome) |
| 2019.09 | PIDC | scRNA | PID | Gene interactions (any kind) |
| 2021.11 | DeepDRIM | scRNA | CNN | TF-target (ChIP-seq) |
| 2022.09 | GRN-transformer | scRNA | GAT\|RF | TF-target (ChIP-seq), PPI (STRING) |
| 2022.10 | GENELink | scRNA | GAT | TF-target (ChIP-seq), PPI (STRING), LOF/GOF |
| 2023.11 | GNNLink | scRNA | GCN | TF-target (ChIP-seq), PPI (STRING), LOF/GOF |

All this is relevant because, if one trains a GFP/GDP/GNP model based on literature-biased data, it is plausible that the predictions themselves may reflect literature bias, for example, favoring genes that have already been extensively studied as opposed to ghost genes. Conversely, if one trains a model on literature-agnostic data, performance may be numerically lower by machine learning (ML) metrics, but the absence of bias may make results more biologically “interesting”, i.e., more in line with the biological truth. This question has never been systematically studied, as most recent papers quantify their models’ performance merely based on ML scores, such as the ROC-AUC, while their training data were literature-biased in (almost) all the cases (see Table 2 and Table 3).

As a prime example, a majority of the recent models were trained on protein-protein interaction (PPI) data, under a probabilistic paradigm known as “guilt by association” (GBA, also known as linkage-based assumption)^49^: essentially, the annotation of a new protein likely reflects that of its PPI partners.^50^ For example, if a protein interacts with many cytoskeletal proteins, chances are it is itself a cytoskeletal protein. This assumption is biologically questionable, for multiple reasons. First, it was shown that the GBA-relevant PPI edges (i.e., edges which cause functionally related genes to be more interconnected) represent less than 0.01%, among millions of other irrelevant connections.^51^ Next, not all cell functions are mediated by physical interaction (second messengers, DNA binding, RNA interference, microRNA activity, pH regulation, membrane permeability…). Next, most reaction-relevant interactions happen at low affinity, as they are meant to be transient, implying they may not be detected by PPIs.^50,52^ Next, PPIs are obtained using a bait-prey system, where baits are chosen by investigators in the context of a specific experiment, causing them to be literature-biased, laboratory-biased,^53^ highly incomplete^53,54^ and inconsistent.^55^ Finally, and most importantly, today’s PPIs are a highly ambiguous category. Originally, PPIs started as experimentally derived physical binding partners (interactomics).^56^ Over the years, their definition has been expanded to a variety of “non-physical” PPIs, which are actually correlative in nature: thing like, sharing a similar expression pattern (coexpression interactions),^57^ sharing perturbation effects (genetic interactions),^58^ sharing orthology/paralogy patterns (phylogenetic interaction)^29^ or sharing similarities in literature coverage.^59^ Most of the GFP/GDP/GNP implementations have lost sight of the biology as they often feed large, catch-all PPI matrices into their model, without distinguishing between different interaction modalities (see Table 4).^29^

To overcome literature bias and incomplete biological granularity of existing GFP/GDP/GNP architectures, we introduce GHOSTBUSTER, a tool that combines state-of-the art deep learning with literature-unbiased data sources to predict novel annotations, with particular focus on ghost genes. We created two versions GhostBuster: (i) Tabular GhostBuster, which targets a provided lists of genes that are known to be involved in a given cell function or disease; it creates an implicit rule of what factors are shared among those lister genes, and prioritizes the other non-lister genes based on how closely they match such rule, for GFP/GDP purposes. Likewise, (ii) a Graph GhostBuster, which targets a provided list of gene pairs that interact in a given biological modality (say, phosphorylation), creates an implicit rule, and prioritizes the other non-lister gene pairs, for GNP purposes. As for the input information, we compared GhostBuster’s performance using five different input channels: three are literature-agnostic as they are experimentally derived, whereas two are literature-biased. We showed that, from a ML metric perspective, literature-biased channels generally outperform the unbiased ones. However, confounder analysis reveals that predictions from unbiased channels are more novel, unbiased, and biologically interesting. This suggests that ML metrics are not a reliable scoring system for GFP/GDP/GNP tasks, as they reward models for learning literature bias.

To the best of our knowledge, GhostBuster is the first modern ML architecture that makes GFP/GDP predictions in a completely literature-unbiased way, and the first GNP architecture that achieves biological mechanistic resolution. These features make it an ideal tool to study a ghost gene’s involvement in a given biological process or disease.

## Results

### Target dataset preparation

In Tabular GhostBuster, the target (***Y***) is a list of genes that are currently known to be involved in a specific “phenotype” of interest (a cell function, cell compartment or disease, say, “autophagy”); the prediction output is a prioritized gene list, based on how closely they match the common factors shared by all positive genes in the target list—i.e., a genome-wide ranking of genes, based on how likely they are to be involved in, say, autophagy. For the sake of validating Tabular GhostBuster, we repeated the training against 81 phenotypes (Supplementary Table 1), selected with the intent to cover biological fields of great scientific relevance. These 81 phenotypes include: 50 cell functions and 4 cell localization from Gene Ontology (GO) dataset^41^; 21 lists of genes involved in disease, taken from the Molecular Signatures Database (MSigDB)^18^; and 6 lists of gene hits from genome-wide association studies (GWAS), sourced from the Phenotype-Genotype Integrator (PheGenI).^21^ Among the 81 target classes, we included three clusters that represent sub-mechanisms of the same biological processes, namely 8 classes of DNA damage response, 6 classes of autophagy and 9 classes of metabolism-related functions. A gene-centric t-distributed stochastic neighbor embedding (t-SNE) plot of the 81 gene classes confirmed that they sample the entire “space” of cell biology in a fairly uniform way (Supplementary Figure 1).

Conversely, in Graph GhostBuster, the target list is represented by a set of gene-gene pairs that interact by a specific biological modality (e.g., phosphorylation); the prediction output is a prioritized gene pair list, based on how closely they match the common factors that are shared by all positive gene pairs in the target list. For the sake of validating Graph GhostBuster, we repeated the training against 21 interaction types (Supplementary Table 2), sourced from our own Biology Mega Graph. The latter is a harmonized knowledge graph encompassing 322 million known gene-gene relations with full biological granularity, and it is part of our VLab ML library.^69^ It was created by harmonizing together 11 biological repositories, namely: ComplexPortal,^70^ human DEPhOsphorylation Database (DEPOD),^71^ Gene Ontology,^6^ MicroT (miRBase and MirGeneDB versions),^72^ miRBase,^73^ NCBI Gene,^74^ PhosphoSitePlus,^75^ Signor,^36^ STRING^76^ and TarBase.^77^ Harmonization was conducted using a well-defined vocabulary of 69 edge labels, which express the biological effect associated with the edge (e.g. “gene *A* `phosphorylates` gene *B*”); of these 69 levels, the top 21 ones with the highest number of annotations (≥460) were utilized as our training sets.

### Training dataset preparation

Moving on to the training dataset (***X***), that is, the information that was used to represent the genes in the ML model, all GhostBuster experiments were independently trained using one of four information channels.

i. The Library of Integrated Network-Based Cellular Signatures (LINCS, 2020 version), is a collection of over 3 million gene expression experiments.^78,79^ Cell cultures were assessed in different genetic perturbation and treatment conditions, which include compounds, ligands, loss of function by shRNA, loss of function by CRISPR and overexpression by cDNA. Replicates were aggregated into a signature, expressing the differential vector of the treated compared to the untreated group.^80^ After pre-processing and filtering, the dataset comprised 12,325 genes (examples) by 282,990 biological signatures (features), encompassing 184 cell lines.
ii. The Cancer Genome Atlas (TCGA) is a collection of harmonized multiomic experiments from thousands of cancerous and healthy human tissue biopsies.^81^ We aggregated all available bulk RNA sequencing datasets (as of September 2024). After pre-processing, resulting dataset comprised 36,027 detected genes (examples) by 24,196 biological samples (features), encompassing 34 human tissues.
iii. Search Tool for the Retrieval of Interacting Genes/Proteins (STRING) is a general-purpose protein-protein interaction database, encompassing 7 interaction modes.^59,76^ We excluded all literature-biased interaction modes, namely `experimental`, `database` and `textmining`. We only included the phylogeny-based modes, namely `coexpression`, `neighborhood`, `cooccurrence` and `fusion`: the former one measures the expression correlation across a large panel of conditions and tissues; the latter three are derived from applying a gene-recognition algorithm across the phylogenetic tree, detecting patterns between gene pairs, that reportedly correlate with the likelihood of the two genes sharing similar functions.^82^ After pre-processing and filtering, the dataset comprised 18,845 interactor genes (examples) by 49,402 partner genes × interaction modalities (features).
iv. Gene Ontology (GO) is a collection of molecular functions, biological processes and cellular compartments, and their respective member genes. We converted the membership lists into a gene-class adjacency matrix. After pre-processing, the dataset comprised 19,237 genes (examples) by 17,873 GO classes (features). Of these channels, LINCS, TCGA and STRING are literature-agnostic, while GO is literature-based, thus, biased. Tabular GhostBuster also included two additional control groups, namely the PPI channel and the GPT channel: differently from the other four, these data sources were not utilized to train a ML model, rather, they relied on a scoring metric.
v. The PPI channel embodies the classic GBA approach: it quantifies a gene’s physical interactions with the target list member genes. Physical interactions were sourced from the `experiments` section of STRING.^76^ Genes were then ranked based on this metric, in line with the GBA assumption that, when a gene’s interactors are marked with a given function, the gene itself has increased chances of sharing such function.
vi. OpenAI Generative Pre-trained Transformer (GPT): the OpenAI GPT-4o mini^83^ API was utilized to quantify, by a standardized score from 0 to 10 (see Supplementary Table 3), how strongly each gene is currently known in literature for being involved in the phenotype of interest. The scoring sets a “satisfactory” threshold at 6, that is, values equal or smaller than 5 indicate no robust association with the phenotype has been found, so far; values of 6 or above indicate that at least some studies have called the gene to participate in the phenotype.

### Tabular GhostBuster literature bias in the input is reflected by the literature bias in the prediction

Tabular GhostBuster was trained in a 10-fold stratified cross-validation setting, independently for each of the 81 phenotypes (Supplementary Table 1), to allow an equal proportion of positive examples in each CV epoch. For each phenotype, the model produced a genome-wide prioritization list for each of the 4 input channels (plus the PPI scoring algorithm). ML metrics were measured in the test dataset prediction, with respect to the original target list (Figure 1, Supplementary Figure 2). The unbiased channels achieved a ROC-AUC of 0.68 (LINCS), 0.80 (TCGA) and 0.81 (STRING); conversely, the biased channels yielded 0.88 (GO) and 0.86 (PPI). Overall, the biased channels outperformed the unbiased ones. These scores are roughly comparable to the current gold standard predictors for biological processes (BPs), showing ROC-AUCs of 0.6-0.8 for BPs^84,85^; in 2023, Struct2GO established a new record at 0.873.^86^

**Figure 1.**
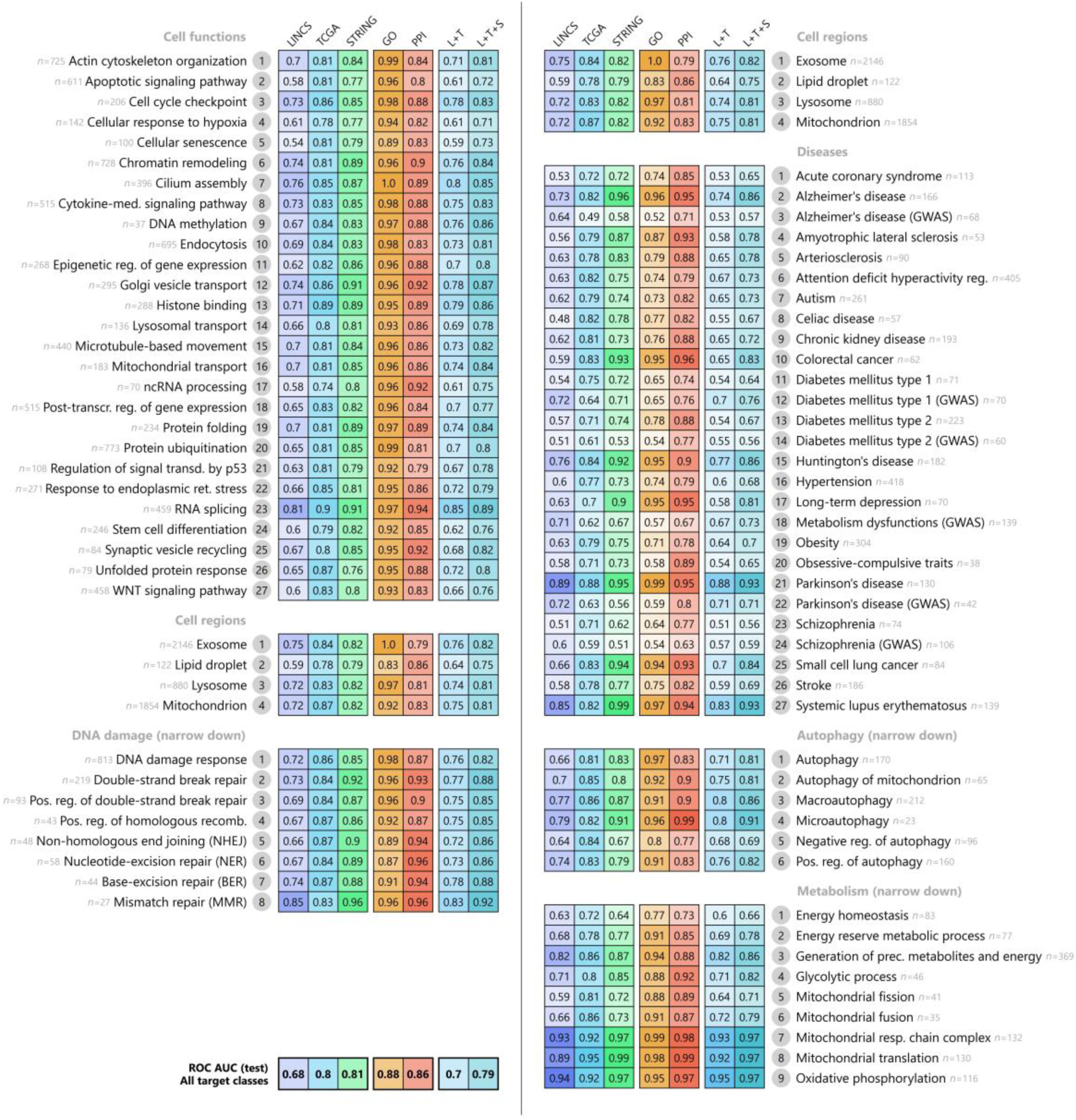
Tabular GhostBuster performance on the 81 target lists, measured by ROC-AUC, in the test dataset prediction. The plot measures GhostBuster’s performance in terms of the receiver operating characteristic area under the curve (ROC-AUC), in the test dataset prediction. Column by column, first, it displays GhostBuster’s performance when trained on the three “unbiased” data sources, namely LINCS (blue), TCGA (cyan) and STRING (green). Next, the performance when trained on the “biased” data source, namely GO (orange). Next, the performance of a simple “guilt by association”-type gene prioritization script, based on the physical protein-protein interactions in STRING (red). Next, the combined performance obtained by averaging out the predictions from LINCS and TCGA (L+T), and from LINCS, TCGA and STRING (L+T+S). The phenotype labels are displayed next to each line, accompanied by the numerosity of the corresponding target gene lists (*n*). The average performance is displayed in the bottom left corner. All cells are color-coded based on the performance score, in a non-linear fashion. Overall, GO achieved the best performance, followed by PPI, STRING, TCGA and finally LINCS. This distribution accurately reflects the amount of literature bias that is present in each of the 5 sources.

These results confirm our initial concern: performance correlates with the amount of literature bias in the data source. Of the five channels, the best performer was GO, which is arguably the “least biological” of the five, as it “predicts literature based on literature itself”. For example, on Parkinson’s and Alzheimer’s MSigDB lists, the GO channel yielded a ROC-AUC of 0.99 and 0.96 respectively, in the test dataset: such extreme accuracy is beyond the biologically plausible at our current (highly incomplete) level of mechanistic understanding of both diseases.^87^ We can conclude that performance becomes misleadingly more gratifying, the more literature-biased our source is, in a “short-circuit”, self-fulfilling-prophecy.

Literature bias cannot be eliminated completely from the output, as it inherently affects the target vector ***Y***. After all, it would be impossible to even *represent* concepts like autophagy or type 1 diabetes, without relying on what we know about such functions or diseases, however biased and imperfect. However, it can be eliminated from the training dataset ***X***. Therefore, we explored whether utilizing an unbiased training dataset ***X*** may “pay off” by reducing literature bias in the predicted output, ***Y***′ . To address this question, we assessed seven confounders that reportedly play a role in literature bias,^3,37,42^ sampling their average values in the top-ranked, non-target genes for each of the data channels. These confounders are, for each gene: (i) the number of annotations reported in GO and MSigDB (“richness”); (ii) the coverage in PubMed articles; (iii) the average log-transformed transcriptomic expression across all RNA-sequenced tissues in TCGA; (iv) the percentage of ghost genes, defined as genes in the lower two thirds of the richness distribution; (v) the percentage of ghost genes, defined as the genes in the lower two thirds of the PubMed coverage distribution; (vi) the number of protein-protein interactions with the target gene members; (vii) the average GPT score. We performed the measurement on the *n* top-ranked output genes, at varying *n* thresholds, starting from the complete gene space (no threshold), followed by the top 5,000 in GhostBusters’s output, moving progressively up to the top 10 in GhostBusters’s output (Figure 2).

**Figure 2.**
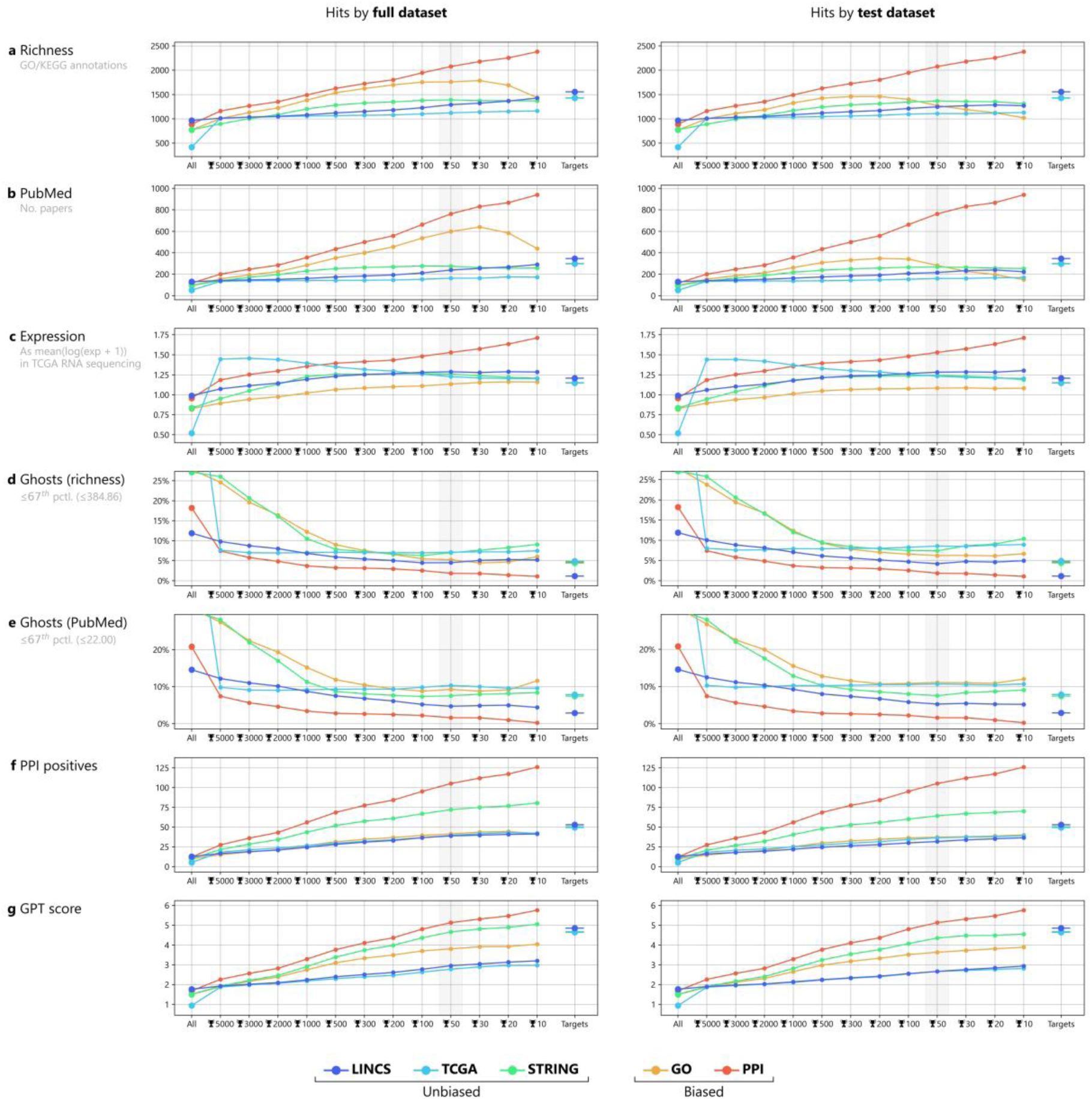
Literature bias confounders in the non-target top-ranked genes by Tabular GhostBuster. The plot quantifies 7 literature bias confounders in the prediction of Tabular GhostBuster across all 81 target lists, with respect to the data channel that was provided in their input. Confounders (y-axis) were evaluated in the output test dataset prediction from each data channel (after excluding the positive genes), by sampling the top-ranked genes in the output at different threshold levels (x-axis). Thresholding starts from the whole dataset (“All”); next, it continues with Tabular GhostBuster’s top *n* genes (🏆 *n*), starting from 5,000 and moving up to the top 10; finally, on the right side, the average values for the positive genes (targets) are shown for comparison. The analysis was repeated for all data channels: Tabular GhostBuster trained on the three “unbiased” data sources, namely LINCS (blue), TCGA (cyan) and STRING (green); next, Tabular GhostBuster trained on the “biased” data source, namely GO (orange); next, a simple GBA-type gene prioritization script, based on the physical protein-protein interactions in STRING (red). The explored confounders are, for each predicted gene: (a) the number of annotations (“richness”) reported in Gene Ontology (GO) and the Molecular Signatures Database (MSigDB); (b) the coverage in PubMed articles; (c) the average log-transformed transcriptomic expression across all tissues in TCGA; (d) the percentage of ghost genes, defined as genes in the lower two thirds of the richness distribution; (e) the percentage of ghost genes, defined as genes in the lower two thirds of the PubMed coverage distribution; (f) the number of protein-protein interactions with the target gene members; (g) the average GPT score of the predicted genes. Overall, predictions from the biased sources generally yield a higher bias across all the explored confounders, namely: higher richness, higher PubMed score, higher average expression, lower percentage of ghost genes, higher PPI score and higher GPT score. Of note, the LINCS channel also output a small minority of ghost genes, however, this is because very few ghost genes are present in the LINCS namespace in the first place, as LINCS only covers 12,325 genes, prioritizing the well annotated ones (see the “All” column, in the left side of each plot).

As expected, biased sources yielded a higher burden in terms of richness bias, publication-number bias and gene expression bias in their prediction output, and they generally prioritized a smaller fraction of ghost genes than their unbiased counterparts. A more nuanced landscape was observed in the final two confounders, namely PPI and GPT. Their interpretation is non-trivial: for example, a model may return a high GPT score either because the ML is performing well or because it is picking up literature bias—the proportion between these two opposed explanations may depend on the completeness of our current target list, which varies on a case-by-case basis. Cohen’s ***κ*** correlation analysis confirmed that the two biased channels’ results show one of the strongest correlations, although weak in absolute terms (***κ*** = 0.2-0.3, Supplementary Figure 3). Overall, these results confirm that using a literature-biased input dataset ***X*** comes with a more biased prediction ***Y***′, suggesting this may be the reason for the boost in ML performance that we observed in literature-biased channels.

### Tabular GhostBuster’s unbiased sources outperform biased sources in predicting novel biological discovery

The ultimate biological use for GhostBuster is to generate *novel* biological knowledge, i.e., identify “unsuspected” genes involved in a given cell function or disease. To this end, the top ranked gene list (in the test dataset’s prediction) should comprise an intermingling of (i) the target genes utilized for training; (ii) some genes that are not part of the “canonical” GO or MSigDB target class, but are still known in literature for playing a role in the phenotype of interest (i.e., have a high GPT score); and (iii) some genes that have never been associated with the provided phenotype before, i.e., have a low GPT score). To check whether this is the case, we measured the percentage of GPT-positive genes (i.e., GPT score of 6 or higher) for each phenotype and data channels (Supplementary Figure 4). Across all five channels, the top-ranked genes exhibited a nice mixture of known genes and unsuspected genes, in the ballpark of a 1-to-1 proportion. Unbiased sources generally produced a higher percentage of unsuspected genes, particularly LINCS (at 82%) and TCGA (at 84%); remarkably, the PPI channel yielded only 50% of unsuspected genes—a testament of the fact that GBA is strongly literature-biased, and biologically uninteresting.^51^ As we can see, unbiased channels’ prediction combines a strong presence of novel genes with a smaller influence from literature bias (as previously observed): this suggests it is likely to contain novel biological knowledge.

One way to assess if this is indeed the case is to retroactively test whether GhostBuster’s top ranked genes are among those whose involvement in the phenotype was recently discovered (and for this reason, did not appear in the target list). To pursue this question, we tasked the OpenAI GPT-4o API to generate a list of novel genes that were discovered between 2022 and 2025, for each of the selected cell functions or diseases. We then excluded the genes that were present in the target list for each of the queried phenotypes. A total of 130 recently discovered genes resulted from this query (Supplementary Table 4). We sampled and manually checked these “recently discovered genes” against the original literature papers, and they appeared for the most part in line with our prompt’s request, with relatively minor errors. For example, in the case of Parkinson’s disease (PD): it is indeed true that *TMEM175* received much attention in recent times, after a 2022 paper published in *Cell*^88^; conversely, *GBA1* has been known to play a role in PD for more than 3 decades,^89^ thus exemplifying an imperfect hit.

Next, for each phenotype, we measured how many of the 130 recently discovered genes were present among those top-ranked by GhostBuster, for each of the data channels, at varying cutoffs. To compare the performance of the biased and the unbiased channels, a logical *OR* was applied to measure how many recently discovered genes could be detected by any of the unbiased channels (LINCS *OR* TCGA *OR* STRING), versus any of the biased channels (GO *OR* PPI), at each of the cutoffs. This was a challenging attempt, not only due to the incomplete inaccuracy of the GPT query results, but especially considering the fact that “recently discovered genes” are inherently a literature-biased category. The core problem of literature bias is indeed the fact that literature preferentially expands towards genes that are rather well known in the first place, or similar to the already well-known ones. Nevertheless, the unbiased sources markedly outperformed the biased ones, by a factor of 2-3× (Figure 3). For example, with a cutoff at the 99^th^ percentile, hits from the unbiased predictors detected 25 of the 140, while those from the biased predictors only detected 10.

**Figure 3.**
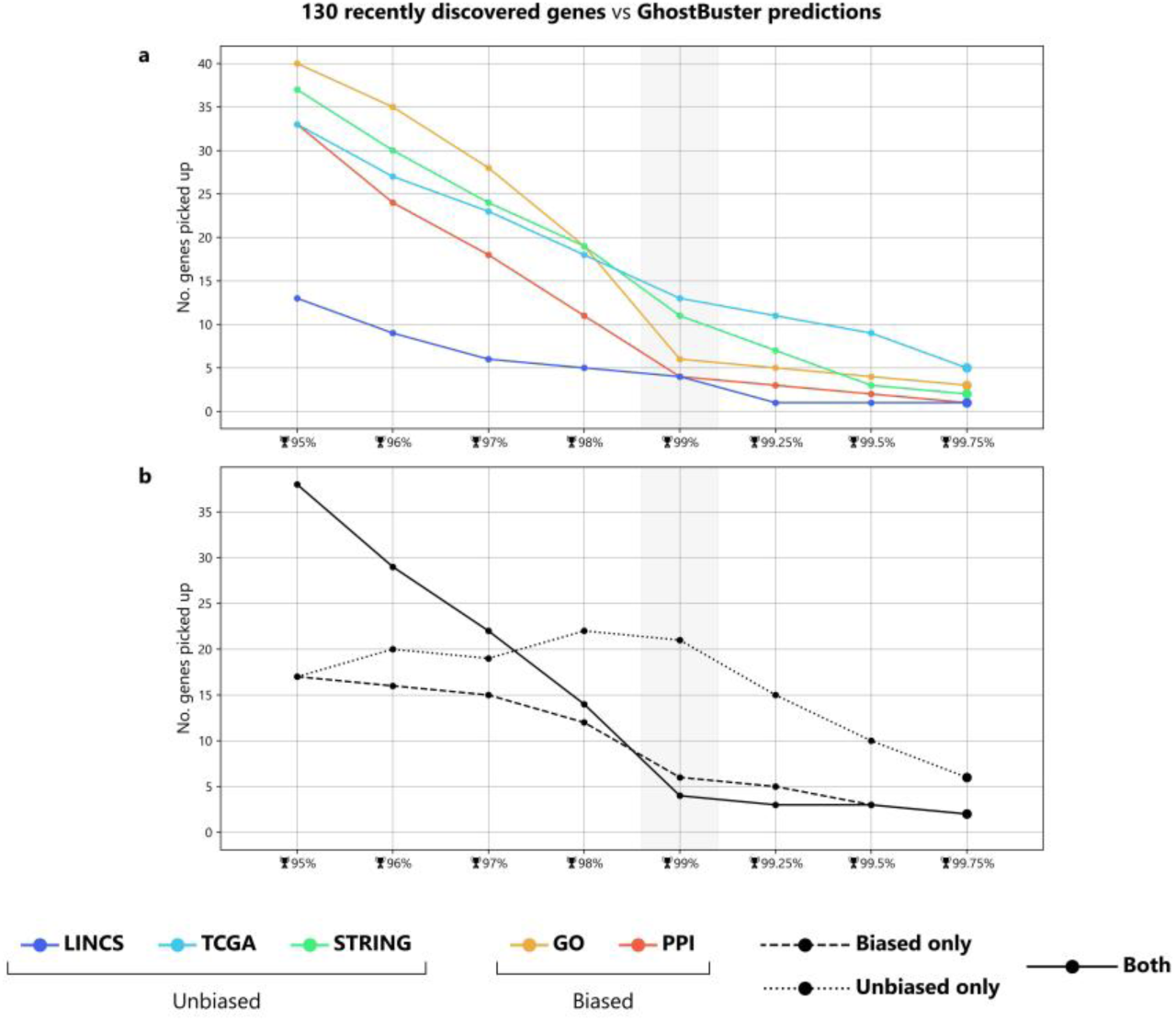
Recently discovered genes in the non-target top-ranked genes by Tabular GhostBuster. The plot shows the aggregated data from all of the 81 selected target lists. The output prediction ***Y***′ from each data channel (after excluding the positive genes) in Tabular GhostBuster was cross-compared with a list of genes that were recently (2022-2025) found to play an important role in the given target cell function or disease. The list of the recently discovered genes was created using OpenAI GPT-4o mini,^83^ and encompasses a total of 130 genes (see Supplementary Table 4). The top-ranked genes in Tabular GhostBuster’s output were sampled at different threshold levels (x-axis). Thresholding starts from the whole dataset (“All”); next, it continues with GhostBuster’s top hits located at/above the *x*^th^ percentile (🏆 *x*%), starting from the 95^th^ and moving up to the 99.75^th^. The analysis was repeated for all data channels: GhostBuster trained on the three “unbiased” data sources, namely LINCS (blue), TCGA (cyan) and STRING (green); next, GhostBuster trained on the “biased” data source, namely GO (orange); next, a simple GBA-type gene prioritization script, based on the physical protein-protein interactions in STRING (red). (a) The performance of the five input channels, individually. (b) The performance of the biased and unbiased channels, aggregated by a logical *OR*: for example, gene *A* is counted at threshold 99.25 of the unbiased channel, if and only if it appears among the top 99.25% genes in the LINCS channel, *OR* among the top 99.25% genes in the TCGA channel, *OR* among the top 99.25% genes in the STRING channel. Overall, the unbiased channels are generally better at detecting the recently discovered genes than the biased ones, by a factor of 2-3×.

### Tabular GhostBuster can be applied to make predictions on novel genes involved in cell functions or diseases

So far, we have established that Tabular GhostBuster is effective at predicting genes that may play a role in a given cell function or disease of interest, even if they are completely unannotated in literature. Specifically, the TCGA channel, and to a lesser extent the STRING and LINCS channels, seem to meet these objectives. We will now exemplify how Tabular GhostBuster can be applied to make novel predictions on a specific phenotype.

One way to maximize robustness is to select hits that were ranked high not only by one channel, but by two channels simultaneously, for example TCGA and STRING. Supplementary Figure 5 shows one such example for a cell function (autophagy), while Figure 4 shows one for a disease (Parkinson’s disease, PD). Let’s consider PD as an example, and check if the predictions are biologically plausible. Of note, we must bear in mind that the target list was the MSigDB list for PD, which in turn was sourced from a KEGG pathway that heavily focuses on mitochondrial mechanisms; as a result, we naturally expect a higher prevalence of mitochondrial genes in our prediction as well. Top-ranked, non-target genes highlighted by Tabular GhostBuster include the following:

- *MPC2* encodes the transporter that leads pyruvate into mitochondria, where it undergoes pyruvate dehydrogenase reaction. In 2024 it was identified as a potential drug target for PD and Alzheimer’s disease.^90^
- *TOMM7* encodes a subunit of a translocase, localized in the outer mitochondrial membrane. It was found in 2022 that its biallelic loss of function can lead to a progeroid syndrome.^91^ Despite having been long known to play a role in Parkin-induced mitophagy, and to influence the mitochondrial import of PINK1 (both Parkin and *PINK1* being known prominent PD genes),^92^ its role in PD pathophysiology is virtually unexplored.
- *NDUFA12* encodes a subunit of mitochondrial complex I. In 2024, it was identified as associated with cognitive decline in elderly women,^93^ while its involvement in PD has never been explored.
- *PRDX3* and *PRDX4* encode a mitochondrion-based and an endoplasmic reticulum-based peroxiredoxin, respectively. The former is known for being a phosphorylation target of LRRK2 (another major PD protagonist); its expression block increases vulnerability to oxidative stress in dopaminergic cells. In March 2025, it was found that by targeting it using gene therapy, it was possible to rescue degeneration in a PD Paraquat mouse model.^94^
- *CHCHD10* encodes a mitochondrial protein involved in cellular stress response and morphology of mitochondrial cristae; its mutations are known to cause frontotemporal dementia and amyotrophic lateral sclerosis. Only in 2019, it was found to be a central player in the pathophysiology of a familial form of PD.^95^

**Figure 4.**
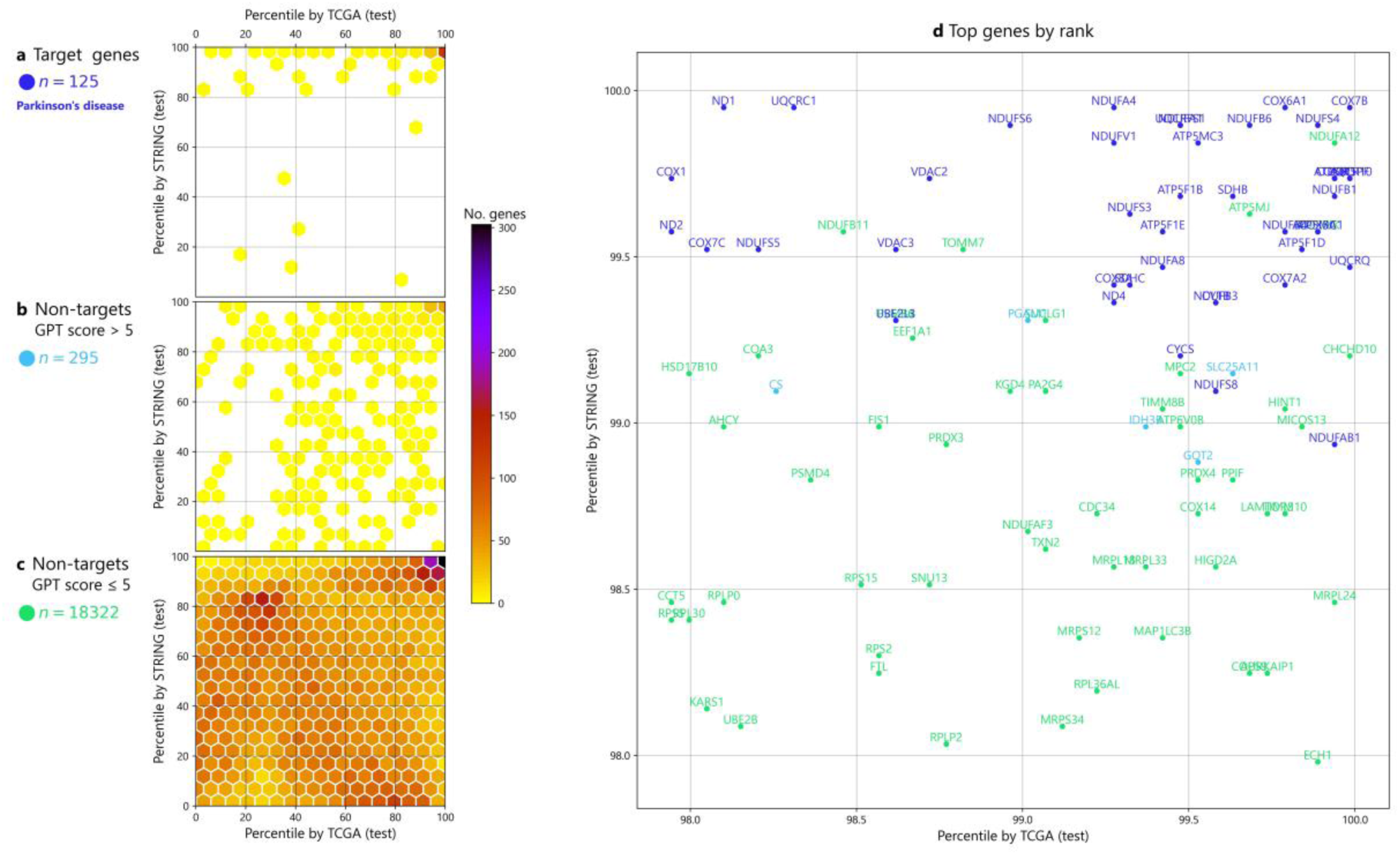
Prediction output of Tabular GhostBuster on Parkinson’s disease, cross-matching the TCGA and STRING unbiased channels, in the test dataset prediction. Tabular GhostBuster was trained to recognize 125 genes in the Parkinson’s disease pathway from the Kyoto Encyclopedia of Genes and Genomes (KEGG), utilizing the TCGA channel or the STRING channel as its training input. At the end of the training, a gene prioritization list was assembled by concatenating the rank-transformed test dataset predictions, for both models. Additionally, GhostBuster’s GPT control group was utilized to obtain GPT-4o mini-derived association scores between each gene and Parkinson’s Disease, in a standardized scale from 1 to 10 (color code). Finally, genes were organized in a scatter plot based on their rank in the TCGA channel (x-axis), versus their rank in the STRING channel (y-axis). (Left) Density plot, decomposed in three parts: namely (a) the 125 target genes (blue genes), (b) the non-target genes with a GPT score of 6 or higher (cyan genes), and (c) the non-target gene with a GPT score of 5 or less (green genes). In all three plots, agreement between the two channels can be observed in the top hits (top-right corner). (Right) (d) Scatter plot detail of the top 4 percentiles, at gene-level resolution. The green genes depicted here were simultaneously selected by both the TCGA and the STRING channel as top hits in the test dataset, despite their involvement in Parkinson’s Disease was largely unexplored in the scientific literature: this makes them the “interesting” hits for the sake of our prediction.

This pattern continues for dozens of other genes: as it is apparent, virtually all top-listers in Tabular GhostBuster’s prediction are either genes whose PD involvement was demonstrated very recently, or novel genes that showcase intriguing preliminary grounds in support of their involvement.

### Tabular GhostBuster can be applied to narrow down a gene’s pathway membership

Another interesting application of Tabular GhostBuster consists in training the ML model to recognize not only the general cell function, but also its many sub-pathways. This way, GhostBuster will not only provide the user with a prioritization list across the genome, but also, for every gene hit, it will estimate the likelihood by which that gene takes part in each sub-mechanism. To exemplify this point, GhostBuster was trained on three different groups of gene lists, sourced from GO^41^:

- DNA damage response, and 7 sub-mechanisms: double-strand break repair, positive regulation of double-strand break repair, positive regulation of homologous recombination, non-homologous end joining (NHEJ), nucleotide-excision repair (NER), base-excision repair (BER) and mismatch repair (MMR) (Figure 5).
- Autophagy, and 6 sub-mechanisms: DNA damage response, autophagy of mitochondrion, macroautophagy, microautophagy, positive regulation of autophagy and negative regulation of autophagy (Supplementary Figure 6).
- Generation of precursor metabolites and energy, and 8 sub-mechanisms: energy homeostasis, mitochondrial respiratory chain complex, oxidative phosphorylation, energy reserve metabolic process, glycolytic process, mitochondrial fission, mitochondrial fusion and mitochondrial translation (Supplementary Figure 7).

**Figure 5.**
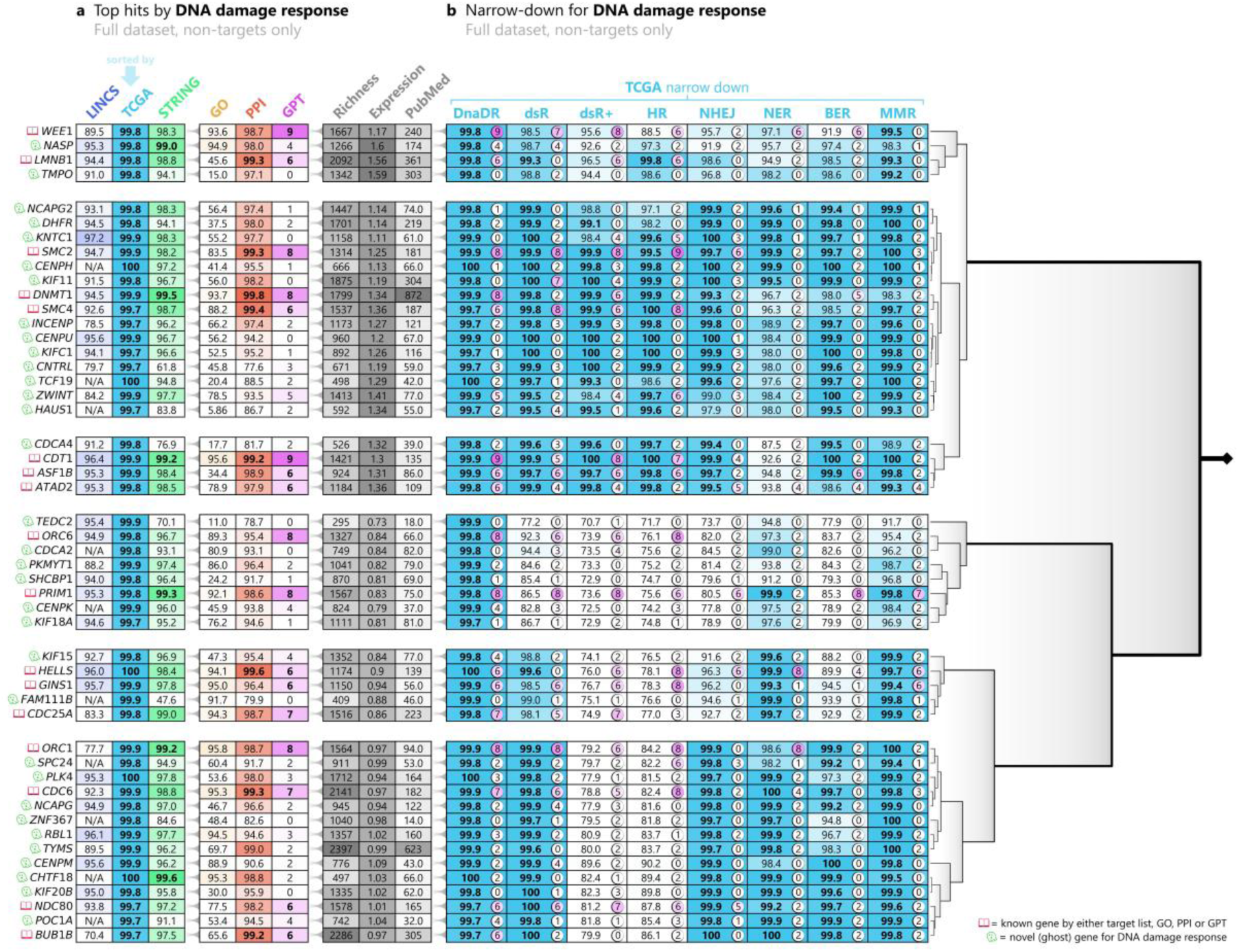
Tabular GhostBuster narrow-down plot across 7 sub-mechanisms of DNA damage response. Tabular GhostBuster was trained to recognize 813 genes in the “DNA damage response” list from Gene Ontology (GO), utilizing the TCGA channel as its training inputs (cyan arrow). At the end of the training, the top 50 non-target genes were selected, based on the prioritization list generated by such model in the full dataset, and are depicted here. The left part of the table showcases the percentile ranks generated by training Tabular GhostBuster against the “DNA damage response” gene list, based on the LINCS (blue), TCGA (cyan), STRING (green) and GO (yellow) channels; next, GhostBuster’s auxiliary PPI scores (red), GPT scores (pink); and the richness, average expression and PubMed scores (grey). A crimson book icon on the left side of genes’ symbols is assigned for genes that are already known for playing a role in the phenotype of interest, either because they are target list members, top-rankers in the GO model (>99^th^ percentile), top rankers in the PPI score (>99^th^ percentile), or GPT-positive (GPT score > 5). Conversely, a green ghost icon indicates that the gene was not predicted by any of the literature-biased sources, namely, the gene is not involved (or is marginally involved) in our current understanding of DNA damage response; hence, it is potentially novel (ghost) for this phenotype of interest. The right part of the table depicts the percentile ranks generated by training TCGA-based GhostBuster against the “DNA damage response” list, and 8 lists representing sub-mechanisms thereof. The pink circle on the right side of each percentile displays the standardized GPT score, associating the gene with each sub-mechanism: essentially, scores of 5 or less indicate that the gene has not been strongly associated with that sub-mechanism, in the scientific literature. The 50 selected genes are sorted based on hierarchical clustering, performed on the sub-mechanism percentile ranks. The clustering tree is depicted on the right of the main table, and helps identify blocks of genes with a potentially similar biology. DnaDR: DNA damage response, and 7 sub-mechanisms; dsR: double-strand break repair; dsR+: positive regulation of double-strand break repair; HR: positive regulation of homologous recombination; NHEJ: non-homologous end joining; NER: nucleotide-excision repair; BER: base-excision repair; MMR: Mismatch repair.

Because most of the predicted genes are largely uncharacterized, it is impossible to provide systematic evidence supporting the reliability of the predicted sub-mechanisms. As a result, the proof for such predictions will necessarily go through experimental validation.

### Tabular GhostBuster can be applied for the prioritization of intergenic GWAS gene hits

One of the most important open questions in gene disease prediction is the prioritization of statistically significant hits from genome-wide association studies (GWAS). These large-scale studies perform genomic characterization on thousands, or even tens of thousands of participants for a given disease of interest: such large sample size enables the detection not only of the high-penetrance genes that may cause the familial forms of the disease, but also of the low-penetrance risk variants that slightly increase the risk (or conversely, protective variants that decrease the risk), in a classic genotype-environment interaction paradigm. Although very powerful, GWAS studies are cursed by a high number of false positives, opening the doors for bioinformaticians to develop efficient prioritization tools. Furthermore, many of the statistically significant hits localize in non-coding regions, giving rise to an interpretability issue.^96–98^ When a hit falls in a region far away from any protein-coding genes, still, in most cases it exerts its action by controlling a coding-gene: in roughly ∼50% of the cases, this trans-regulated gene happens to be the closest protein-coding gene in the chromosome’s primary sequence to the hit locus.^99^ However, in the remainder 50% cases, the controlled gene can be located elsewhere, even on a different chromosome, and may end up localizing in proximity of the hit locus due to the 3D DNA folding. Traditionally, the closest genes in the primary sequence have been used as a proxy, although acknowledging that this may often lead to false positives.^96–98^

To showcase another potential use case of Tabular GhostBuster, we applied it on a GWAS prioritization task for 6 clinically relevant phenotypes, namely PD, AD, schizophrenia, type 1 diabetes mellitus, type 2 diabetes mellitus and metabolism. For each of these, we trained two independent models, differing by the nature of their respective target lists:

- The MSigDB-based model was trained to recognize a list of genes that are known for being mechanistically involved in the phenotype, sourced from MSigDB.^18^ As a result, this model scores how closely each gene “looks like” the known genes implicated in phenotype of interest’s pathway.
- The PheGenI-based model was trained to recognize a list of genes highlighted by GWAS studies for that phenotype, sourced from PheGenI.^21^ Such list includes genes housing (or neighboring) a statistically significant mutation, for which the identity of the target gene is known; it also includes genes that are the closest, at primary sequence level, to an intergenic hit, whose involvement is thereby uncertain. The inclusion of the latter in the target list is debatable, as only a portion of them will be true positives. We made the choice to include them, mainly to boost example numerosity. After all, GhostBuster’s job is to generalize the best rule that encompasses the target list *as a whole*: then, it follows that the inclusion of a minority of scattered false positives does not undermine the model’s prediction substantially, as the model only picks up factors that are *the most shared* across the entirety of the list. Overall, this model scores how closely each gene “looks like” the GWAS gene hits for the phenotype of interest.

After training both models, a prioritization plot was performed, while limiting the plot to only the genes that are intergenic hits for that phenotype, according to PheGenI. The general idea is that, the intergenic list contains a majority of false positives; of these, the genes that are most susceptible to be true positives are those that either match the shared factors among the mechanistically involved genes, or match the shared factors of the whole GWAS hit list (including the intragenic and near-genic); or even better, the “dual hits” that satisfy both of such criteria at once.

Representative results for PD, type 2 diabetes mellitus and schizophrenia are reported in Figure 6, Supplementary Figure 6 and Supplementary Figure 7, respectively. While it would be impossible to prove the validity of these results without performing a mechanistic study, a literature search for many of the single or dual hits suggests that their disease implication is strongly justified on theoretical grounds. For example:

- *HLA-DRB1* (dual hit for PD) is a critical component of the human leukocyte antigen system, making it a central player in inflammation-related diseases like neurodegeneration; unsurprisingly, limited evidence already exists that demonstrates a specific association between *HLA-DRB1* and PD^100^; interestingly, the interaction between alpha-synuclein and the HLA-DRB1 protein at intestinal level has been shown to be a determinant factor in an early-stage model of PD.^101^
- *DDRGK1* (single hit for PD) encodes an essential regulator of the endoplasmic reticulum homeostasis and the NF-κB inflammation pathway, both of which make its role in PD highly plausible.^102^
- *FITM2* (dual hit for type 2 diabetes mellitus) encodes an essential factor in lipid droplet formation, a process that is required to store saturated fatty acids into lipid droplets within adipocytes. Notably, the excess of saturated fatty acids is a known toxicity mediator on beta-cells in type 2 diabetes mellitus: although still very limited, experimental evidence has been disclosed that specifically highlights *FITM2* as a possible therapeutic target.^103^
- *CAMK1D* (single hit for type 2 diabetes mellitus) encodes a calcium/calmodulin-dependent protein kinase that participates in a wide variety of cellular processes; again, a small number of papers have confirmed its involvement in insulin release and insulin sensitivity, as well as its role as a genetic risk for type 2 diabetes mellitus.^104^
- *PDZD8* (dual hit for schizophrenia) encodes a lipid transfer protein involved in lipophagy; interestingly, a recent work showed that its knock-out in mice primarily affects the brain, leading to changes in emotional, cognitive and mnestic tasks, displaying some degree of overlap to schizophrenia.^105^
- *Nectin1* (single hit for schizophrenia) encodes a cell adhesion molecule involved in the formation and maintenance of synaptic joints, as well as neuroplasticity. Although the specific role of nectin 1 in schizophrenia has not been well characterized yet, cell adhesion molecules in general (and the nectin family in particular) are thought to be some of the core mechanistic players in neurodevelopmental disorders, including schizophrenia.^106^

**Figure 6.**
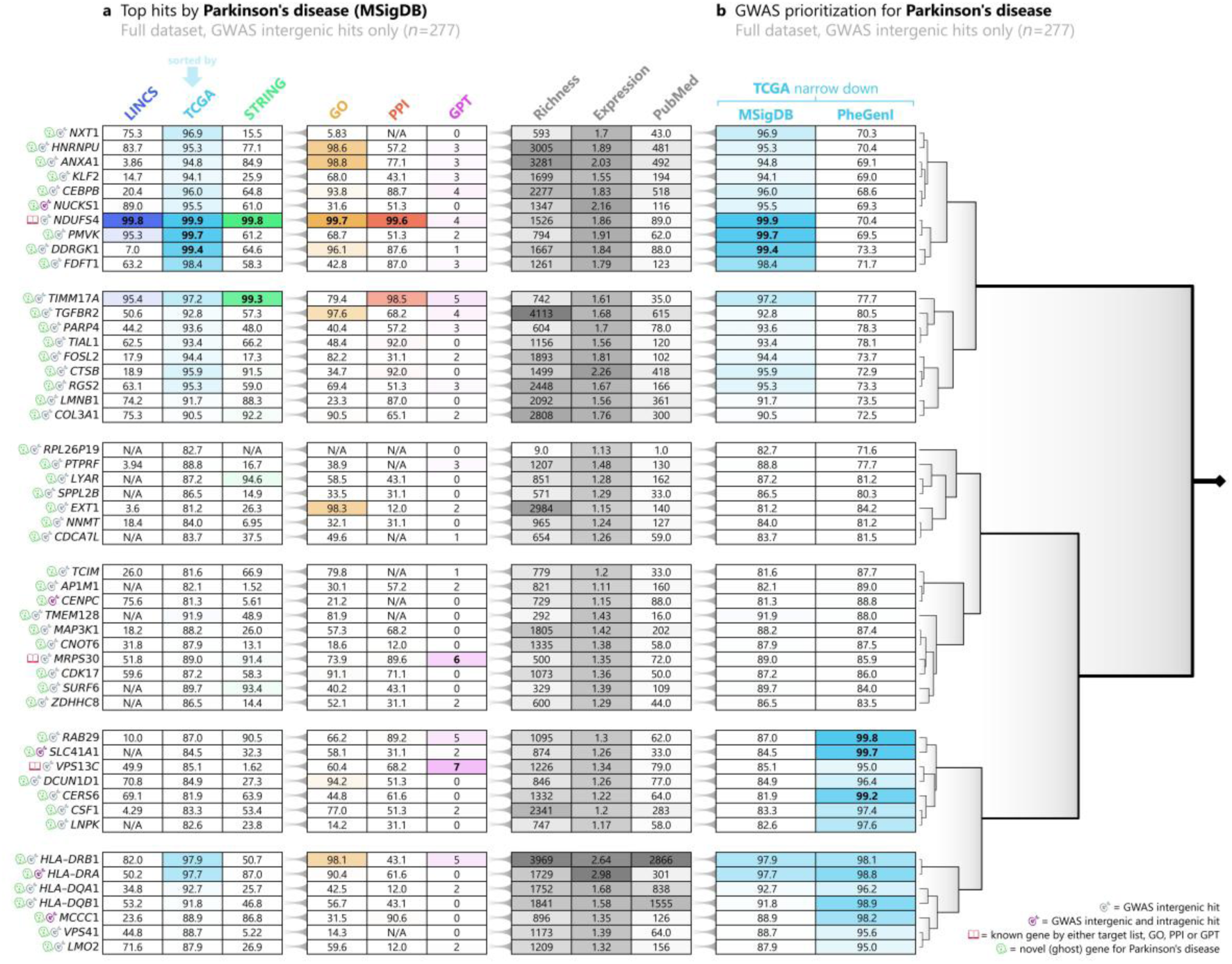
Tabular GhostBuster prioritization of intergenic GWAS gene hits in Parkinson’s disease. Tabular GhostBuster was trained to recognize 130 genes in the “Parkinson’s disease” (PD) pathway, sourced from the Molecular Signatures Database (MSigDB); and 75 genes that were found as hits in genome-wide association studies (GWAS) on PD, sourced from the Phenotype-Genotype Integrator (PheGenI). The latter included genes that hosted an intragenic or near-genic GWAS hits, as well as “intergenic gene hits”, that is, genes that were the closest in the primary sequence to an intergenic GWAS hit. Training was performed utilizing the TCGA channel as its training inputs (cyan arrow). At the end of the training, only the 184 intergenic gene hits were selected: of these, the top 50 genes were utilized for plotting, based on the prioritization list generated by the model trained on MSigDB in the full dataset, and are depicted here. Because intergenic gene hits are known for including a 50-75% share of false positives, the aim of this plot is to identify which ones are the most likely true positives, based on their similarity to the mechanistically involved genes (MSigDB), or to the GWAS hit list as a whole (PheGenI). The left part of the table showcases the percentile ranks generated by training Tabular GhostBuster against the PD MSigDB gene list, based on the LINCS (blue), TCGA (cyan), STRING (green) and GO (yellow) channels; next, GhostBuster’s auxiliary PPI scores (red), GPT scores (pink); and the richness, average expression and PubMed scores (grey). A grey target icon on the left side of genes’ symbols indicates that a gene is only a GWAS intergenic hit, while a purple target icon indicates that it is both an intergenic and an intragenic GWAS hit. A crimson book icon is assigned for genes that are already known for playing a role in the phenotype of interest, either because they are target list members, top-rankers in the GO model (>99^th^ percentile), top rankers in the PPI score (>99^th^ percentile), or GPT-positive (GPT score > 5). Conversely, a green ghost icon indicates that the gene was not predicted by any of the literature-biased sources, namely, the gene is not involved (or is marginally involved) in our current understanding of PD; hence, it is potentially novel (ghost) for such phenotype. The right part of the table showcases the percentile ranks generated by training Tabular GhostBuster against the PD MSigDB list, and the corresponding PheGenI list. We define as “single hits” the genes that reach the 99^th^ percentile in the MSigDB channel or in the PheGenI, whereas “dual hits” are the genes that achieve the 99^th^ percentile in both channels at the same time. The 50 selected genes are sorted based on hierarchical clustering, performed on the MSigDB and PheGenI percentile ranks. The clustering tree is depicted on the right of the main table, and helps identify blocks of genes with a potentially similar biology.

The list could extend considerably. To recapitulate, the reported hit genes were selected among the intergenic gene hits, on the basis that they showed similarity to either the known disease pathways, or the rest of the GWAS hit list as a whole, or both. Such striking coincidence is allusive that each of these genes is among the true positive hits. In fact, the reported literature analysis on all the representative example genes confirmed that their involvement is biologically plausible. This panel exemplifies how Tabular GhostBuster can be deployed for a systematic prioritization of GWAS intergenic hits.

### Graph GhostBuster trained on literature-unbiased sources determines less literature bias in the predictions

Graph GhostBuster was trained on an edge prediction task, whereby nodes represented genes, and edges represented directed gene-gene relations. Positive examples of gene-gene edges were sourced from 21 individual levels from the Biology Mega Graph knowledge graph (Supplementary Table 2), corresponding to different biological or biochemical gene-gene relation types (e.g., `methylates`, `relocates`, `downregulates_quantity_by_repression`, etc.). The 21 levels were selected in terms of number of edge annotations; filtering the levels with at least 460 annotations, and discarding the others. Conversely, negative edges were sourced in two ways, corresponding to two independent edge prediction tasks:

- **R task** (random negatives): edges were randomly generated, in a 50:1 proportion to the number of positive examples. The rationale was based on the sparsity of the positive gene-gene edges (density: min 0.00029%, max 1.54%, mean 0.12%). As a result, by taking random edges, we can assume to be introducing a very limited proportion of false negatives in the negative edge dataset.
- **C task** (curated negatives): given a positive layer of the Biology Mega Graph (say, “`upregulates activity`”), edges were sourced from its corresponding antonymic layer (“`downregulates activity`”), when available; this was only possible for 10 out of 21 levels, namely, the upregulation- and downregulation-based layers. The rationale was that, when a gene upregulates another gene, usually it does not *also* downregulate the same gene; this is of course an imperfect assumption, as some regulatory relations are known for being cell type-dependent, even in their sign.^36^

Node features were sourced from the same data channels used to train Tabular GhostBuster, namely the three literature-unbiased sources LINCS, TCGA, STRING; and the literature-biased source GO.

A graph convolutional neural network (GCN) model was employed, in a classic edge prediction task. Training was conducted independently for each of the 21 gene-gene relation types, for each task (R or C), and for each of the 4 data sources, resulting in (21 + 10) × 4 = 124 models being trained. In each model, the positive and the negative edges were randomly split into a training, validation and test dataset, in 8 : 1 : 1 proportions, respectively.

At the end of the training, performance was measured by ROC-AUC in the test dataset (Figure 7). Surprisingly, the TCGA channel achieved a comparable performance to that of the biased GO channel, with most ROC-AUCs falling in the 0.90-0.95 range. Conversely, the LINCS and STRING markedly fell back, performance-wise. Across the 29 relation type/task combinations, TCGA was reported as the best performer in about 50% cases—even more successful than the corresponding result from Tabular GhostBuster, where the use of literature-unbiased sources came with a small penalty in the ML metrics. Additionally, it further confirmed TCGA’s role as an ideal, unbiased information source for GAP tasks.

**Figure 7.**
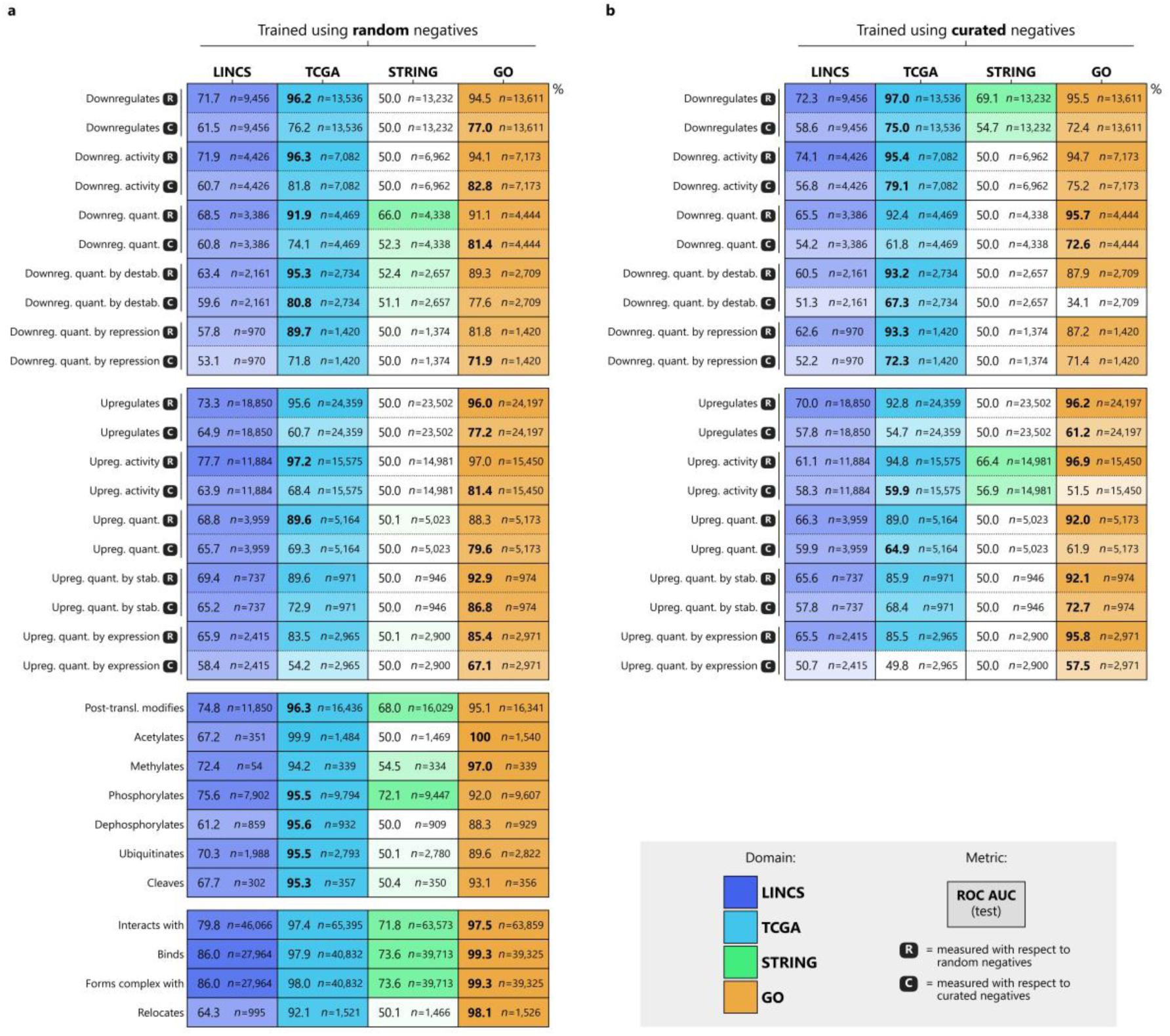
Graph GhostBuster performance on the 21 interaction modalities, measured by the ROC-AUC in the test dataset prediction. The plot measures Graph GhostBuster’s performance in terms of receiver operating characteristic area under the curve (ROC-AUC), in the test dataset prediction. A graph convolutional neural network (GCN) was trained independently for each of the 21 interaction modalities (rows), in an edge prediction task, where positive edges were sourced from the Biology Mega Graph. Some interaction modalities came with an antonymic counterpart (e.g., “upregulates quantity” is the antonym of “downregulates quantity”); for these modalities that admitted an antonym, two models were trained: one model (left table) using random negative edge examples, generated in a 50× quantity of the positive edge examples; another independent model (right table) using the antonymic modality as the “curated” negative edge examples; in either case, ROC-AUC was always measured both with respect to the random negatives and the curated negatives (“R” and “C” black icons). Node features were sourced from four data sources: three “unbiased” data sources, namely LINCS (blue), TCGA (cyan) and STRING (green); and one “biased” data source, namely GO (orange). The ROC-AUC scores are expressed as percentages (%), and are accompanied by the number of positive edge examples available in their respective data source’s namespace. All cells are color-coded based on the performance score, in a non-linear fashion. Overall, TCGA and GO achieved a comparable performance, whereas GO and STRING often failed to learn the task, despite being training repeated across multiple learning rate values, picking up the best setting by grid search.

Next, we checked whether the use of a literature-unbiased data source paid off in terms of literature bias in the results, analogously to what had been done for Tabular GhostBuster. The confounders were again metrics that reportedly correlate with literature bias,^3,37,42^ namely, for every edge: (i) the number of GO and MSigDB annotations of the two genes delimiting the edge (“richness”); (ii) their coverage in PubMed articles; (iii) their average log-transformed transcriptomic expression across all RNA-sequenced tissues in TCGA; (iv) the percentage of ghost genes, defined as genes in the lower two thirds of the richness distribution; (v) the percentage of ghost genes, defined as genes in the lower two thirds of the PubMed coverage distribution. They were measured by cutting the GNN predicted edges at different thresholds based on the logits in the predicted adjacency matrix, starting from the complete adjacency matrix (no threshold); next moving on to the genes in the top 50,000 highest-ranked predicted edges; then the top 30,000, etc., up to the top 100 (Figure 8).

**Figure 8.**
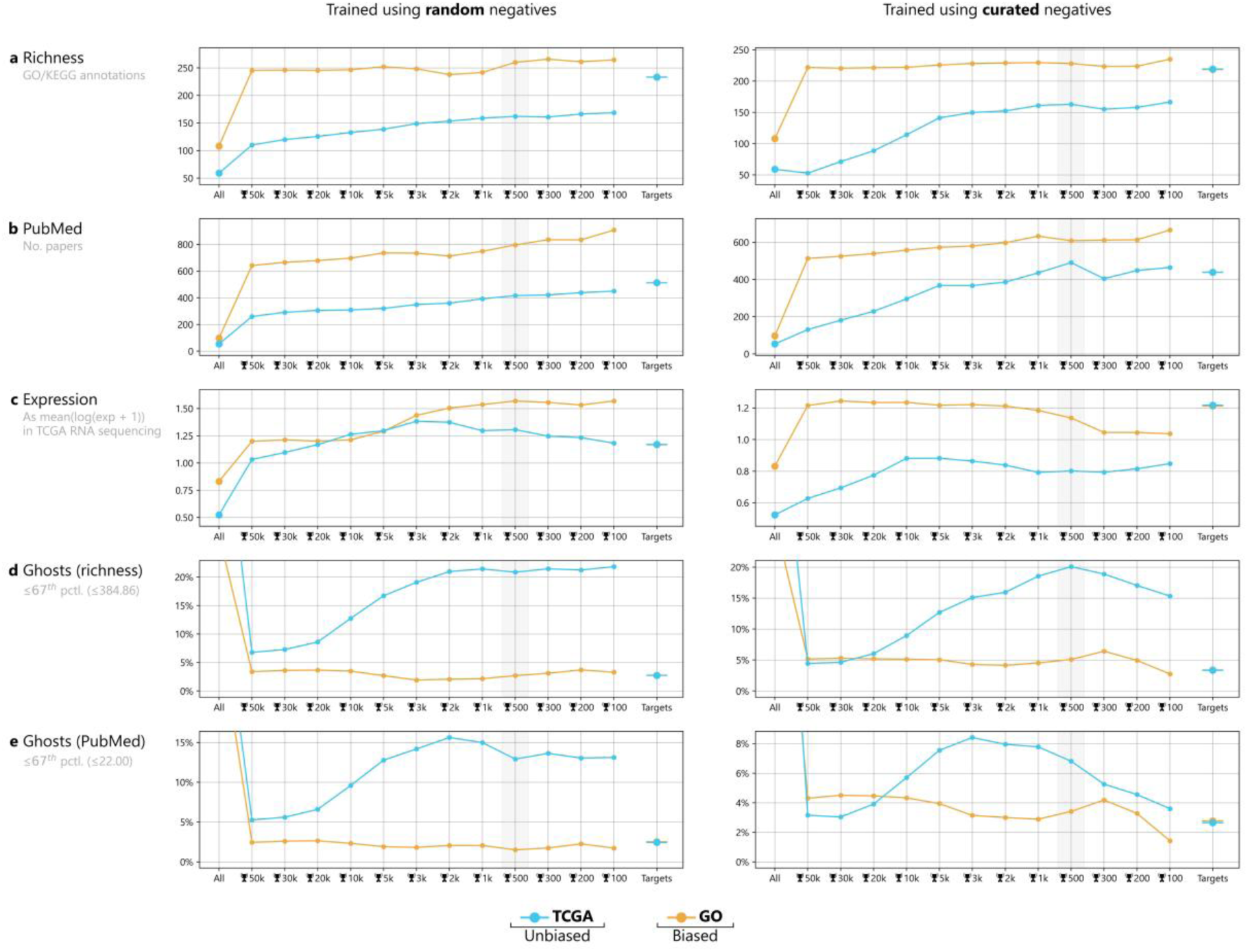
Literature bias confounders in the genes involved in the top-ranked edges by Graph GhostBuster. The plot quantifies 5 literature bias confounders in the prediction of Graph GhostBuster across all 21 interaction modalities, with respect to the data channel that was provided in their input. Confounders (y-axis) were evaluated in the output adjacency matrix from each data channel, by sampling the top-ranked edges with respect to their logits at different threshold levels (x-axis). Thresholding starts from the whole dataset (“All”); next, Graph GhostBuster’s top *n* edges (🏆 *n*), starting from 5,000 and moving up to the top 10; finally, on the right side, the average values for the positive edges (targets) are shown for comparison. The analysis was repeated for two data channels: Graph GhostBuster trained on the best-performing “unbiased” data source, TCGA (cyan); next, Graph GhostBuster trained on the “biased” data source, namely GO (orange). The other data channels (LINCS, STRING) were excluded due to their unsatisfactory performance (see Figure 7). The explored confounders are: (a) the number of GO and MSigDB annotations of the predicted genes (“richness”); (b) the coverage in PubMed articles; (c) the average log-transformed transcriptomic expression across all tissues in TCGA; (d) the percentage of ghost genes, defined as genes in the lower two thirds of the richness distribution; (e) the percentage of ghost genes, defined as genes in the lower two thirds of the PubMed coverage distribution. The remainder two confounders from Tabular GhostBuster (namely the number of protein-protein interactions with the target gene members, and the GPT score) could not be calculated, as they were based on the premise of a positive gene list, which was not applicable to Graph GhostBuster. Overall, predictions from the biased source generally yield a higher bias across all the explored confounders.

Like in the case of Tabular GhostBuster, once again, the biased source (GO) reported a higher literature-bias burden over the best-performing literature-unbiased source (TCGA) across every explored confounder, thus confirming the hypothesis.

### Graph GhostBuster can be utilized to predict gene interactors with modality-specific resolution

So far, we have established that Graph GhostBuster is effective at predicting gene-gene interactions at full mechanistic granularity, namely distinguishing biological interaction types such as up/downregulation, activation/deactivation, phosphorylation, acetylation, etc. However, the ultimate step is to check whether novel biological insight can be extracted from such predictions.

Of note, because the current implementation relies on a classic dot product-based GCN, the output is agnostic of directionality (e.g., *A* `phosphorylates` *B*, versus *B* `phosphorylates` *A*); however, directionality is usually inferable from the biological background of the two genes (e.g., if *B* is a kinase, then it is *B* that `phosphorylates` *A*, and not vice-versa).

As an example, let’s consider again alpha-synuclein (αS protein, *SNCA* gene). Among the top 28 hits for up- or downregulation-related terms (Supplementary Table 5), 18 interactors had already been annotated in the Biology Mega Graph; conversely, 13 of them were novel, including the following:

- *NOTCH2* (hit for `downregulates_quantity` in *SNCA*) encodes one of the notch receptors, which are a cornerstone of the ontogenesis of multiple tissues, including the brain. Interestingly, *NOTCH2* has been often found significantly dysregulated in alpha-synucleinopathy patients. For example, a recent transcriptomic panel pinpointed the notch pathway, and specifically *NOTCH2*, as the most upregulated pathway in PD patients’ blood.^107^ Likewise, a “major intrinsically disordered NOTCH2-associated receptor 2” (*KIAA1024L*) was recently identified as a possible central mediator of αS vulnerability, as its knock-out recapitulates many of the PD hallmarks, including specific dopaminergic loss in the substantia nigra.^108^
- *PTK2* (hit for `upregulates_activity` in *SNCA*) encodes protein tyrosine kinase 2, generally known for being a high-level regulator in cell cycle and survival. Interestingly, research from the early 2020’s highlighted a key role in tau- and TDP-43-mediated neurodegeneration, with specific focus on vulnerability to the ubiquitin-proteasome system impairment. Kinase inhibitor screening highlighted it as the best neurotoxicity suppressor on a proteasome impairment model^109^; the mechanism was later identified to involve PTK2’s ability to phosphorylate p62, regulating the p62 pathway which promotes autophagy and misfolded proteins’ turnover.^110^ While *PTK2*’s potential role in αS vulnerability has not been investigated yet, it remains extremely plausible based on such background.
- *MLC1* (hit for `upregulates_quantity` in *SNCA*) encodes an astrocytic membrane transporter, which has been implicated in astrocyte activation—a key component in the pro-inflammatory mechanisms that are shared across virtually all neurological conditions, including neurodegeneration, stroke, trauma and epilepsy. A specific relation with αS has never been investigated.^111^

As an alternative example, let’s consider TDP-43 (*TARDBP* gene), which is the central protein in TDP-43 proteinopathies such as amyotrophic lateral sclerosis (ALS) and the TDP-43 subtype of frontotemporal dementia. Again, among the top 28 hits for up- or downregulation-related terms (Supplementary Table 6), 16 interactors already appeared in the Biology Mega Graph, whereas 12 were novel, for example:

- *CDKN1A* (hit for `downregulates_quantity` in *TARDB*) encodes a cyclin-dependent kinase inhibitor, which is a direct transcriptional target of p53 and has been implicated in neuronal death. Most notably, a 2018 paper explicitly reported that “expression of Cdkn1 reduces endogenous TDP-43 levels suggesting a reciprocal interaction with TDP-43”^112^; this matches exactly Graph GhostBuster’s prediction, despite this relation did not appear in the Biology Mega Graph that was used to train it.
- *SMAD3* and *SMAD4* (hits for `downregulates_activity` in *TARDB*) encode some of the signal transducers of the TGB-β pathways, involved in cell growth, differentiation and apoptosis. Interestingly, a paper reported the accumulation of various SMAD proteins within TDP-43 intracellular inclusions found in the spinal cord of ALS patients, suggesting that TDP-43 accumulation brings about an impairment of the TGF-β pathway.

Strikingly, almost all of the novel biological predictions from Graph GhostBuster are biologically interesting, and plausible based on the limited existing literature. These representative examples confirm that Graph GhostBuster is capable of predicting novel biological knowledge, that was not part of its training dataset.

In conclusion, the elegance and usefulness of both Tabular and Graph GhostBuster ultimately lie in their unbiased quality: the ability to pinpoint, within an ocean of reported “limited evidence”, which hypotheses are the worthiest of being explored further, making the tool widely adoptable by the biological community.

## Discussion

Scientific literature suffers from well-known disparity issues, by which it favors a small subgroup of highly annotated genes. Research social structures often encourage investigations to keep focusing on the genes that we already know best, in a sort of “bandwagon effect”.^37^ Conversely, the (relatively infrequent) efforts to characterize select ghost genes have often led to major scientific breakthroughs. Literature bias also affects computational efforts in the fields of gene function prediction (GFP), gene disease prediction (GDP) and gene network prediction (GNP). Only a small minority of papers take into account the extreme incompleteness in our gene annotation, even in *H. sapiens*.^3,37,42,113^ Much ML literature claims to improve prediction accuracy based on ML performance metrics (ROC-AUC, PR-AUC, etc.) *as if* the training target was a perfectly complete ground truth. This observation led us to the following fundamental question, about existing literature—a sort of “self-fulfilling prophecy” hypothesis.

Let’s clarify this question with an example. If both the ML training data and the target are literature-biased, they will contain a mixture of some true and some false statements (e.g., a ghost gene is believed *not* to play a role in a certain cell function, while in actuality it does); then, a model that yields record-breaking AUC may do so because it better learns the lies as well as the truths. For example, let’s say that gene *A* has functional annotations *f*_2_ and *f*_6_; but in actuality, it contributes to *f*_1_, *f*_2_, *f*_3_, *f*_4_, *f*_6_. If a model predicts *f*_2_ and *f*_6_, it will score a better AUC than a model that predicts all the real ones. Because ML publications get judged essentially by their prediction’s AUC against the training target, they are encouraged to overfit the errors in the annotations, rather than predict real, novel biology.

This issue may also explain why the biological community has shown little trust and adoption of ML-based predictors, despite AUC’s consistently building up in the recent years.^4^ To tackle this complex bioinformatic question, we developed GHOSTBUSTER, to the best of our knowledge the first ML architecture that combines modern deep learning computational firepower with completely literature-unbiased sources for GFP, GDP and GNP.

GhostBuster was trained independently on 3 literature-unbiased and 2 biased channels. Remarkably, Tabular GhostBuster’s performance clearly correlated with the amount of bias that was present in the training data, confirming the existence of a short-circuit phenomenon. Subsequent confounder analysis revealed that, in both Tabular and Graph GhostBuster, the use of a biased information channel exacerbated literature bias in the predictions themselves, rendering them less biologically novel, along the lines of “more of what we already know”.

The importance of explicitly making this comparison cannot be overstated, as this short circuit phenomenon has never been systematically studied in pre-existing literature, to the best of our knowledge. On the contrary, the performance-boosting effect has been not only accepted, but sometimes even sought after. A notable example is the winner of the recent Critical Assessment of Function Annotation (CAFA). CAFA encourages submitters to employ “any other data available to them” to train their models, and judge the models based on the gene annotations that are added to GO in the ∼6 months ensuing the deadline.^114^ However, biocuration comes with a certain delay: evidence is first published in pre-prints, then in literature, and only weeks or months later, it finds its way into the harmonized repositories. If one scraped today’s literature, they would probably be able to anticipate the annotations that will be added to GO in the next weeks or months. This is, in a nutshell, the principle that (explicitly) inspired GOCurator/GORetriever, winner of the CAFA 5 challenge in 2023.^115^ Indeed, their approach is invaluable to help biocurators harmonize the published literature, yet, it does not aim to add *novel* biological knowledge. Overall, our analysis on GhostBuster’s performance concluded that ML scores alone are an unsuitable—and sometimes even deceptive—metric for GFP/GDP/GNP tasks, as they may inversely correlate with the biological “interestingness” of the predictions.

To provide a better-suited metric, we employed large language models such as OpenAI’s GPT-4o mini to systematically quantify the degree of association between a gene and each of the 81 phenotypes of interest in Tabular GhostBuster, based on existing literature. Results showed that indeed, models trained from literature-biased data tend to output genes whose involvement in the phenotype was already known in the first place; conversely, unbiased models are considerably less affected by this issue, and generally output a healthy mixture of known genes and novel genes—the latter being the “interesting” ones. Similarly, we employed GPT 4o to generate a list of genes, whose involvement in the phenotype of interest was only discovered between 2022 and 2025; once again, unbiased sources outperformed the biased ones at predicting such genes, by a factor of 2-3×.

As far as we are aware, GhostBuster represents the first case in bioinformatic literature where an LLM was utilized for such tasks. One could argue that LLM scoring comes with incomplete accuracy; and indeed, even a small, sampled check in our own LLM results immediately shows that a minority of the LLM’s scoring predictions are incorrect, but they become negligible in such a wide array of queries. We are of the opinion that, by the next few years, systematic LLM scoring across millions of gene-function pairs or gene-disease pairs will become the go-to approach, for every bioinformatic question where manually curated data are not available.

Finally, we validated the biological relevance of tabular and Graph GhostBuster by showcasing four applications that may be of interest for a hypothetical, real-world biologist. These are namely: predicting novel genes involved in a function or disease; narrowing down the exact gene’s pathway membership within a broader function; prioritizing the intergenic gene hits from genome-wide association studies (GWAS); and predicting novel gene-gene interactions at full biological granularity. The latter is also largely novel: due to the popularization of “protein-protein interactions” (PPIs) as a general, yet ambiguous term (encompassing anything from physical interactions to phylogeny, coexpression and even indirect transactions), most authors have deployed edge predictors on these broad-scoped networks, rather than exploring them at the level of individual relation types. Across all the manual literature searches that we performed on GhostBuster’s outputs, the top-listers always included a mixture of known and novel genes, where most of the novel ones still offered plausible explanations that supported their potential involvement in the considered cell function, disease, or interactor. This confirms that GhostBuster succeeded in its mission to become the first bioinformatic tool to efficiently prioritize genes on GFP, GDP and GNP tasks, in a literature-agnostic fashion.

### Tabular GhostBuster architecture

The choice of the ML model was one on which we devoted considerable attention, starting from Tabular GhostBuster. Its task differs from most of the published ones in a fundamental way: most papers predict *all* (or at least, multiple) gene functions in parallel, rather than one by one. Their target matrix ***Y*** is usually made up of hundreds of columns, whereas our ***Y*** is just a single column vector.^116,117^ The reason underlying our choice has to do with cross-validation (CV) stratification. Considering that most of the target classes fall in the 50-200 numerosity range, and that CV is customary for rigorous performance measurement in most GFP and GDP literature,^49,51,52^ the 10% validation dataset is reduced to as few as 5-20 examples. Stratification is imperative to guarantee that an equal number of examples is present in each fold. For this reason, we made the choice of training an independent model for each of the target phenotypes, even if this comes with substantially higher computational costs.

The next question was, how to deal with the fact that our gene prioritization task in Tabular GhostBuster is not a classic positive-negative classification, but rather, a positive-unlabeled one (or alternatively, a hidden positive one).^96^ Our target is a subset of genes that are positive for certain (the target list), while we do not know if the other genes are true negatives (genes that do not participate in the phenotype) or false negatives (genes that actually participate, but the scientific community has not found out yet); existing data sources such as GO store almost exclusively positive information, while negative information (genes that do *not* play a role in a phenotype) is extremely limited.^96,118^ Indeed, there has been recent ML literature providing various adaptations to train deep learning on positive-unlabeled tasks.^119^ However, these architectures remain a small niche in the ML community, and were not designed to face the exceptionally challenging conditions posed to GhostBuster (extreme unbalance, curse of dimensionality, etc.).

One could argue, why not to simply prioritize genes based on the logits from a standard classifier. The problem is that most classifiers are only trained to correctly allocate the examples above versus below the 0.5 threshold—there is no additional drive for the classify to “behave” in any specific way within the [−∞, 0.5) and [0.5, +∞] intervals. However, the one family of architectures that inherently have such drive are the margin-based ones, such as SVMs. These algorithms fit a geometric boundary “around” the known positives, and then quantify the geometric distance (i.e., margin) between each example and the boundary. To combine the power of deep learning with margins—thereby taking the good of both words—we eventually settled on an encoder-decoder combo, where the deep-learning encoder learns a low-dimensionality latent representation of the genes, and the SVM decoder takes over to map these embeddings into margins. Of the 150+ papers that we reviewed in the introduction throughout the GFP and GDP literature (see Table 2, Table 3 and Table 4), only STRING2GO has adopted a similar structure, although claiming different motivations.^120^

As an alternative to SVMs, we could have also utilized ensemble-based models, such as random forests, as others have done; GENIE3 (2010)^121^; Yan *et al.* (2010)^122^; Browne *et al.* (2015)^123^; GRN-transformer (2022).^124^ The idea is that each classifier in the ensemble “votes” on the class label for each example, thus forcing the logits to behave like an actual probability. While this method may work well in proximity to the boundary, it would lose generality for genes that are farther away, making it impossible to tell whether an all-0 gene is reasonably close to the boundary or very far away from it. On these grounds, we gave our preference to SVMs.

From a purely formal perspective, despite the use of margins, GhostBuster does not fully address the issue of the positive-unlabeled setting: we are still teaching our model that the negatives are “real negatives”—which is biologically incorrect, as they are technically unlabeled examples. This issue is considerably mitigated by the fact that GhostBuster weighs the positive and the “negative” examples inversely proportionally to their abundance. Indeed, among the ∼20,000 non-target genes, there will be some that are positive in biological reality, while we are explicitly teaching the machine that they are reliable negatives. Still, the impact of such wrongful teaching will be in the order of 1/200 of the impact exerted by each of the target members, rendering it negligible. Based on these considerations, we argue that, despite the formally incorrect treatment of the unlabeled examples, from a practical standpoint GhostBuster is a highly optimized compromise. To support this claim, across the wide world of GFP and GDP literature, only a small minority of papers have formally accounted for the unlabeled issue,^96,118^ while the others have generally not brought up the problem.

### Graph GhostBuster architecture

Moving on to Graph GhostBuster, and its GCN architecture: while GCNs internally utilize an encoder-decoder structure, Graph GhostBuster does not employ margins, thus contradicting the prioritization assumption that we just highlighted. This was a deliberate choice, in order not to overload the—already substantial—technical complexity of training a GNN across as many as 36,000 nodes, with node feature vectors as large as 280,000. It follows that, in the current implementation, any use where Graph GhostBuster’s logits are employed to *rank* the predicted interactors of a given GOI is formally incorrect, even if, in practice, it still usually outputs sensible results. However, for most biological applications, ranking is not strictly needed. For example, a biologist that is studying αS in microglia may be looking for a novel kinase, expressed in microglia, that has the ability to phosphorylate αS. After intersecting the predicted interactors from the “`phosphorylates`” model, with the kinases expressed in the microglia, and excluding the already known ones, the list of kinases to be experimentally investigated goes down to just a handful genes, that can be individually checked in the wet lab.

Another major limitation is that, due to the use of classic GCNs, the current version of Graph GhostBuster is incapable of distinguishing the subject from the object of a biological action in its output. As exemplified, this information can usually be inferred from the known biology of the involved genes, as long as they are not completely uncharacterized.

Both such limitations are to be listed among the future developments of Graph GhostBuster. Firstly, daisy-chaining an SVM with the GCN’s decoder operation in order to output a continuous margin, consistently with Tabular GhostBuster. Secondly, replacing the standard GCN with an architecture that supports directional output, such as PyTorch Geometric’s `GATConv` (graph attention convolution).

### Training data

Finally, some considerations are due, regarding the choice of the input data channels: LINCS, TCGA and STRING. The LINCS L1000 dataset offers over 3 million gene expression experiments, where cell cultures were subjected to a variety of stimuli (perturbations); these were aggregated into 1.2 million differential signatures, highlighting the perturbation’s effect over the rest state.^78,79^ The main strength of this approach lies in the fact that it picks up causative effects, rather than just correlation or association; most perturbations, such as chemical compounds, would activate or block single genes or pathways, thus pinpointing their causal, downstream effects. Another strength is that, while these are not single-cell data, their culture-based setting implies that samples are biologically homogeneous—though mostly in cancerous or immortalized conditions. Conversely, the main drawback lies in the L1000 technique itself, which was designed to prioritize scalability: its intrinsic noise is further exacerbated by the fact that only 978 genes were directly measured, whereas the rest were ML-derived. While LINCS was an interesting attempt for GhostBuster, especially considering that it had never been used for GFP/GDP/GNP purposes before (except for Endeavour, which however did not make use of ML^125^), such drawbacks took a substantial toll in terms of the ML performance.

The Cancer Genome Atlas (TCGA) offers the methodological advantages of employing state-of-the-art RNA sequencing across over 25,000 samples, all processed using a standardized protocol. The use of bulk RNA sequencing maximizes sensitivity for weakly expressed genes, and its multi-organ scope empowers it to convey broad pleiotropic information, enhancing GhostBuster’s generalizability. However, the cancerous nature of most samples introduces biological complexity due to both driver and passenger mutations. Unlike LINCS, which often isolates single pathway effects by targeted perturbations, cancer reflects the simultaneous dysregulation of a large number of pathways, and even in the same biopsy cells may come with a diverse set of mutational landscapes, due to cancer heterogeneity. Additionally, due to the bulk approach, shifts in cell type composition (due to cancer, aging, or inflammation) are picked up in the gene count matrices, acting as a confounding variable.

Finally, the STRING channel leverages phylogeny, enabling to detect functions that may not be apparent from a merely phenotypic characterization; although not unprecedented, phylogenetic data have been relatively neglected in recent architectures, in favor of catch-all (heavily literature-biased) PPIs. Conversely, the rationale of phylogeny ultimately remains a form of guilt by association, even though literature-unbiased.

In conclusion, in this work we introduced GhostBuster, a novel machine learning framework designed to prioritize gene functions, disease associations, and interaction modalities in a literature-unbiased manner. Among the data channels evaluated, TCGA emerged as the most promising, offering minimal literature bias, while still maintaining robust predictive performance—comparable to, or only marginally below, that of heavily literature-biased sources like Gene Ontology. These findings support the adoption of TCGA as the default data channel for GhostBuster and lay the groundwork for scaling the platform into a publicly accessible resource. We envision GhostBuster evolving into the first online repository of literature-unbiased gene prioritizations, enabling researchers to identify and explore “ghost genes” relevant to their specific domains. This repository could be structured into three user-driven channels: (i) physiology-based, for querying genes potentially involved in specific cellular functions or compartments; (ii) disease-based, for identifying novel contributors to complex, polygenic conditions such as neurodegeneration, cancer, and autoimmune disorders; and (iii) gene-centric, for exploring granular interaction networks of individual genes. While experimental validation remains a critical next step, GhostBuster represents a foundational advance in unbiased gene discovery. By enabling systematic exploration of under-characterized genes, it offers a powerful tool for accelerating functional genomics and, ultimately, therapeutic innovation.

## Materials and methods

### Virtual Lab

GhostBuster was created using our Virtual Lab (VLab) ML framework.^69^ It was developed in Python 3.9.19, using the Anaconda v23.7.4 distribution to manage libraries and environments, and the PyCharm 2022.2.5 (Professional Edition) environment. The ML code relied on the PyTorch (`torch`) and Scikit-learn (`sklearn`) frameworks. All training was conducted using Nvidia CUDA^126^ v12.3.2 to leverage GPU acceleration for computational efficiency.

### LINCS dataset

Library of Integrated Network-Based Cellular Signatures (LINCS) 2020 was accessed through the Clue.io data portal of the Broad Institute of MIT and Harvard.^78,79^ Specifically, level 5 signatures were manually downloaded for all perturbation types (`trt_cp`, `trt_misc`, `trt_oe`, `trt_sh`, `trt_xpr` files), alongside their metadata. A total of 3,026,460 transcriptomic measurements and 1,201,944 signatures were retrieved. Next, signatures were filtered according to the quality control metrics included in the signature information, specifically, only those marked as high quality (`is_hiq` column) were passed; this step brought the number of signatures down to 282,990. Next, namespace conversions were applied. Reads were originally expressed as 12,328 Entrez IDs; these were updated by replacing obsolete IDs with the current ones, and mapped to the namespace of the NCBI Gene repository; 3 Entrez IDs were dropped. Finally, data were aggregated into an object of the `MultiOmicDataset` class from our VLab library, consisting of only 1 omic (Tx), with 12,325 reads by 282,990 samples.

### TCGA dataset

The Cancer Genome Atlas (TCGA)^127^ was employed, by aggregating together all RNA-sequencing cases available in the repository. The dataset was accessed through the NIH GDC data portal, version 2.^128^ Files were filtered by the “experimental strategy” field, selecting “RNA-Seq”; and by the “Data Type” field, selecting “Gene Expression Quantification”. After applying the file selection, a File manifest, Biospecimens manifest, and a Clinical manifest were generated. The number of retrieved samples was 25,259 (November 2024). A dataframe was assembled from the downloaded files: the `tpm_unstranded` column was used, in line with established transcriptomics guidelines, suggesting to utilize transcripts per million mapped reads, TPM.^129,130^ Next, namespace conversions were applied: RNAseq data were originally expressed as 60,660 Ensembl IDs; these were demultiplexed into 39,467 Entrez IDs; synonymous IDs were aggregated by sum; all-0 reads were dropped, resulting in 36,027 approved reads. Finally, data were aggregated into an object of the `MultiOmicDataset` class from our VLab library, consisting of only 1 omic (Tx), with 36,027 reads by 25,259 samples.

We applied pre-processing to make data distribution more suitable for ML processing. After mapping genes into the Entrez namespace, we re-applied the normalization by transcripts per million mapped reads (TPM), as recommended by Tx guidelines.^129,130^ Next, we log-transformed the data to ensure that all reads would occupy the same order of magnitude.

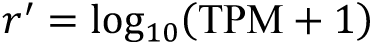

### STRING dataset

Search Tool for the Retrieval of Interacting Genes/Proteins (STRING) was accessed through the repository’s official website (www.stringdb-downloads.org).^76^ The file corresponding to *H. sapiens* (“9606.protein.links.detailed.v12.0.txt.gz”) was manually downloaded from their FTP server. It was expressed in the form of a graph incidence matrix, listing 13,715,404 protein-protein relations in terms of 8 scores: `neighborhood`, `fusion`, `cooccurence`, `coexpression`, `experimental`, `database`, `textmining` and `combined_score`. Only `neighborhood`, `fusion`, `cooccurrence` and `coexpression` were utilized for the sake of training GhostBuster, the others were dropped. Next, namespace conversion was applied. Data were originally expressed as 19,622 Ensembl protein IDs; these were demultiplexed into 19,199 Entrez IDs; synonymous IDs were aggregated by max; this reduced the number of edges to 6,707,960. Next, the incidence matrix was converted into a pseudo-adjacency matrix. First, all relations were duplicated with the two genes (i.e., the departing node and the destination node) swapped, as STRING edges are meant to be non-directional; this increased the number of edges by a factor of 2×. Next, each destination node was multiplexed into 4 “pseudo”-interactors, one for each of the criteria of interest, namely `neighborhood`, `fusion`, `cooccurrence` and `coexpression`. Next, a dataframe was created, connecting each of the selected 19,093 Entrez IDs to the 19,093 × 4 = 76,372 pseudo-interactors, whereby a given cell in the dataframe would report the interaction score between that row’s gene and that column’s gene, according to that column’s criterion. Next, all-0 rows and columns were dropped. Finally, data were aggregated into an object of the `MultiOmicDataset` class from our VLab library, consisting of only 1 omic (phylogenomics), with 18,845 reads (the origin nodes) by 49,402 interactor proteins (the receiving nodes-criterion pairs, playing the role of “samples”).

### GO dataset

Gene Ontology was accessed through the official website of the repository (www.geneontology.org). The files corresponding to *H. sapiens* (namely, “goa_human.gaf.gz”, “goa_human_complex.gaf.gz”, “goa_human_isoform.gaf.gz”, “goa_human_rna.gaf.gz”) were manually downloaded from their FTP server, alongside the GO tree structure file (“go-basic.obo”). The tree structure came in the form of an incidence matrix, associating each GO term with other GO terms, while specifying the nature of such relation (`is_a`, `regulates`, `positively_regulates`, `negatively_regulates`, `part_of`); specifically, the two columns `is_a` and `part_of` are guaranteed to form a directed acyclic graph^131^; they were the only columns to be utilized for the sake of this dataset, while the other relation types were dropped. The human annotation came in the form of an incidence matrix, associating human genes or proteins and GO terms, with additional metadata such as evidence type and source; specifically, a total of 876,365 relations were listed, involving 812,829 proteins (expressed as Uniprot^132^ IDs), 54,821 RNA genes (expressed as RNAcentral^133^ IDs) and 8,725 complexes (expressed as ComplexPortal^70^ IDs; as the current implementation of GhostBuster focuses on protein-coding genes, only proteins were utilized, while the other entries were dropped. All relation types were included, except those marked as “negative”, as they were too few to be utilized as negative examples for ML (0.17%).

Surprisingly, we found that, in the original form of GO, a gene’s membership in a given class did not necessarily imply its membership in the related superclasses—despite GO being described as a hierarchical tree. For example, term `GO:0007127` “meiosis I” is a descendent class of `GO:0022414` “reproductive process”, but the former includes 21 members (either proteins, RNAs or complexes), while the latter includes only 8, none of which are shared between the two classes. To compensate for this mismatch, GO was “enriched” to guarantee that all member-class pairs would be reflected across all the upstream superclasses, by applying matrix multiplication between the annotation adjacency matrix and the tree structure adjacency matrix. The enrichment increased the number of member-class relations from 474,446 to 3,456,842 (+628.61%).

Next, namespace conversion was applied. Data were originally expressed as 85,087 Ensembl protein IDs; these were demultiplexed into 19,237 Entrez IDs; synonymous IDs were aggregated by max; this reduced the number of edges to 1,731,514. Finally, for the sake of training GhostBuster, the incidence matrix was converted into a Boolean adjacency matrix; then, it was converted into an object of the `MultiOmicDataset` class from our VLab library, consisting of only 1 “pseudo-omic” (GO membership), with 19,237 reads (genes) by 17,873 GO terms (playing the role of “samples”) and a total of 1,731,514 positive values. Of the 19,237 genes, 12,567 were biological processes (BP), 1,760 were cell components (CC) and 3,546 were molecular functions (MF). Conversely, for the sake of using GO classes as a source for target gene lists, the incidence matrix was converted into an object of the `OntologyDataset` class from our VLab library, with identical properties.

### Tabular GhostBuster target datasets

In Tabular GhostBuster, a total of 81 gene lists were used as training targets, which included 54 classes from GO,^41^ 6 classes from the Phenotype-Genotype Integrator (PheGenI)^21^ and 21 classes from the Molecular Signatures Database (MSigDB).^18^ The latter were, in turn, originally provided by different repositories, namely: 8 classes from the Kyoto Encyclopaedia of Genes and Genomes (KEGG) pathways,^16^ 12 classes from the Human Phenotype Ontology (HPO),^134^ while 1 class included genes from HPO, Biocarta^30^ and MSigDB itself. Their details can be found in Supplementary Table 1. The GO classes were created by combining one or more membership lists from the GO dataset described in the previous paragraph.

MSigDB pathways were accessed through the Gene Set Enrichment Analysis (GSEA) repository of the University of California San Diego and the Broad Institute of MIT and Harvard (www.gsea-msigdb.org).^135,136^ The entire list of 68 human collections was downloaded, which included, among other things, disease-relevant KEGG, HPO and Biocarta pathway member lists. No namespace conversions were necessary, as the member genes were already expressed in terms of Entrez IDs. A total of 4,058,136 gene-phenotype relations were retrieved, all of which were mapped in the NCBI Gene namespace. Data were aggregated into an object of the `OntologyDataset` class from our VLab library, consisting of 34,837 classes, 42,494 member genes and 4,058,136 gene-class relations.

PheGeni was accessed through the National Center for Biotechnology (NCBI) website (www.ncbi.nlm.nih.gov/gap/phegeni). The “PheGenI_Association_full.tab” file was downloaded from their FTP server. No namespace conversions were necessary, as the member genes were already expressed in terms of Entrez IDs. A total of 106,464 gene-phenotype relations were retrieved; both intragenic and intergenic hits were included; for the latter, the nearest genes were used as a first approximation. Next, gene-phenotype relations were filtered with respect to p-value, selecting for *p* < 10^−8^, thus reducing the number of gene-phenotype relations to 17,281; of these, 100 were dropped as their genes did not appear in the NCBI Gene namespace. Data were aggregated into an object of the `OntologyDataset` class from our VLab library, consisting of 789 classes (diseases), 7,122 member genes and 17,187 gene-class relations.

### Graph GhostBuster target datasets

In Graph GhostBuster, a total of 21 gene-gene interaction modalities were used as independent training targets, for a graph edge prediction task. These edges were sourced from the Biology Mega Graph. Their details are depicted in Supplementary Table 2. 11 relational datasets, encompassing gene-gene or miRNA-gene relations, were downloaded from their respective sources and parsed, namely: ComplexPortal,^70^ human DEPhOsphorylation Database (DEPOD),^71^ GO,^6^ MicroT (MirBase and MirGeneDB versions),^72^ Mirbase,^73^ NCBI Gene,^74^ PhosphoSitePlus,^75^ Signor,^36^ STRING^76^ and TarBase.^77^ GO was parsed as described above. Relation labels underwent manual standardization and curation, to make them cross-compatible across the different data sources. All genes were converted from their original namespaces into the NCBI Entrez namespace, and miRNA names into the miRBase namespace^73^; unmatched names were dropped; synonymous entries were aggregated by a logical *OR*. The edge labels were harmonized into a common, fixed vocabulary. The final result was a knowledge graph encompassing 322,255,334 known regulatory relations, among 193,384 gene Entrez IDs and 3,537 miRNAs.

### Encoder-decoder model

For all input datasets, and for all target classes, an MLP model (encoder) was trained on a binary classification task, to predict the gene membership status from the input dataset, treating the members of the target class as positive examples (1.0) and the non-members as negative examples (0.0). Genes (examples) were split into 10 cross-validation (CV) folds, each composed of 80% training examples, 10% validation examples and 10% test examples; splitting was conducted in a stratified way, with respect to the target class membership status. The MLP was instantiated from the PyTorch `Sequential` class; it was trained using the `AdamW` optimizer, and the `BCEWithLogitsLoss` loss (sigmoid layer followed by binary cross-entropy). Due to the highly imbalanced target, class weights were applied, which were inversely proportional to their respective abundances. To prevent overfitting, early stopping was employed, based on the Matthew’s correlation coefficient (MCC) in the validation dataset. At each prediction, grid search was automatically conducted by VLab to select the best-performing hyperparameters in the validation set. At the end of the training, latent representations (embeddings) for all genes were generated by feeding the input data into the MLP, and collecting the activation values of the neurons in the second-last layer. Embedding generation was conducted independently for each CV fold. Of note, GhostBuster automatically excludes all input dataset columns that are either identical, a subclass or a superclass to the target one. This is crucial to prevent data leakage, for example, when GhostBuster is trained using the GO channel, while targeting a gene list that is also sourced from GO.

Next, a support vector machine (SVM) binary model (decoder) was trained on a binary classification task, to predict the gene membership status from the embeddings, again, treating the members of the target class as positive examples (1.0) and the non-members as negative examples (0.0). The same CV folds were utilized as for the training of the encoder. The SVM was instantiated from the Scikit-learn `SVC` class. At each prediction, grid search was automatically conducted by VLab to select the best-performing hyperparameters in the validation set; this included retroactively selecting the best hyperparameters in the encoder, that would yield the best performance in the encoder—provided that the CV splits were identical between the two of them. At the end of the training, margins were calculated for each gene, expressing the geometric distance from the hyperplane boundary between the positive and the negative examples.

For each grid search epoch, two sets of margins were generated: (1) a test dataset version, which is a concatenation of the model’s output corresponding to the 10% test dataset examples, repeated across the 10 CV folds, so as to cover the entirety of the dataset. Because margins are not directly comparable from one CV fold to another, they were transformed into ranks within each CV fold; as a result, the test dataset margins would include 10 genes ranked as no. 1; 10 genes ranked as no. 2, etc. (2) A full dataset version, which was created by averaging the margins of all genes, across the 10 CV folds; as a result, each gene yielded an average value across 10 predictions: 8 of these predictions were in its capacity of a training dataset example; 1 in its capacity of a validation dataset example; 1 in its capacity of a test dataset example.

### Graph convolutional neural network

For all input datasets, and for all interaction modalities, a graph convolutional neural network (GCN) was trained to predict the known gene-gene relations from the input dataset, fed as node features. The graph was represented as a PyTorch Geometric `Data` object. The known gene-gene relations (edged) served as the positive examples; negative examples were randomly generated, in a 50:1 ratio to the positive ones. The positive and the negative examples were randomly split into a training dataset (80%), a validation dataset (10%) and a test dataset (10%), in a stratified way. The model was instantiated from the PyTorch Geometric `GAE` (graph autoencoder) class, with the encoder derived from the `GCNEncoder` class; it was trained in an edge prediction setting, using the `AdamW` optimizer. Due to the highly imbalanced edge prediction task, class weights were applied, which were inversely proportional to their respective abundances; to achieve this, a manually defined loss was employed, which corresponded to the weighted sum of the correctly classified positive edges and the correctly classified negative edges. To prevent overfitting, early stopping was employed, based on the Matthew correlation coefficient (MCC) in the validation dataset. At each prediction, grid search was automatically conducted by VLab to select the best-performing hyperparameters in the validation set. By the end of the training, latent representations (embeddings) for all genes were learned. By subjecting the embeddings to matrix multiplication and sigmoid transformation, a predicted adjacency matrix was generated for the entire graph.

Some of the interaction modalities came with a cognate antonymic modality (see Supplementary Table 2); for example, if gene *g*_1_ “upregulates quantity” of gene *g*_2_, one could assume the opposite to be false, namely that *g*_1_ “downregulates quantity” of gene *g*_2_ is false; however, this assumption is not always verified, as interactor genes may sometimes yield opposed effects based on the cell type. For the cases where a cognate antonymic modality was available, two models were trained independently:

- R model (random negatives), where the negative edges used in training are the randomly generated ones.
- C model (curated negatives), where the negative edges used in training are the curated ones, that is, the known annotations from the antonymic modality.

In both models, no matter whether R or C, ML metrics were independently measured both with respect to the random negatives, and with respect to the curated negatives.

### Protein-Protein Interaction score

One of the control groups of GhostBuster consisted in the protein-protein interaction (PPI) score, which quantified the total number of interactors that a given gene has, that are also members of the target list. STRING was downloaded as described in the previous paragraph. Yet, this time, it was not filtered for any specific column, rather, it was applied “off the shelf”, to simulate its classic use case in the guilt by association paradigm. For each gene-target list pair, the incidence matrix (consisting of 6,707,960 Entrez edges) was filtered for entries that connected either the provided gene with any member of the target list, or vice versa; next, the number of unique in-list partners was counted, and returned as the PPI score for that gene-target list pair. No ML was involved in computing the PPI score, other than the ML that had been used to create the STRING dataset in the first place, by its authors.^59^

### PubMed score and richness score

Another control group of GhostBuster consisted of two metrics that express how thoroughly annotated a gene is, i.e., where it is positioned along the “ghostliness” spectrum. These metrics are:

- Richness score: it expresses the number of annotations in the GO and MSigDB datasets, combined. The GO adjacency matrix was prepared as explained above; likewise, the MSigDB adjacency matrix was prepared as explained above; the two matrices were concatenated; the number of edges (i.e., connections between a gene and a GO/MSigDB class) was counted for every gene; to improve interpretability, the scores were normalized by dividing them by the maximum (achieved by gene *TMEM207*, with 7,201 annotations) and multiplying them by 1000; as a result, the scores are expressed in a floating-point numeric scale from 0.0 to 1000.0, whereby 0.0 indicates the lack of annotations, and 1000.0 is achieved only by *TMEM207*.
- PubMed score: it simply indicates the number of publications about a given gene in the PubMed repository. Because this metric was highly interpretable *per se*, it was not further normalized; as a result, the scores are expressed in an integer scale from 0 to 11,581, whereby 0 indicates the lack of publications, and the maximum value of 11,581 is only achieved by gene *TP53*.

### GPT score

One last control group of GhostBuster consisted in the Generative Pre-trained Transformer (GPT) score, that is, a standardized quantification of the association degree between a given gene and a “phenotype” of interest— either a cell physiological function, a cell organelle or a disease. Scoring was performed by OpenAI GPT-4o mini.^83^ For every phenotype, a script interrogated the OpenAI application programming interface (API) by submitting a prompt and a batch of 200 genes, while expecting a structured output made up of a score and a brief motivation. For the minority of genes where OpenAI reported null output, the query was recursively repeated until an output was achieved. The prompt (Supplementary Table 3a) was designed to guarantee objectiveness and reproducibility; the GPT-aided prompt engineering tool on the GPT playground application (platform.openai.com/playground/) was used to optimize the prompt text for this purpose. A total of 2,722,950 phenotype-gene pairs were submitted to GPT-4o mini across ∼11,000 requests, spending ∼86 million input tokens, for a total cost of $12.53 in input, and $70.39 in output.

### GPT validation of recently discovered genes

To validate GhostBuster’s predictions, GPT 4o (full version)^137^ was tasked to make a list of recently discovered genes, for each of the phenotypes of interest. A script was created that, for every phenotype, interrogated the OpenAI API by submitting the phenotype name and a brief description, while expecting a structured output made up of gene identifiers and the literature sources that justified such decision. Again, the prompt (Supplementary Table 3b) was designed to guarantee objectiveness and reproducibility; the GPT-aided prompt engineering tool on the GPT playground application^138^ was utilized to optimize the prompt text to this end. Tokens and spend was negligible compared to those associated with calculating GPT scores.

### Hardware

All development was conducted on a high-performance mainframe computer, featuring Nvidia 4090 GPU and Intel i9-14900K 3.20 GHz CPU.

## Data availability

The VLab library^69^ source code, including the full GhostBuster module, can be found in the following repository: https://github.com/giuliodeangeli/vlab

## Statements

### TCGA/CPTAC studies

The results shown here are in whole or part based upon data generated by the TCGA Research Network: https://www.cancer.gov/tcga.

Data used in this publication were generated in part by the Clinical Proteomic Tumor Analysis Consortium (CPTAC).

### ROSMAP study

The results published here are in whole or in part based on data obtained from the AD Knowledge Portal (https://adknowledgeportal.org).

Study data were provided by the Rush Alzheimer’s Disease Center, Rush University Medical Center, Chicago. Data collection was supported through funding by NIA grants P30AG10161 (ROS), R01AG15819 (ROSMAP; genomics and RNAseq), R01AG17917 (MAP), R01AG30146, R01AG36042 (5hC methylation, ATACseq), RC2AG036547 (H3K9Ac), R01AG36836 (RNAseq), R01AG48015 (monocyte RNAseq) RF1AG57473 (single nucleus RNAseq), U01AG32984 (genomic and whole exome sequencing), U01AG46152 (ROSMAP AMP-AD, targeted proteomics), U01AG46161(TMT proteomics), U01AG61356 (whole genome sequencing, targeted proteomics, ROSMAP AMP-AD), the Illinois Department of Public Health (ROSMAP), and the Translational Genomics Research Institute (genomic).

TMT Proteomics: Study data were provided through the Accelerating Medicine Partnership for AD (U01AG046161 and U01AG061357) based on samples provided by the Rush Alzheimer’s Disease Center, Rush University Medical Center, Chicago. Data collection was supported through funding by NIA grants P30AG10161, R01AG15819, R01AG17917, R01AG30146, R01AG36836, U01AG32984, U01AG46152, the Illinois Department of Public Health, and the Translational Genomics Research Institute.

RNA-seq Bulk Brain: “Annie J. Lee, Yiyi Ma, Lei Yu, Robert J. Dawe, Cristin McCabe, Konstantinos Arfanakis, Richard Mayeux, David A. Bennett, Hans-Ulrich Klein, and Philip L. De Jager. Multi-region brain transcriptomes uncover two subtypes of aging individuals with differences in Alzheimer’s disease risk and the impact of APOEε4. bioRxiv 2021”

### MSBB study

The results published here are in whole or in part based on data obtained from the AD Knowledge Portal (https://adknowledgeportal.org/). These data were generated from postmortem brain tissue collected through the Mount Sinai VA Medical Center Brain Bank and were provided by Dr. Eric Schadt from Mount Sinai School of Medicine.

TMT Proteomics: The results published here are in whole or in part based on data obtained from the AD Knowledge Portal (https://adknowledgeportal.org/). These data were provided by Dr. Levey from Emory University based on postmortem brain tissue collected through the Mount Sinai VA Medical Center Brain Bank provided by Dr. Eric Schadt from Mount Sinai School of Medicine.

### ROSMAP-IN

The results published here are in whole or in part based on data obtained from the AD Knowledge Portal (https://adknowledgeportal.org). Study data were provided by the Young-Pearse Lab of the Ann Romney Center for Neurologic Disease at Brigham and Women’s Hospital (BWH), Boston. Stem cell lines used in this study were generated through collaboration between BWH, the New York Stem Cell Foundation, and the Rush Alzheimer’s Disease Center, Rush University Medical Center, Chicago.

## Acknowledgements

The authors would like to express their sincere gratitude to all those who provided invaluable advice, insightful discussions, and constructive feedback throughout this project. Their expertise and guidance greatly contributed to the development of this work. In particular, we acknowledge the contributions of Cristiano Peron, Antonio Cisternino, Pietro Barbiero, Niccolò Pancino, Michelangelo Diligenti, Lorenzo Giusti, Donato Crisostomi, Giuseppe Marra, Gabriele Ciravegna, Francesco Giannini, Aviva Tolkovski and Giorgio Vivacqua.

This research was supported by funding from the Cambridge Trust and the Medical Research Council Doctoral Training Partnership (MRC DTP). We acknowledge their generous support in facilitating this study.

## Author contributions

G.D. was responsible for dataset preparation, machine learning model development and training, explainability analysis, literature review, manuscript writing.

M.G.S. and P.L. provided conceptual guidance and contributed to manuscript revisions.

## Supplementary Figures

**Supplementary Figure 1.**
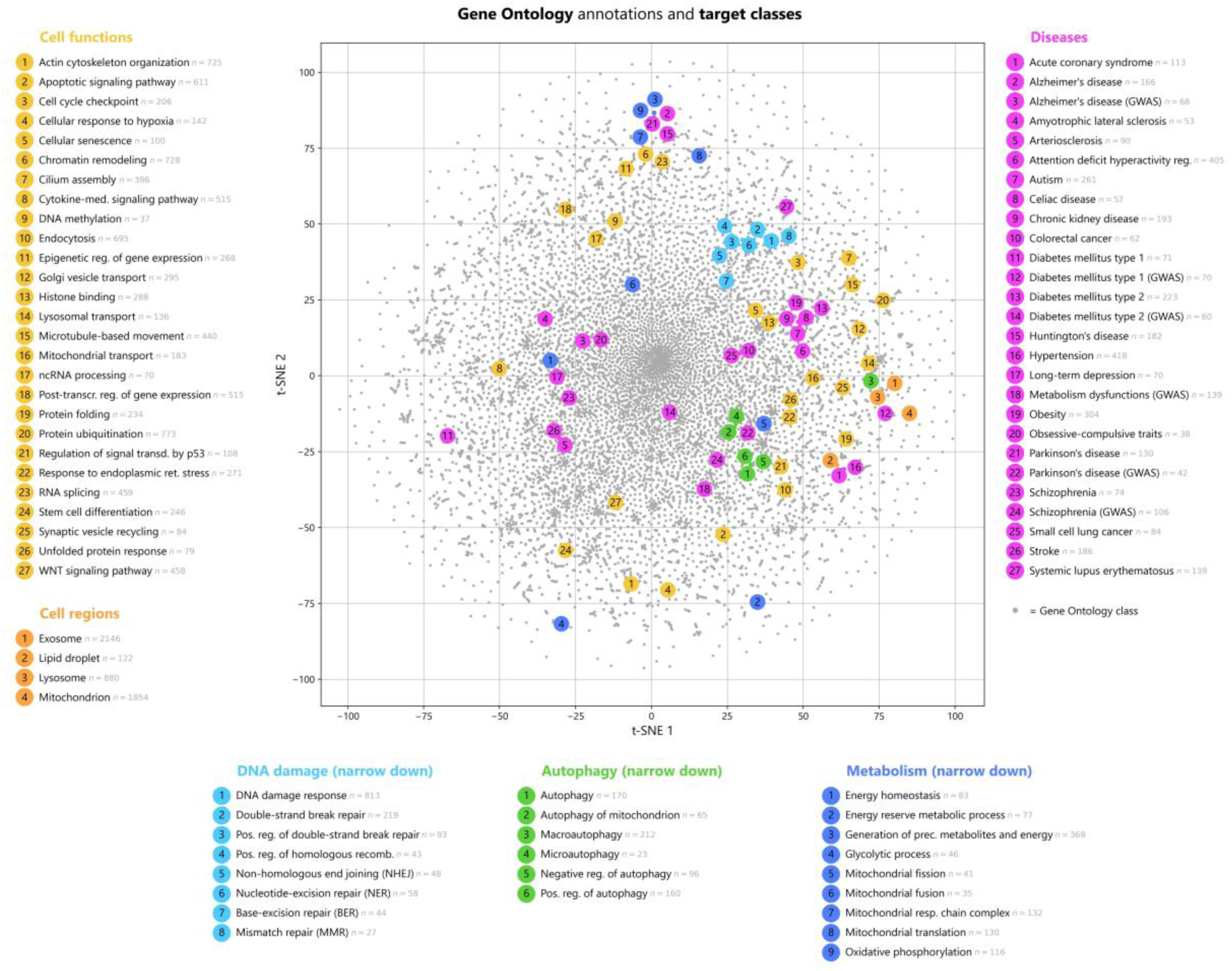
t-SNE plot of the target classes used in Tabular GhostBuster experiments. Tabular GhostBuster provides a literature-unbiased prediction of which genes are more likely to belong to a certain class, on the basis of a list of genes that are already known to belong to such class (target list). To validate its efficacy, a selection of 81 target lists of genes was used, namely 27 physiological cell functions and 4 cell regions sourced from Gene Ontology, 21 disease pathways sourced from the Molecular Signatures Database (MSigDB), and 6 hit lists from genome-wide association studies (GWAS) sourced from the Phenotype-Genotype Integrator (PheGenI). These 81 target lists of genes were organized into 3 general-purpose groups: 27 physiological cell functions (yellow), 4 cell regions or organelles (orange), 27 diseases (pink); and three narrow-down groups, meant to demonstrate how GhostBuster can provide a deeper focus into the submechanisms within a cell function of interest: 8 for DNA damage response (cyan), 6 for autophagy (green) and 9 for metabolism (blue). Note: in disease, classes are meant as MSigDB pathway memberships, unless “GWAS” is explicitly specified. This is a class-centric t-distributed stochastic neighbor embedding (t-SNE) plot, depicting the relative location of the 81 target lists (colored, numbered circles) on the background of all GO classes (grey dots). It is easy to note that gene classes that are biologically close (e.g., the three narrow-down groups) also appear close in the t-SNE space, confirming that the plot is effectively capturing real biological signal. The plot confirms that the 81 target lists are sampling across the entirety of cellular biology, in an unbiased way. It is remarkable to observe, in the disease-related classes (e.g., Parkinson’s), how distant the MSigDB membership lists localize from their cognate GWAS classes; the former are dominated by high-penetrance familial mutations, while the latter are low-effect size “risk factors”, which are generally enriched in the sporadic forms of the disease. They appear completely divergent from a biological standpoint, *as if* the familial and sporadic forms had completely different biologies.

**Supplementary Figure 2.**
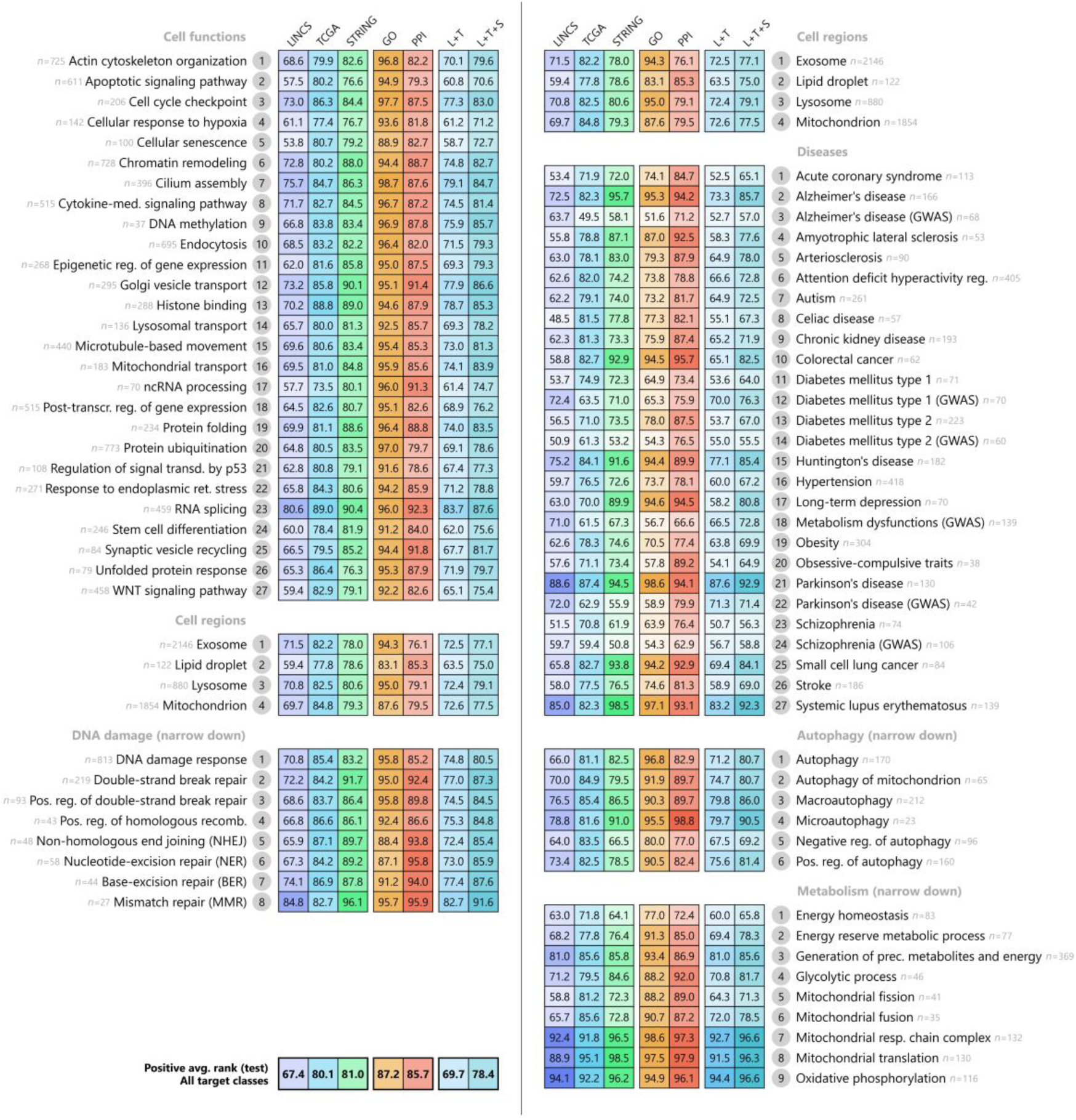
Tabular GhostBuster performance on the 81 target lists, measured by average rank of the positive class members, in the test dataset prediction. The plot measures GhostBuster’s performance in terms of the average rank of the positive examples within the output prioritization hierarchy, in the test dataset prediction. Column by column, first, it displays GhostBuster’s performance when trained on the three “unbiased” data sources, namely LINCS (blue), TCGA (cyan) and STRING (green). Next, the performance when trained on the “biased” data source, namely GO (orange). Next, for comparison, the performance of a simple GBA-type gene prioritization script, based on the physical protein-protein interactions in STRING (red). Next, the combined performance obtained by averaging out the predictions from LINCS and TCGA (L+T), and from LINCS, TCGA and STRING (L+T+S). The phenotype labels are displayed next to each line, accompanied by the numerosity of the corresponding target gene lists (*n*). The average performance is displayed in the bottom left corner. All cells are color-coded based on the performance score, in a non-linear fashion. Overall, GO achieves the best performance, followed by PPI, STRING, TCGA and finally LINCS. This distribution accurately reflects the amount of literature bias that is present in each of the 5 sources.

**Supplementary Figure 3.**
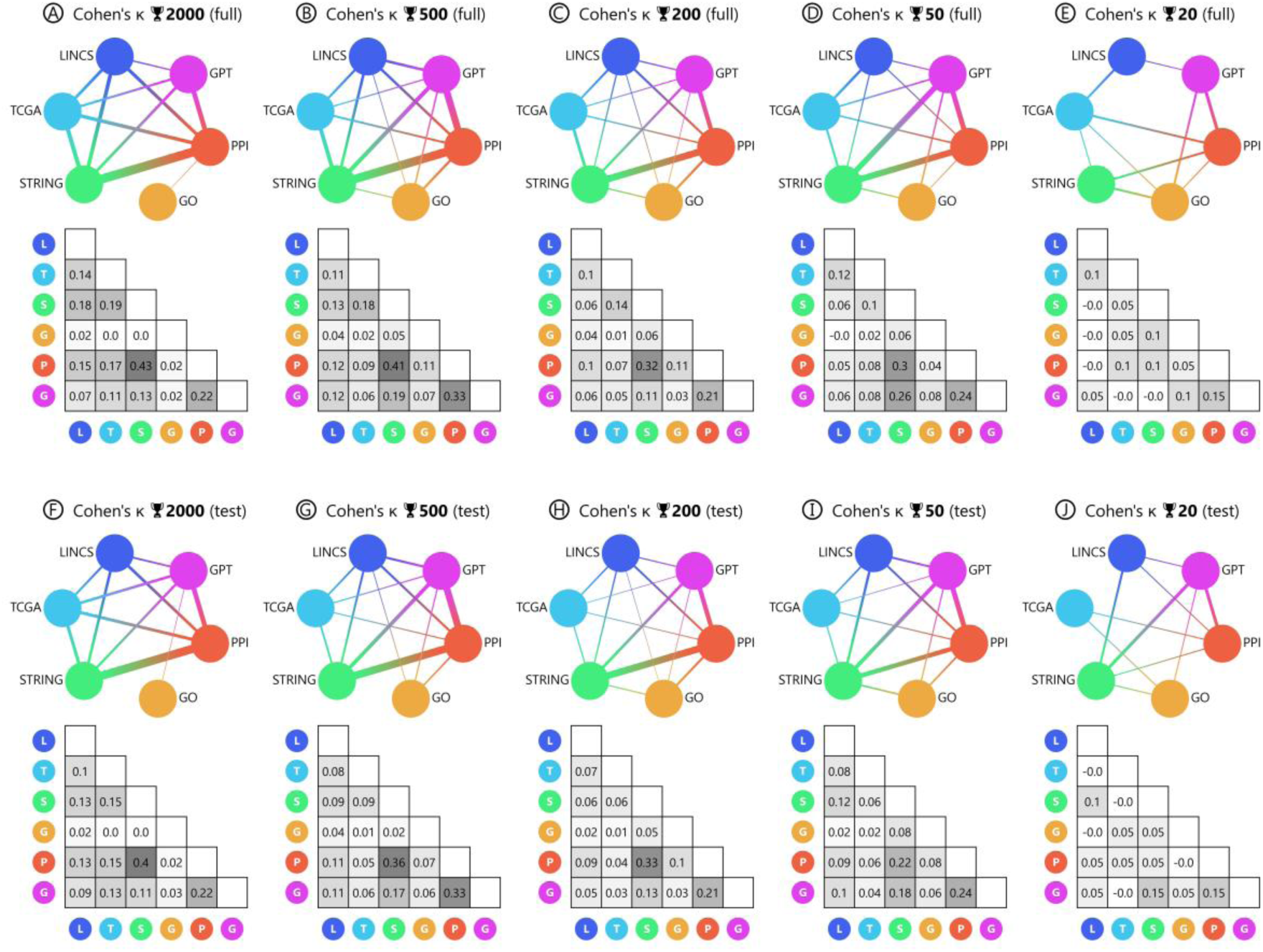
Correlation between the top-ranked, non-target genes generated by GhostBuster from different data sources, measured by Cohen’s *k*. The plot shows the correlation in the top-ranked gene lists (once removed the target ones), obtained from different data channels in the test dataset prediction: GhostBuster trained on the three “unbiased” data sources, namely LINCS (blue), TCGA (cyan) and STRING (green); next, GhostBuster trained on the “biased” data source, namely GO (orange); next, a simple GBA-type gene prioritization script, based on the physical protein-protein interactions in STRING (red); next, the GPT score (pink). The analysis was carried out on the top *n* genes (🏆 *n*), starting from 2,000 and moving up to the top 20. The analysis was repeated on the full dataset prediction (upper row) and on the test dataset prediction (lower row). Each iteration of the analysis is displayed as a chord diagram, with connection thickness proportional to the Cohen’s ***κ*** between the corresponding data channels; and as a table, displaying the Cohen ***κ*** numerical values.

**Supplementary Figure 4.**
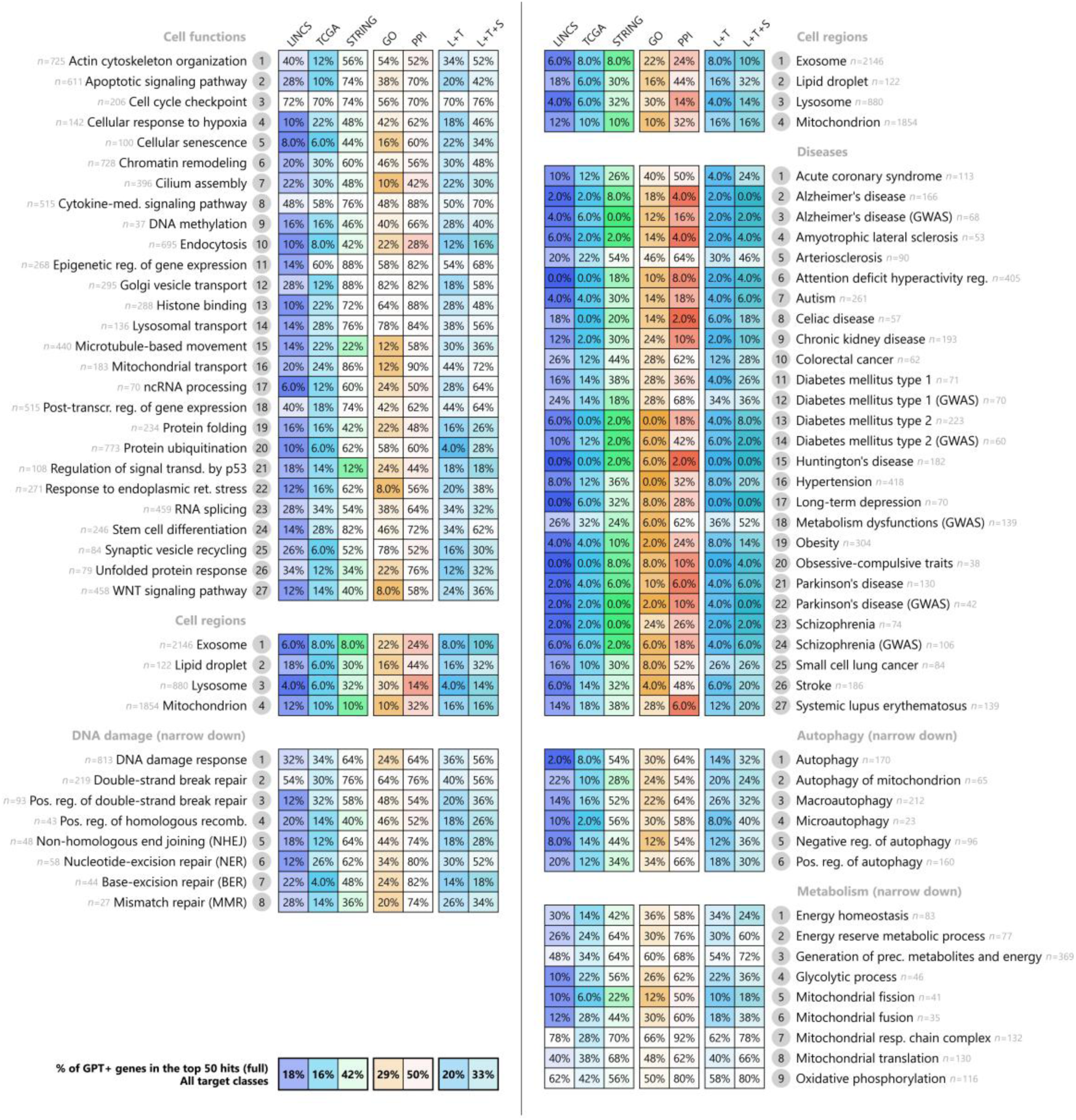
Tabular GhostBuster performance on the 81 target lists, measured by the percentage of GPT-positive genes in the top 50 hits, in the full dataset prediction. The plot measures GhostBuster’s performance in terms of the percentage of GPT-positive genes in the top 50 hits, in the full dataset prediction. GPT-positivity indicates that a gene achieved a score of 6/10 or higher for that target gene class, in the GPT control group of GhostBuster. The latter interrogates the OpenAI API, and specifically the GPT-4o mini^83^ model, to quantify the association strength between a gene and a given physiological function or disease, according to current scientific literature. Column by column, first, the plot displays GhostBuster’s performance when trained on the three “unbiased” data sources, namely LINCS (blue), TCGA (cyan) and STRING (green). Next, the performance when trained on the “biased” data source, namely GO (orange). Next, the performance of a simple “guilt by association”-type gene prioritization script, based on the physical protein-protein interactions in STRING (red). Next, the combined performance obtained by averaging out the predictions from LINCS and TCGA (L+T), and from LINCS, TCGA and STRING (L+T+S). The phenotype labels are displayed next to each line, accompanied by the numerosity of the corresponding target gene lists (*n*). The average performance is displayed in the bottom left corner. All cells are color-coded based on the performance score, in a non-linear fashion. Overall, we can conclude that TCGA achieves the highest content of GPT-negative genes in its prediction, followed by LINCS, GO, STRING and PPI. Such a high score for LINCS is not particularly surprising due to its generally poor machine learning (ML) performance. Conversely, TCGA’s performance was only slightly inferior to that of the biased sources (see Figure 1 and Supplementary Figure 2), suggesting that the TCGA channel achieves a robust predictive power, while also minimizing literature bias.

**Supplementary Figure 5.**
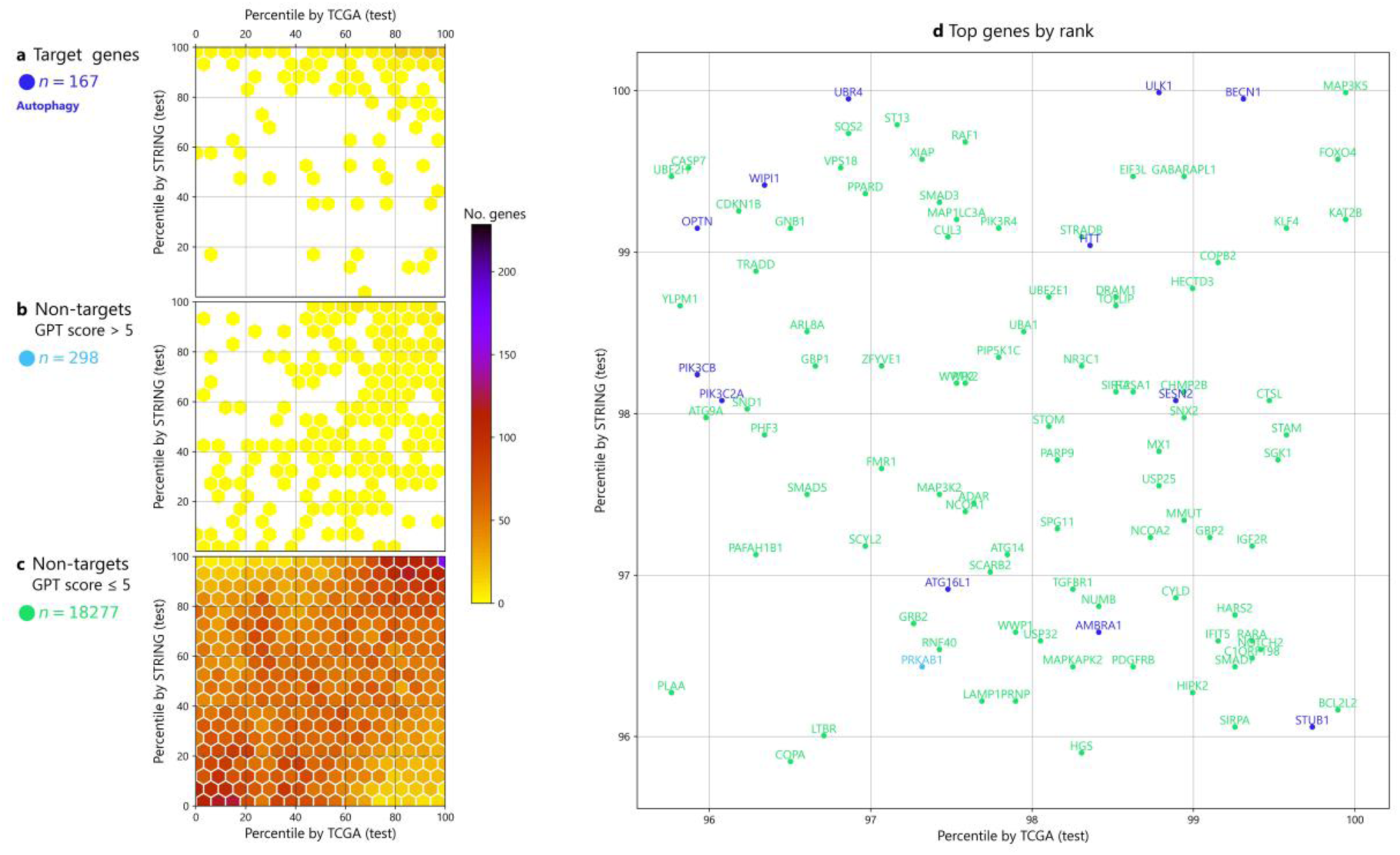
Prediction output of Tabular GhostBuster on autophagy, cross-matching the TCGA and STRING unbiased channels, in the test dataset prediction. Tabular GhostBuster was trained to recognize 167 genes in the autophagy list from Gene Ontology (GO), utilizing the TCGA channel or the STRING channel as its training input. At the end of the training, a gene prioritization list was assembled by concatenating the rank-transformed test dataset predictions, for both models. Additionally, GhostBuster’s GPT control group was utilized to obtain GPT-4o mini-derived association scores between each gene and autophagy, in a standardized scale from 1 to 10 (color code). Finally, genes were organized in a scatter plot based on their rank in the TCGA channel (x-axis), versus their rank in the STRING channel (y-axis). (Left) Density plot, decomposed in three parts: (a) the 167 target genes (blue genes), (b) the non-target genes with a GPT score of 6 or higher (cyan genes), and (c) the non-target gene with a GPT score of 5 or less (green genes). In all three plots, agreement between the two channels can be observed in the top hits (top-right corner). (Right) Scatter plot detail of the top 4 percentiles, at gene-level resolution. The green genes depicted here were simultaneously selected by both the TCGA and the STRING channel as top hits in the test dataset, despite their involvement in autophagy was largely unexplored in the scientific literature: this makes them the “interesting” hits for the sake of our prediction.

**Supplementary Figure 6.**
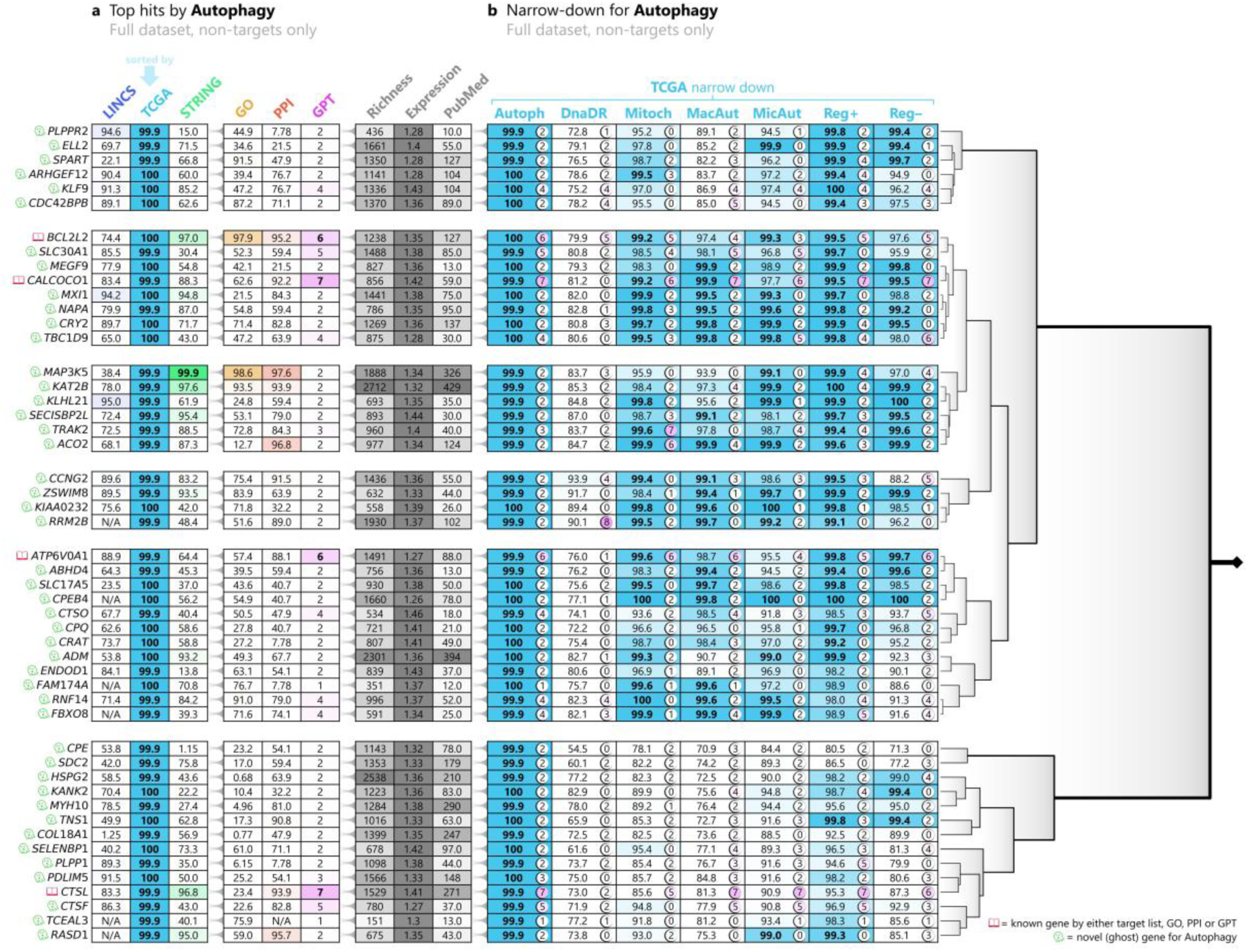
Tabular GhostBuster narrow-down plot across 7 sub-mechanisms of autophagy. Tabular GhostBuster was trained to recognize 170 genes in the “autophagy” list from Gene Ontology (GO), utilizing the TCGA channel as its training inputs (cyan arrow). At the end of the training, the top 50 non-target genes were selected, based on the prioritization list generated by such model in the full dataset, and are depicted here. The left part of the table showcases the percentile ranks generated by training Tabular GhostBuster against the “autophagy” gene list, based on the LINCS (blue), TCGA (cyan), STRING (green) and GO (yellow) channels; next, GhostBuster’s auxiliary PPI scores (red), GPT scores (pink); and the richness, average expression and PubMed scores (grey). A crimson book icon on the left side of genes’ symbols is assigned for genes that are already known for playing a role in the phenotype of interest, either because they are target list members, top-rankers in the GO model (>99^th^ percentile), top rankers in the PPI score (>99^th^ percentile), or GPT-positive (GPT score > 5). Conversely, a green ghost icon indicates that the gene was not predicted by any of the literature-biased sources, namely, the gene is not involved (or is marginally involved) in our current understanding of autophagy; hence, it is potentially novel (ghost) for this phenotype of interest. The right part of the table depicts the percentile ranks generated by training TCGA-based GhostBuster against the “autophagy” list, and 8 lists representing sub-mechanisms thereof. The pink circle on the right side of each percentile displays the standardized GPT score, associating the gene with each sub-mechanism: essentially, scores of 5 or less indicate that the gene has not been strongly associated with that sub-mechanism, in the scientific literature. The 50 selected genes are sorted based on hierarchical clustering, performed on the sub-mechanism percentile ranks. The clustering tree is depicted on the right of the main table, and helps identify blocks of genes with a potentially similar biology. Autoph: autophagy; DnaDR: DNA damage response; Mitoch: autophagy of mitochondrion; MacAut: macroautophagy; MicAut: microautophagy; Reg+: positive regulation of autophagy; Reg–: negative regulation of autophagy.

**Supplementary Figure 7.**
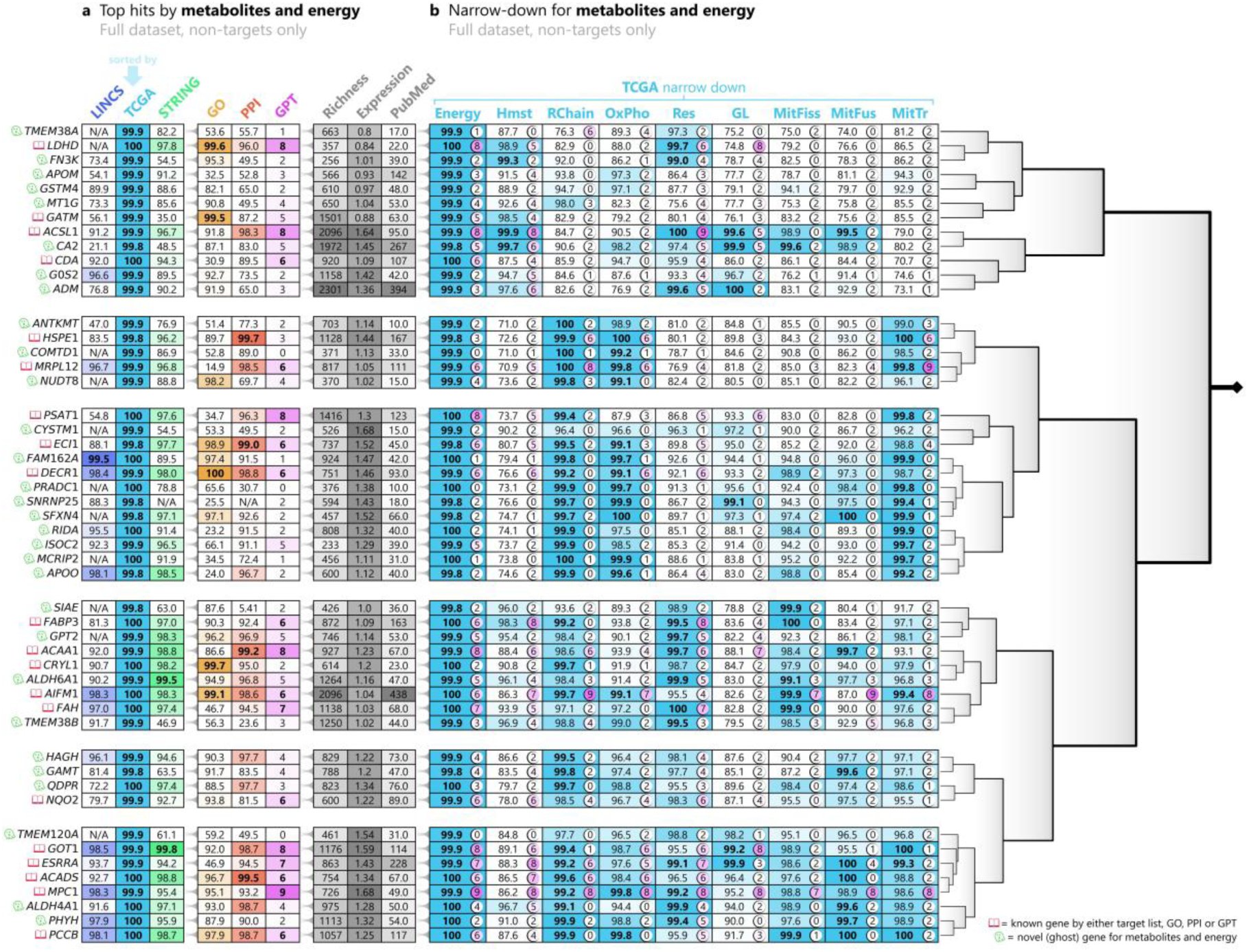
Tabular GhostBuster narrow-down plot across 7 sub-mechanisms of metabolism and energy. Tabular GhostBuster was trained to recognize 369 genes in the “generation of precursor metabolites and energy” list from Gene Ontology (GO), utilizing the TCGA channel as its training inputs (cyan arrow). At the end of the training, the top 50 non-target genes were selected, based on the prioritization list generated by such model in the full dataset, and are depicted here. The left part of the table showcases the percentile ranks generated by training Tabular GhostBuster against the “generation of precursor metabolites and energy” gene list, based on the LINCS (blue), TCGA (cyan), STRING (green) and GO (yellow) channels; next, GhostBuster’s auxiliary PPI scores (red), GPT scores (pink); and the richness, average expression and PubMed scores (grey). A crimson book icon on the left side of genes’ symbols is assigned for genes that are already known for playing a role in the phenotype of interest, either because they are target list members, top-rankers in the GO model (>99^th^ percentile), top rankers in the PPI score (>99^th^ percentile), or GPT-positive (GPT score > 5). Conversely, a green ghost icon indicates that the gene was not predicted by any of the literature-biased sources, namely, the gene is not involved (or is marginally involved) in our current understanding of metabolism and energy; hence, it is potentially novel (ghost) for this phenotype of interest. The right part of the table depicts the percentile ranks generated by training TCGA-based GhostBuster against the “generation of precursor metabolites and energy” list, and 8 lists representing sub-mechanisms thereof. The pink circle on the right side of each percentile displays the standardized GPT score, associating the gene with each sub-mechanism: essentially, scores of 5 or less indicate that the gene has not been strongly associated with that sub-mechanism, in the scientific literature. The 50 selected genes are sorted based on hierarchical clustering, performed on the sub-mechanism percentile ranks. The clustering tree is depicted on the right of the main table, and helps identify blocks of genes with a potentially similar biology. Energy: generation of precursor metabolites and energy; Hmst: energy homeostasis; RChain: mitochondrial respiratory chain complex; OxPho: oxidative phosphorylation; Res: energy reserve metabolic process; GL: glycolytic process; MitFiss: mitochondrial fission; MitFus: mitochondrial fusion; MTrnsl: mitochondrial translation.

**Supplementary Figure 8.**
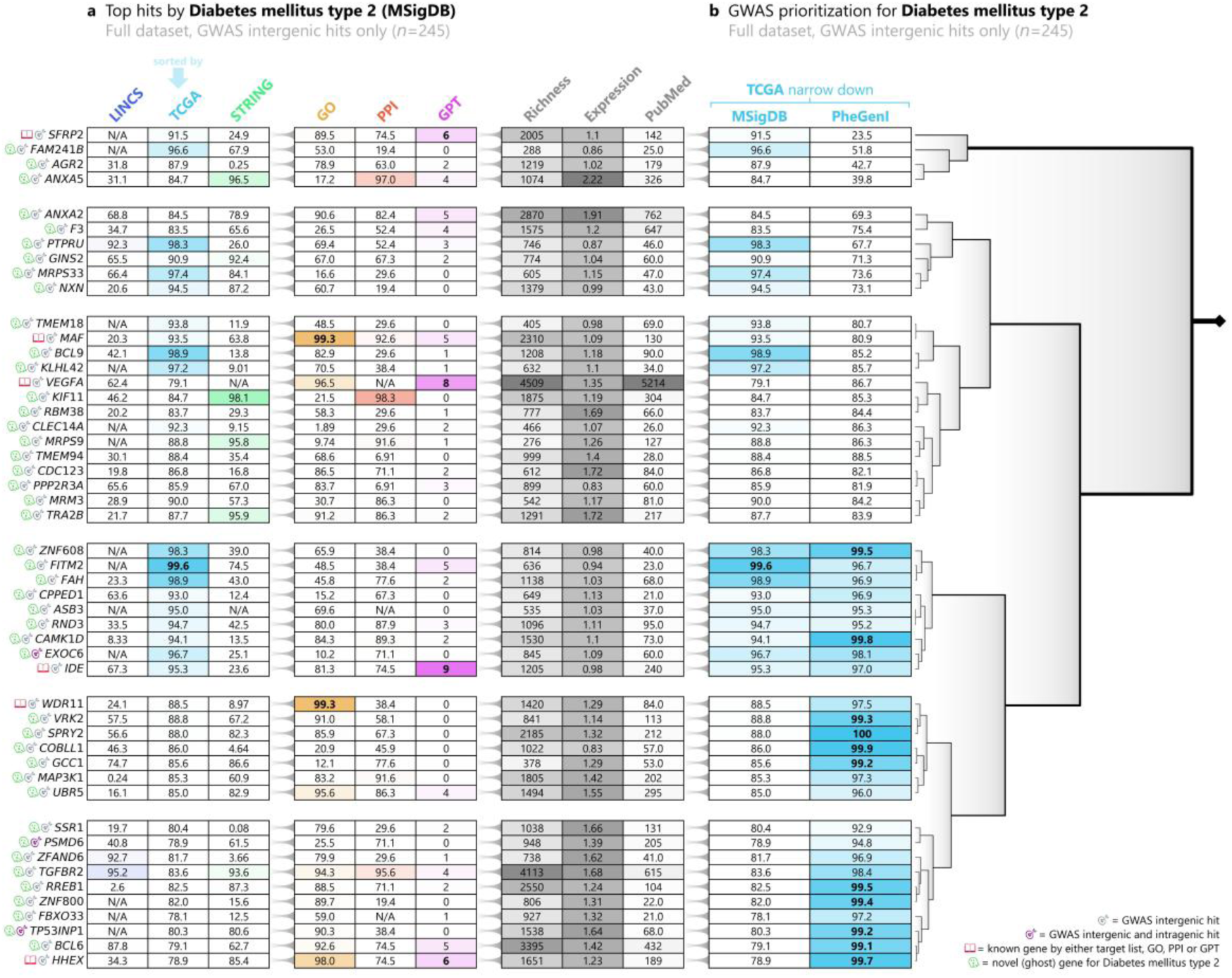
Tabular GhostBuster prioritization of intergenic GWAS gene hits in type 2 diabetes mellitus. Tabular GhostBuster was trained to recognize 223 genes in the “type 2 diabetes mellitus” (TD2) pathway, sourced from the Molecular Signatures Database (MSigDB); and 116 genes that were found as hits in genome-wide association studies (GWAS) on T2D, sourced from the Phenotype-Genotype Integrator (PheGenI). The latter included genes that hosted an intragenic or near-genic GWAS hits, as well as “intergenic gene hits”, that is, genes that were the closest in the primary sequence to an intergenic GWAS hit. Training was performed utilizing the TCGA channel as its training inputs (cyan arrow). At the end of the training, only the 177 intergenic gene hits were selected: of these, the top 50 genes were utilized for plotting, based on the prioritization list generated by the model trained on MSigDB in the full dataset, and are depicted here. Because intergenic gene hits are known for including a 50-75% share of false positives, the aim of this plot is to identify which ones are the most likely true positives, based on their similarity to the mechanistically involved genes (MSigDB), or to the GWAS hit list as a whole (PheGenI). The left part of the table showcases the percentile ranks generated by training Tabular GhostBuster against the TD2 MSigDB gene list, based on the LINCS (blue), TCGA (cyan), STRING (green) and GO (yellow) channels; next, GhostBuster’s auxiliary PPI scores (red), GPT scores (pink); and the richness, average expression and PubMed scores (grey). A grey target icon on the left side of genes’ symbols indicates that a gene is only a GWAS intergenic hit, while a purple target icon indicates that it is both an intergenic and an intragenic GWAS hit. A crimson book icon is assigned for genes that are already known for playing a role in the phenotype of interest, either because they are target list members, top-rankers in the GO model (>99^th^ percentile), top rankers in the PPI score (>99^th^ percentile), or GPT-positive (GPT score > 5). Conversely, a green ghost icon indicates that the gene was not predicted by any of the literature-biased sources, namely, the gene is not involved (or is marginally involved) in our current understanding of TD2; hence, it is potentially novel (ghost) for such phenotype. The right part of the table showcases the percentile ranks generated by training Tabular GhostBuster against the TD2 MSigDB list, and the corresponding PheGenI list. We define as “single hits” the genes that reach the 99^th^ percentile in the MSigDB channel or in the PheGenI, whereas “dual hits” are the genes that achieve the 99^th^ percentile in both channels at the same time. The 50 selected genes are sorted based on hierarchical clustering, performed on the MSigDB and PheGenI percentile ranks. The clustering tree is depicted on the right of the main table, and helps identify blocks of genes with a potentially similar biology.

**Supplementary Figure 9.**
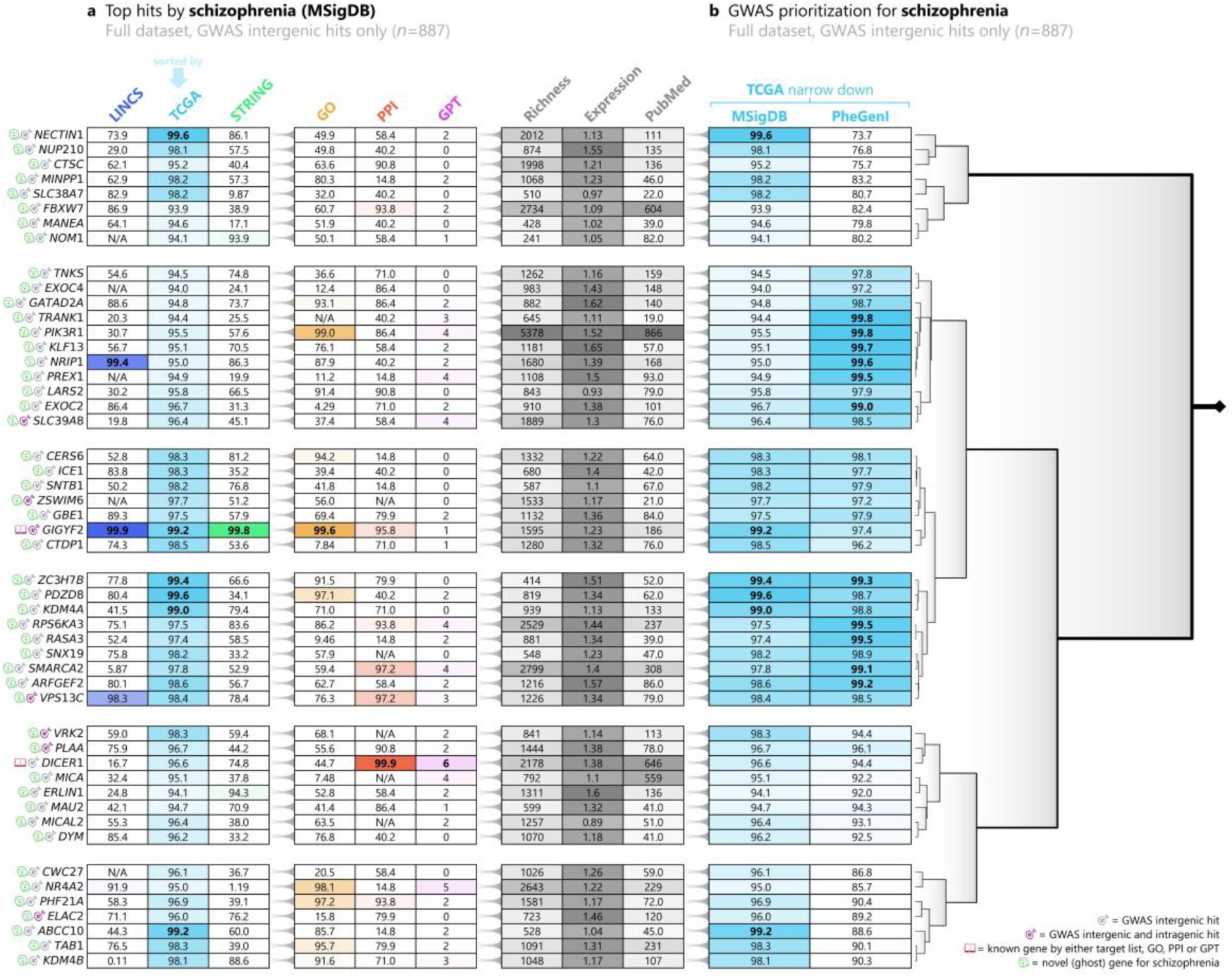
Tabular GhostBuster prioritization of intergenic GWAS gene hits in schizophrenia. Tabular GhostBuster was trained to recognize 74 genes in the schizophrenia pathway, sourced from the Molecular Signatures Database (MSigDB); and 212 genes that were found as hits in genome-wide association studies (GWAS) on schizophrenia, sourced from the Phenotype-Genotype Integrator (PheGenI). The latter included genes that hosted an intragenic or near-genic GWAS hits, as well as “intergenic gene hits”, that is, genes that were the closest in the primary sequence to an intergenic GWAS hit. Training was performed utilizing the TCGA channel as its training inputs (cyan arrow). At the end of the training, only the 624 intergenic gene hits were selected: of these, the top 50 genes were utilized for plotting, based on the prioritization list generated by the model trained on MSigDB in the full dataset, and are depicted here. Because intergenic gene hits are known for including a 50-75% share of false positives, the aim of this plot is to identify which ones are the most likely true positives, based on their similarity to the mechanistically involved genes (MSigDB), or to the GWAS hit list as a whole (PheGenI). The left part of the table showcases the percentile ranks generated by training Tabular GhostBuster against the schizophrenia MSigDB gene list, based on the LINCS (blue), TCGA (cyan), STRING (green) and GO (yellow) channels; next, GhostBuster’s auxiliary PPI scores (red), GPT scores (pink); and the richness, average expression and PubMed scores (grey). A grey target icon on the left side of genes’ symbols indicates that a gene is only a GWAS intergenic hit, while a purple target icon indicates that it is both an intergenic and an intragenic GWAS hit. A crimson book icon is assigned for genes that are already known for playing a role in the phenotype of interest, either because they are target list members, top-rankers in the GO model (>99^th^ percentile), top rankers in the PPI score (>99^th^ percentile), or GPT-positive (GPT score > 5). Conversely, a green ghost icon indicates that the gene was not predicted by any of the literature-biased sources, namely, the gene is not involved (or is marginally involved) in our current understanding of schizophrenia; hence, it is potentially novel (ghost) for such phenotype. The right part of the table showcases the percentile ranks generated by training Tabular GhostBuster against the schizophrenia MSigDB list, and the corresponding PheGenI list. We define as “single hits” the genes that reach the 99^th^ percentile in the MSigDB channel or in the PheGenI, whereas “dual hits” are the genes that achieve the 99^th^ percentile in both channels at the same time. The 50 selected genes are sorted based on hierarchical clustering, performed on the MSigDB and PheGenI percentile ranks. The clustering tree is depicted on the right of the main table, and helps identify blocks of genes with a potentially similar biology.

## Supplementary Tables

**Supplementary Table 1.**
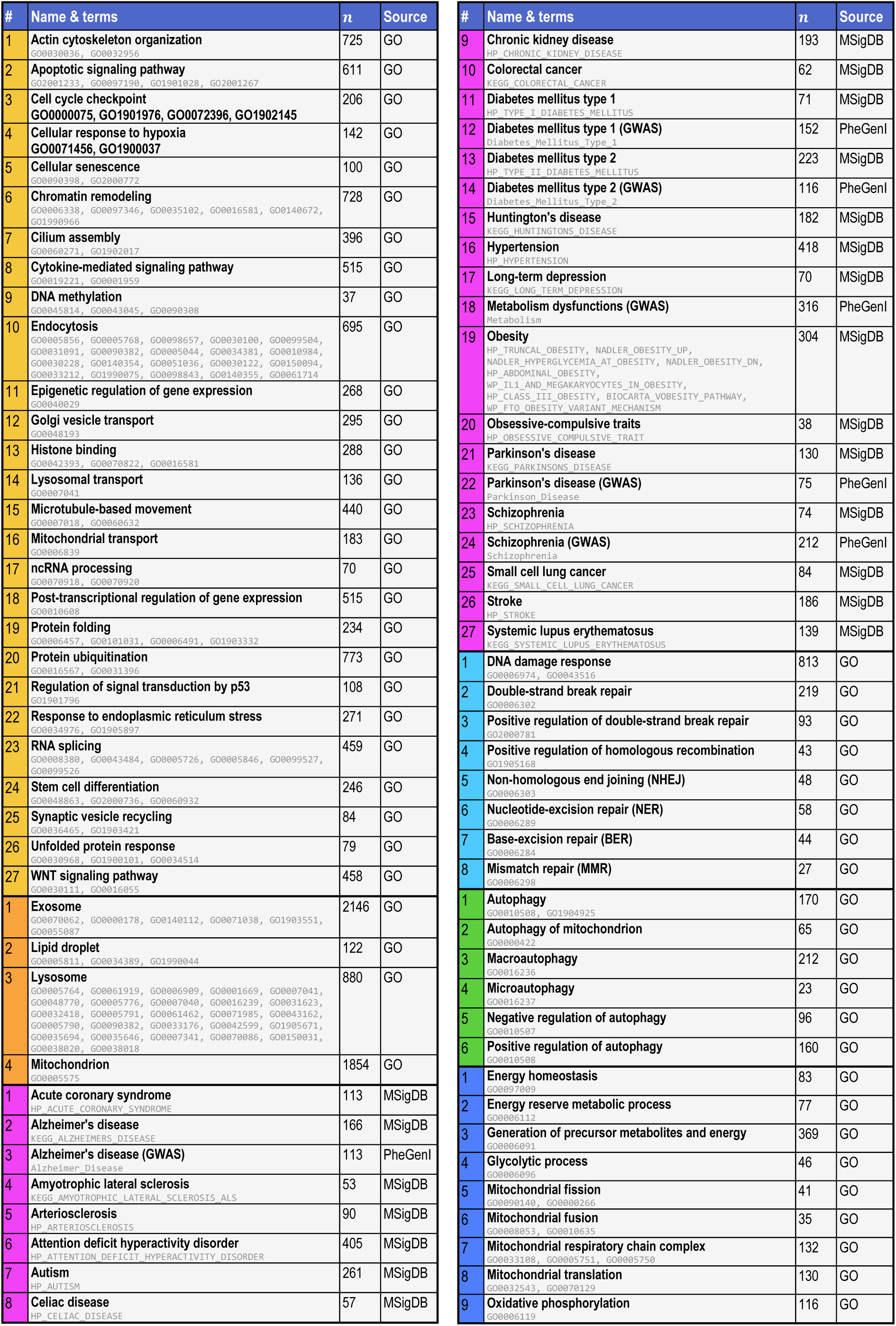
Target gene membership classes utilized to train Tabular GhostBuster. The table recapitulates the positive classes that were used as a target, to train the tabular version of GhostBuster. Column “#” encodes the class number within each of the 6 class domains (see Supplementary Figure 1): 27 physiological cell functions (yellow), 4 cell regions or organelles (orange), 27 diseases (pink), 8 DNA damage response narrow-down classes (cyan), 6 autophagy narrow-down classes (green) and 9 metabolism narrow-down classes (blue). The “Name & terms” column includes the class label that was utilized within GhostBuster, and lists all the original dataset’s terms that were pooled together to create such class, expressed in the original dataset’s nomenclature. The “*n*” column expresses the number of unique gene members in the final pooled class. The “Source” column expresses from which source the class was obtained: Gene Ontology (GO), the Molecular Signatures Database Hallmark (MSigDB) or the Phenotype-Genotype Integrator (PheGenI). In turn, the MSigDB classes originally came from either the Kyoto Encyclopedia of Genes and Genomes (KEGG), the Human Phenotype Ontology (HPO), Biocarta or MSigDB itself, as can be identified by their name prefixes. Of note, when the classes were deployed as positive targets in a machine learning task fed with a given dataset (e.g., TCGA), only the class members that appeared in the dataset’s namespace could be utilized. This is why, the class numerosity in the actual machine learning was ∼5-10% smaller than reported here.

**Supplementary Table 2.**
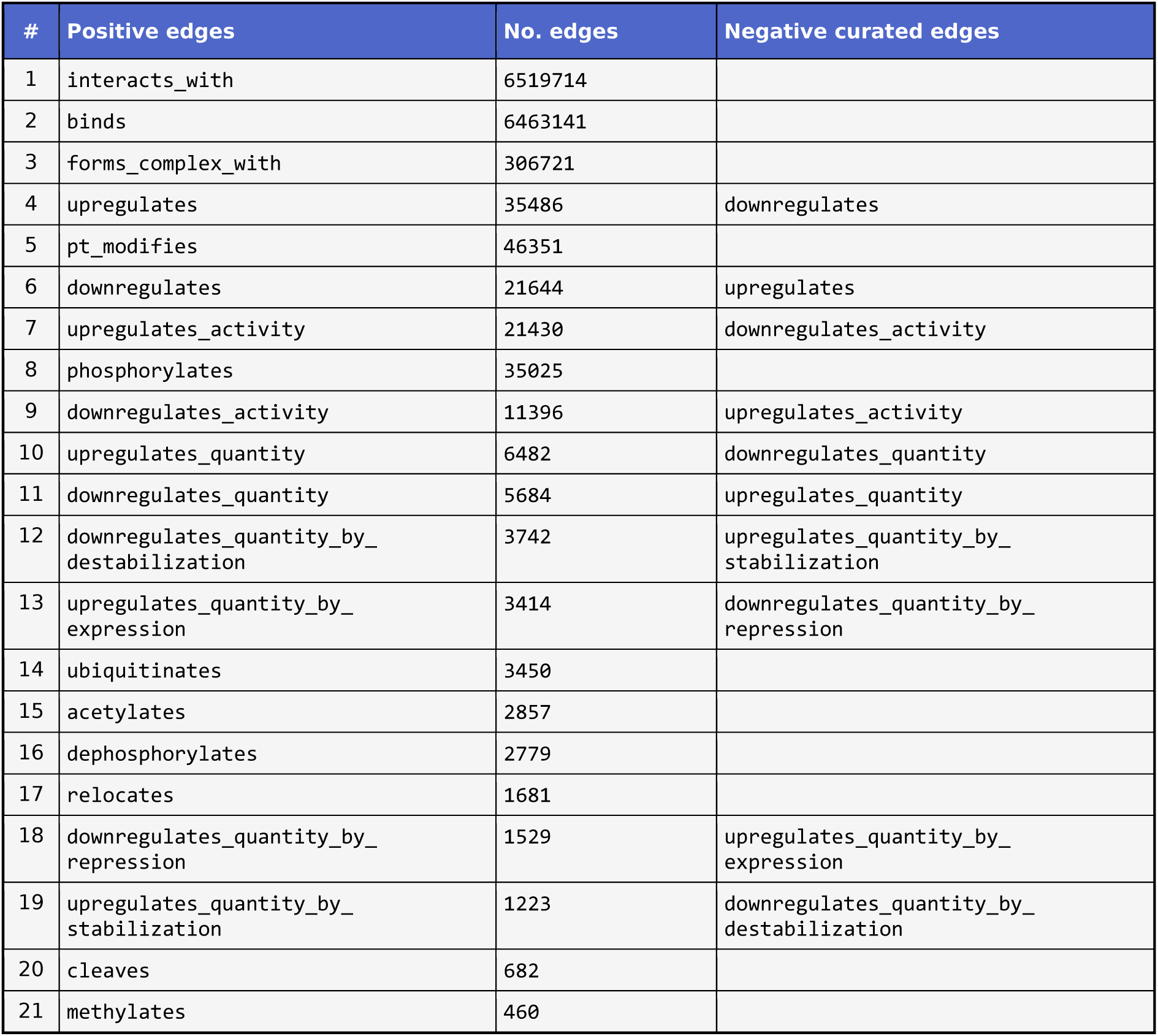
Target gene-gene relations utilized to train Graph GhostBuster. The table recapitulates the positive and curated negative gene-gene relations that were used as a target, to train the graph version of GhostBuster. All relations were sourced from the Biology Mega Graph, specifically, each relation type represents one level from such knowledge graph. Column “Positive edges” represents the level which was used as positive gene-gene edges in the edge prediction task, and column “No. edges” represent the numerosity thereof. Negative examples were generated randomly (random negatives). However, in the case where pairs of antonymic edge types were available, these were utilized as negatives (curated negatives), as an alternative to the randomly generated ones. The matched antonym for each positive edge type can be found in the “Negative curated edges” column. Of note, when the edges were deployed as positive (or negative) targets in a machine learning task fed with a given dataset (e.g., TCGA), only the gene nodes that appeared in the dataset’s namespace could be utilized. This is why, the edge numerosity in the actual machine learning was ∼5-10% smaller than reported here.

**Supplementary Table 3.**
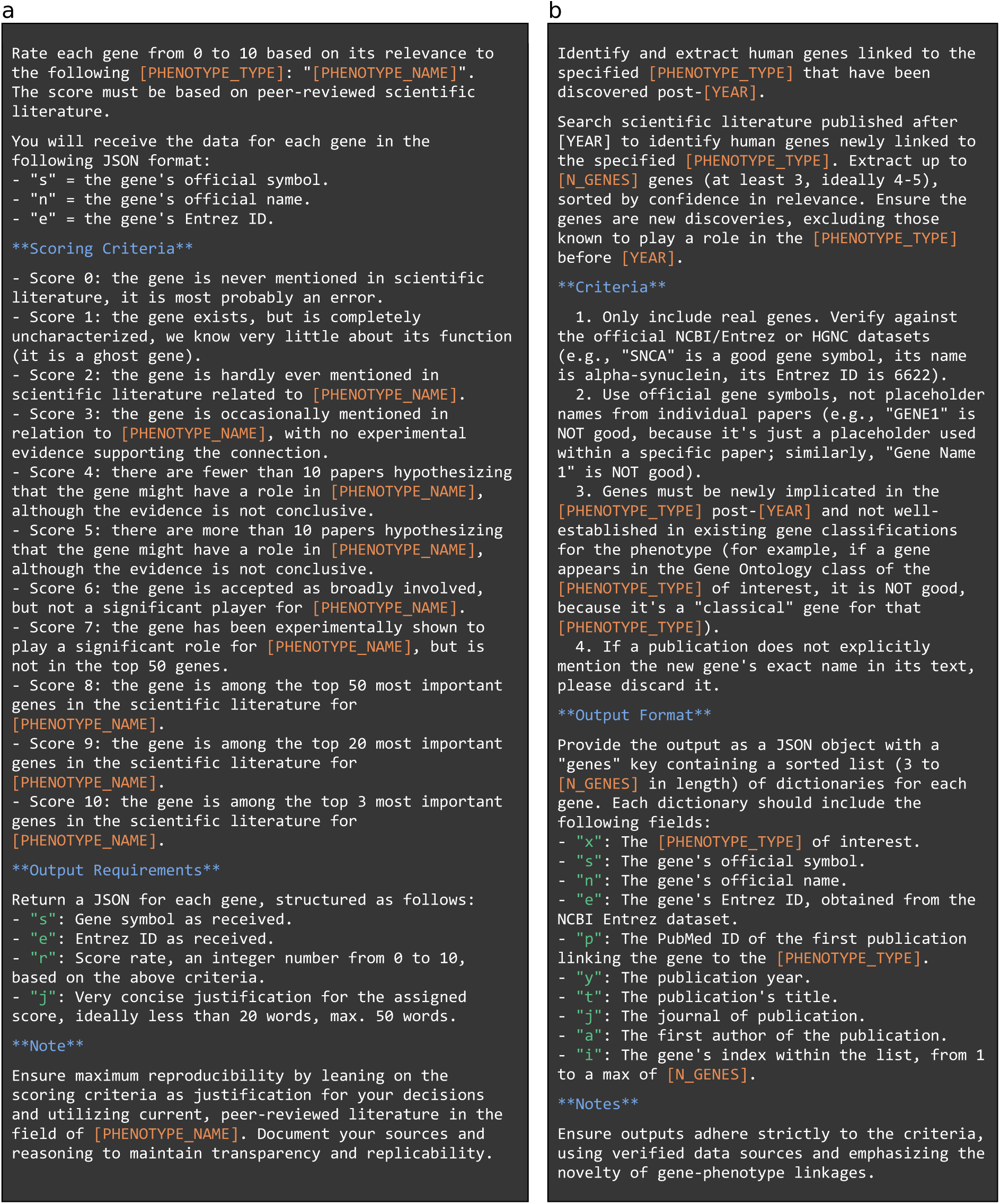
Natural language prompts submitted to the OpenAI API to generate the GPT control group of GhostBuster, and the list of recently discovered genes. (a) This prompt was utilized to create the Generative Pre-trained Transformer (GPT) control group of Tabular GhostBuster. It was submitted to the OpenAI application programming interface (API), selecting the GPT-4o mini model. The model was interrogated for every one of the 81 (of which, 75 unique) “phenotypes” of interest (i.e., a cell physiological function, a cell organelle or a disease) in Tabular GhostBuster, and for everyone of the 36,306 genes involved in Tabular GhostBuster, resulting in a total of 2,722,950 queries. Field [PHENOTYPE_NAME] was populated by the phenotype label, as reported in Supplementary Table 1. Field [PHENOTYPE_TYPE] was populated by the appropriate string among “cell function”, “cell organelle”, “cell region”, “disease” and “psychiatric disorder”. At each submission, the script was accompanied by a batch of 200 genes until the entire gene universe was explored. (b) This prompt was utilized to create the list of “recently discovered genes”, to validate Tabular GhostBuster’s ability to detect novel genes involved in a phenotype. It was submitted to the OpenAI application programming interface (API), selecting the GPT-4o (full) model. The model was interrogated for every one of the 81 (of which, 75 unique) “phenotypes” of interest, resulting in a total of 75 queries. Field [PHENOTYPE_TYPE] was populated as in (a). Field [N_GENES] was set to 30. Field [YEAR] was set to 2022. At each submission, the script was accompanied by a batch of 1 phenotype of interest (expressed by the phenotype label, as reported in Supplementary Table 1) until the entire phenotype universe was explored.

**Supplementary Table 4.** A list of 130 genes that were recently discovered (2022-2025) to be associated with Tabular GhostBuster’s target gene lists, according to GPT 4o mini. An automated script (see Supplementary Table 3) was created to interrogate OpenAI GPT-4o mini, to generate a list of genes that were recently (2022-2025) found to play an important role in the 81 biological processes or diseases (phenotypes) used to validate GhostBuster (see Supplementary Table 1). The target phenotype is shown in the left column, while the right columns list the genes highlighted by GPT-4o mini, in terms of gene symbol and Entrez ID.

| # | Target gene list | Recently discovered genes according to GPT-4o mini |  |  |  |
| --- | --- | --- | --- | --- | --- |
| 1 | Actin cytoskeleton organization | **ARHGAP36** (Entrez 158763) | **PLEKHG5** (Entrez 57449) | **TNS3** (Entrez 64759) |  |
| 2 | Acute coronary syndrome | **FAM177A1** (Entrez 283635) | **GPR15** (Entrez 2838) |  |  |
| 3 | Alzheimer's disease | **SORL1** (Entrez 6653) | **TREM2** (Entrez 54209) | **CLU** (Entrez 1191) |  |
| 4 | Amyotrophic lateral sclerosis | **NEK1** (Entrez 4750) | **TBK1** (Entrez 29110) | **CCNF** (Entrez 899) | **KIF5A** (Entrez 3798) |
| 5 | Apoptotic signaling pathway | **FAM210A** (Entrez 125228) | **TMEM123** (Entrez 114908) |  |  |
| 6 | Arteriosclerosis | **LINC01018** (Entrez 255167) | **FAM13A** (Entrez 10144) | **GPR15** (Entrez 2838) | **ADAMTS7** (Entrez 11173) |
| 7 | Attention deficit hyperactivity disorder | **GRM7** (Entrez 2917) | **SLC6A3** (Entrez 6531) |  |  |
| 8 | Autism | **GRIN2B** (Entrez 2904) |  |  |  |
| 9 | Autophagy | **TMEM41B** (Entrez 440026) | **VMP1** (Entrez 81671) | **ATG14** (Entrez 22863) |  |
| 10 | Celiac disease | **IL18RAP** (Entrez 8807) | **CCR9** (Entrez 10803) | **SH2B3** (Entrez 10019) |  |
| 11 | Cell cycle checkpoint | **CENPE** (Entrez 1062) | **KIF18B** (Entrez 146909) | **ANLN** (Entrez 54443) |  |
| 12 | Cellular response to hypoxia | **NDUFA4L2** (Entrez 56901) |  |  |  |
| 13 | Cellular senescence | **GPR68** (Entrez 8111) | **TMEM135** (Entrez 65084) | **SLC25A51** (Entrez 92014) | **ZNF827** (Entrez 152485) |
| 14 | Chromatin remodeling | **ZMYND8** (Entrez 23613) |  |  |  |
| 15 | Chronic kidney disease | **TMEM63C** (Entrez 57156) | **SLC47A2** (Entrez 146802) |  |  |
| 16 | Colorectal cancer | **GREM1** (Entrez 26585) | **RNF43** (Entrez 54894) | **NCOA4** (Entrez 8031) | **ZNRF3** (Entrez 84133) |
| 17 | Cytokine-mediated signaling pathway | **TNFSF15** (Entrez 9966) |  |  |  |
| 18 | DNA damage response | **NUSAP1** (Entrez 51203) | **CENPF** (Entrez 1063) | **SPDL1** (Entrez 54908) | **KIF18A** (Entrez 81930) |
| 19 | DNA methylation | **ZBTB38** (Entrez 253461) | **TET3** (Entrez 200424) | **UHRF2** (Entrez 115426) |  |
| 20 | Endocytosis | **TMEM106B** (Entrez 54664) | **GPRC5B** (Entrez 51704) | **ANKRD13C** (Entrez 81573) |  |
| 21 | Epigenetic regulation of gene expression | **ZC3H4** (Entrez 23211) | **KDM8** (Entrez 79831) | **SETD9** (Entrez 133383) |  |
| 22 | Exosome | **NDUFAF5** (Entrez 79133) | **TMEM106B** (Entrez 54664) | **SLC25A46** (Entrez 91137) |  |
| 23 | Golgi vesicle transport | **TMEM165** (Entrez 55858) | **GOLGA8A** (Entrez 23015) | **CCDC186** (Entrez 55088) | **TMEM87B** (Entrez 84910) |
| 24 | Histone binding | **BRWD1** (Entrez 54014) | **SP140** (Entrez 11262) |  |  |
| 25 | Huntington's disease | **BACH1** (Entrez 571) | **ATXN1** (Entrez 6310) |  |  |
| 26 | Hypertension | **GPRC5B** (Entrez 51704) | **SLC25A45** (Entrez 283130) | **TMEM63C** (Entrez 57156) | **ZNF804A** (Entrez 91752) |
| 27 | Long-term depression | **GPR158** (Entrez 57512) | **SLC6A15** (Entrez 55117) | **TMEM161B** (Entrez 153396) |  |
| 28 | Lysosomal transport | **SNX14** (Entrez 57231) |  |  |  |
| 29 | Microtubule-based movement | **DCTN4** (Entrez 51164) | **SPAG5** (Entrez 10615) |  |  |
| 30 | Mitochondrial respiratory chain complex | **COA7** (Entrez 65260) |  |  |  |
| 31 | Mitochondrial translation | **MTFMT** (Entrez 123263) | **GTPBP3** (Entrez 84705) | **MTO1** (Entrez 25821) |  |
| 32 | Mitochondrial transport | **MIEF2** (Entrez 125170) |  |  |  |
| 33 | ncRNA processing | **ZCCHC7** (Entrez 84186) | **RBM33** (Entrez 155435) | **LARP7** (Entrez 51574) | **ZNF827** (Entrez 152485) |
| 34 | Obesity | **GPR151** (Entrez 134391) | **KLF14** (Entrez 136259) | **ADCY3** (Entrez 109) |  |
| 35 | Obsessive-compulsive traits | **GRIN2B** (Entrez 2904) | **SLC1A1** (Entrez 6505) | **DRD4** (Entrez 1815) |  |
| 36 | Parkinson's disease | **GBA1** (Entrez 2629) | **TMEM175** (Entrez 84286) |  |  |
| 37 | Post-transcriptional regulation of gene expr. | **PTBP3** (Entrez 9991) |  |  |  |
| 38 | Protein folding | **NUB1** (Entrez 51667) | **HSPA12A** (Entrez 259217) | **BAG6** (Entrez 7917) |  |
| 39 | Protein ubiquitination | **TRIM67** (Entrez 440730) |  |  |  |
| 40 | Regulation of signal transduction by p53 | **ZMAT3** (Entrez 64393) | **RPS27L** (Entrez 51065) | **TP53INP1** (Entrez 94241) | **CDKN1A** (Entrez 1026) |
| 41 | Response to endoplasmic reticulum stress | **ZNF827** (Entrez 152485) |  |  |  |
| 42 | Schizophrenia | **GRIA3** (Entrez 2892) | **CACNA1I** (Entrez 8911) | **NRXN1** (Entrez 9378) |  |
| 43 | Small cell lung cancer | **GPRC5A** (Entrez 9052) | **KIF5B** (Entrez 3799) | **RAB3B** (Entrez 5865) |  |
| 44 | Stem cell differentiation | **ZBTB46** (Entrez 140685) | **KLF17** (Entrez 128209) | **PRDM16** (Entrez 63976) | **ZNF521** (Entrez 25925) |
| 45 | Stroke | **GPR17** (Entrez 2840) | **NLRP3** (Entrez 114548) | **TRPM4** (Entrez 54795) |  |
| 46 | Synaptic vesicle recycling | **RPH3A** (Entrez 22895) | **SYT17** (Entrez 51760) | **VAMP7** (Entrez 6845) | **SNAP47** (Entrez 116841) |
| 47 | Systemic lupus erythematosus | **NFKB1** (Entrez 4790) | **IRF5** (Entrez 3663) | **TNFAIP3** (Entrez 7128) |  |
| 48 | Type I diabetes mellitus | **PTPN22** (Entrez 26191) |  |  |  |
| 49 | Type II diabetes mellitus | **SLC16A11** (Entrez 162515) | **ZBED3** (Entrez 84327) | **TMEM163** (Entrez 81615) |  |
| 50 | Unfolded protein response | **NUPR1** (Entrez 26471) | **DNAJC3** (Entrez 5611) |  |  |

**Supplementary Table 5.**
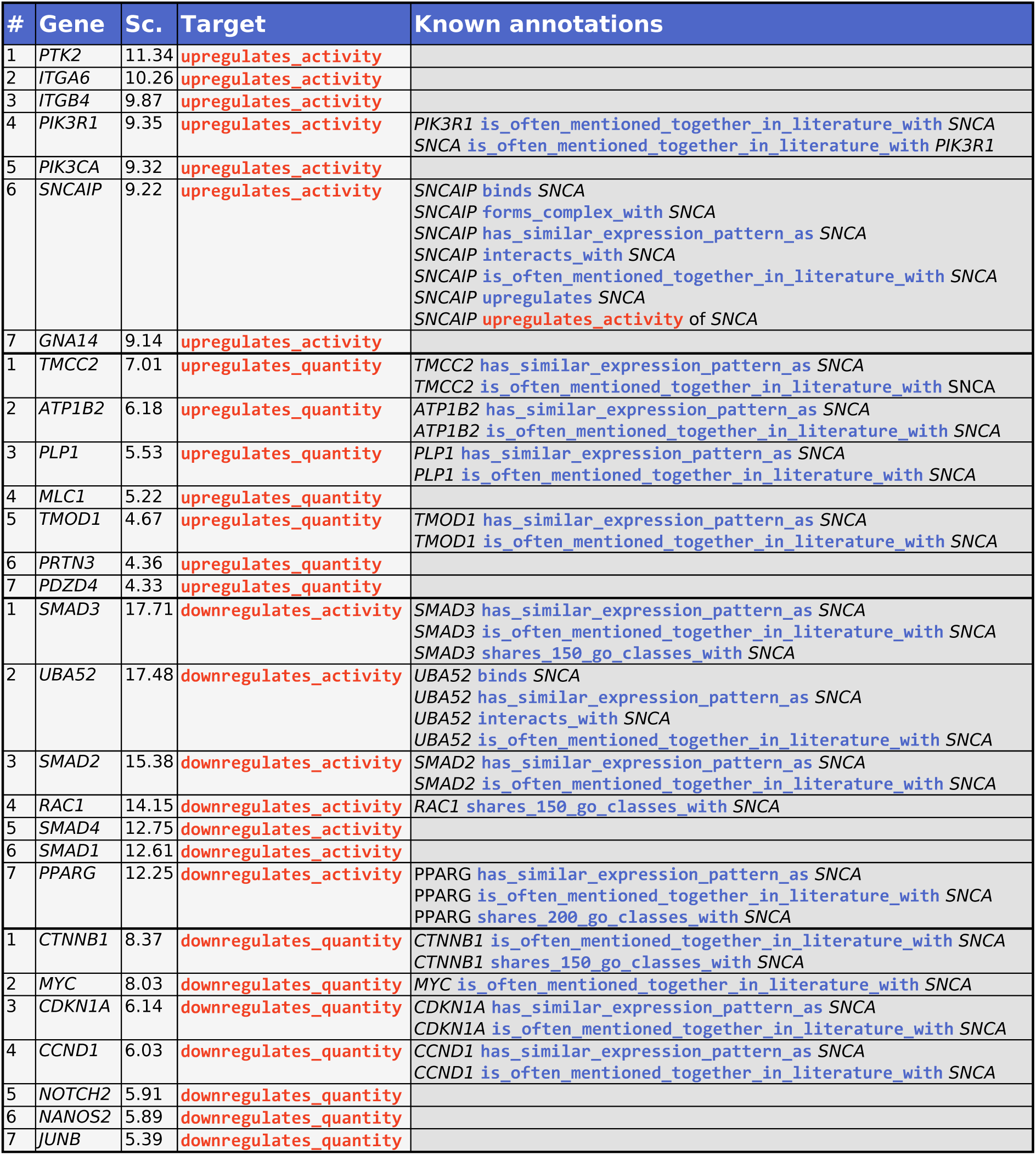
Graph GhostBuster results for alpha-synuclein, showcasing the top 7 predicted edges across 4 gene-gene interaction types. The table lists the predicted interactors of alpha-synuclein (αS protein, *SNCA* gene), according to Graph GhostBuster; specifically, the top 7 interactors are included from each of the following interaction modalities: `upregulates_activity`, `upregulates_quantity`, `downregulates_activity`, `downregulates_quantity`. A graph convolutional neural network (GCN) was trained independently for each of these interaction modalities, in an edge prediction task, where positive edges were sourced from the Biology Mega Graph, and node features were sourced from 24,196 RNA-sequencing-based transcriptomic vectors from The Cancer Genome Atlas (TCGA). The “#” column represents the interactor gene’s rank, based on the GCN’s logit, i.e., the raw dot-product between the best learned latent representation of the corresponding nodes. The “Gene” column expresses the interactor gene’s standardized gene symbol. The “Score” column expresses the logit value (before passing it through a sigmoid activation function). The “Known annotations” column lists the known interactions between the helper gene and alpha-synuclein, and their modality, as they appear in the Biology Mega Graph. Of note, because a GCN was utilized, predictions are computed by dot product, which is a commutative operation; as a result, the output is non-directional, i.e., it cannot distinguish between proteins that (say) upregulate αS, or are upregulated by αS. Histone proteins, RNA polymerase member proteins and ribosome proteins were filtered out from this table, as they represent a “trivial” interactor for the sake of upregulation.

**Supplementary Table 6.** Graph GhostBuster results for TDP-43, showcasing the top 7 predicted edges across 4 gene-gene interaction types. The table lists the predicted interactors of TDP-43 (*TARDBP* gene), according to Graph GhostBuster; specifically, the top 7 interactors are included from each of the following interaction modalities: `upregulates_activity`, `upregulates_quantity`, `downregulates_activity`, `downregulates_quantity`. A graph convolutional neural network (GCN) was trained independently for each of these interaction modalities, in an edge prediction task, where positive edges were sourced from the Biology Mega Graph, and node features were sourced from 24,196 RNA-sequencing-based transcriptomic vectors from The Cancer Genome Atlas (TCGA). The “#” column represents the interactor gene’s rank, based on the GCN’s logit, i.e., the raw dot-product between the best learned latent representation of the corresponding nodes. The “Gene” column expresses the interactor gene’s standardized gene symbol. The “Score” column expresses the logit value (before passing it through a sigmoid activation function). The “Known annotations” column lists the known interactions between the helper gene and alpha-synuclein, and their modality, as they appear in the Biology Mega Graph. Of note, because a GCN was utilized, predictions are computed by dot product, which is a commutative operation; as a result, the output is non-directional, i.e., it cannot distinguish between proteins that (say) upregulate αS, or are upregulated by αS. Histone proteins, RNA polymerase member proteins and ribosome proteins were filtered out from this table, as they represent a “trivial” interactor for the sake of upregulation.

| # | Gene | Sc. | Target | Known annotations |
| --- | --- | --- | --- | --- |
| 1 | EIF2S1 | 5.80 | upregulates_activity | EIF2S1 binds TARDBP<br>EIF2S1 has_similar_expression_pattern_as TARDBP<br>EIF2S1 interacts_with TARDBP<br>EIF2S1 is often mentioned together in literature with TARDBP |
| 2 | EIF2S2 | 5.64 | upregulates_activity | EIF2S2 has_similar_expression_pattern_as TARDBP<br>EIF2S2 is often mentioned together in literature with TARDBP |
| 3 | EIF2S3 | 5.52 | upregulates_activity | EIF2S3 binds TARDBP<br>EIF2S3 has_similar_expression_pattern_as TARDBP<br>EIF2S3 interacts_with TARDBP<br>EIF2S3 is often mentioned together in literature with TARDBP |
| 4 | RBX1 | 5.51 | upregulates_activity | RBX1 binds TARDBP<br>RBX1 has_similar_expression_pattern_as TARDBP<br>RBX1 interacts_with TARDBP<br>RBX1 is often mentioned together in literature with TARDBP |
| 5 | RPS10 | 5.19 | upregulates_activity | RPS10 binds TARDBP<br>RPS10 has_similar_expression_pattern_as TARDBP<br>RPS10 interacts_with TARDBP<br>RPS10 is often mentioned together in literature with TARDBP |
| 6 | RPS5 | 5.08 | upregulates_activity | RPS5 binds TARDBP<br>RPS5 has_similar_expression_pattern_as TARDBP<br>RPS5 interacts_with TARDBP<br>RPS5 is often mentioned together in literature with TARDBP |
| 7 | CRCP | 4.99 | upregulates_activity |  |
| 1 | CDKN2B | 15.04 | upregulates_quantity |  |
| 2 | XPOT | 15.00 | upregulates_quantity | XPOT binds TARDBP<br>XPOT has_similar_expression_pattern_as TARDBP<br>XPOT interacts_with TARDBP<br>XPOT is often mentioned together in literature with TARDBP |
| 3 | CDKN2A | 13.35 | upregulates_quantity | CDKN2A binds TARDBP<br>CDKN2A has_similar_expression_pattern_as TARDBP<br>CDKN2A interacts_with TARDBP<br>CDKN2A is often mentioned together in literature with TARDBP |
| 4 | MYC | 12.60 | upregulates_quantity | MYC binds TARDBP<br>MYC has_similar_expression_pattern_as TARDBP<br>MYC interacts_with TARDBP<br>MYC is often mentioned together in literature with TARDBP |
| 5 | CDKN1A | 10.38 | upregulates_quantity |  |
| 6 | HES1 | 9.99 | upregulates_quantity |  |
| 7 | TP53 | 9.94 | upregulates_quantity | TP53 is often mentioned together in literature with TARDBP |
| 1 | UBA52 | 7.85 | downregulates_activity |  |
| 2 | SMAD3 | 7.10 | downregulates_activity |  |
| 3 | SMAD2 | 6.07 | downregulates_activity | SMAD2 has_similar_expression_pattern_as TARDBP<br>SMAD2 is often mentioned together in literature with TARDBP |
| 4 | RAC1 | 5.08 | downregulates_activity |  |
| 5 | LOC102724334 | 4.98 | downregulates_activity |  |
| 6 | SMAD4 | 4.85 | downregulates_activity |  |
| 7 | PPARG | 4.78 | downregulates_activity | PPARG binds TARDBP<br>PPARG interacts_with TARDBP<br>PPARG is often mentioned together in literature with TARDBP |
| 1 | CTNNB1 | 17.39 | downregulates_quantity | CTNNB1 binds TARDBP<br>CTNNB1 has_similar_expression_pattern_as TARDBP<br>CTNNB1 interacts_with TARDBP<br>CTNNB1 is often mentioned together in literature with TARDBP |
| 2 | MYC | 17.10 | downregulates_quantity | MYC binds TARDBP<br>MYC has_similar_expression_pattern_as TARDBP<br>MYC interacts_with TARDBP<br>MYC is often mentioned together in literature with TARDBP |
| 3 | NOTCH2 | 13.02 | downregulates_quantity |  |
| 4 | CDKN1A | 12.71 | downregulates_quantity |  |
| 5 | CCND1 | 12.50 | downregulates_quantity | CCND1 binds TARDBP<br>CCND1 interacts_with TARDBP<br>CCND1 is often mentioned together in literature with TARDBP |
| 6 | NANOS2 | 12.35 | downregulates_quantity |  |
| 7 | PUM2 | 11.57 | downregulates_quantity | PUM2 binds TARDBP<br>PUM2 has_similar_expression_pattern_as TARDBP<br>PUM2 interacts_with TARDBP<br>PUM2 is often mentioned together in literature with TARDBP |

